# A transcriptional code controlling fluid shear stress-induced gene expression

**DOI:** 10.64898/2026.02.04.703710

**Authors:** Lucija Fleisinger, Susann Bruche, Hyewon Lim, Anna Rataj, Helena Rodriguez-Caro, Amaury Genovese, Vinesh Vinayachandran, Svanhild Nornes, Dorota Szumska, Dhruv S Gupta, Indrika Ratnayaka, Kira Chouliaras, Marek Giers, Simon J Conway, Alice Neal, Sophie Payne, Martin A Schwartz, Mukesh K Jain, Brian G Coon, Sarah De Val

**Author notes:** contributed equally (Lucija Fleisinger, Susann Bruche).

## Abstract

The formation and health of the vascular system is dependent on fluid shear stress (FSS), a hemodynamic force exerted onto endothelium by flowing blood. FSS strongly induces the endothelial expression of Krüppel-like factor 2 (KLF2), an atheroprotective TF essential for vascular development and homeostasis. Despite its early and crucial role in the cascade of cardiovascular events triggered by FSS, the transcriptional mechanisms by which FSS regulates *KLF2* expression remain unclear, although they are known to involve the widely expressed MEF2 proteins. Here, we identified and characterized two FSS-dependent enhancers for *KLF2* which collectively recapitulate endogenous endothelial *KLF2* expression, and determined the TFs contributing to their regulation. This analysis identified an essential and precisely spaced MEF2-TBP double motif also shared by the FSS-sensitive *KLF2* promoter. MEF-TBP double motifs are extremely rare across the genome but were also found within regulatory elements of three other FSS-induced *KLF* genes, including *KLF4*. Although normally part of the basal transcriptional machinery, TBP specifically bound all *KLF* elements at the MEF-TBP double motifs in a FSS-dependent manner. Collectively, this work demonstrates a specific and targetable requirement for combined MEF2-TBP binding during FSS-induced gene activation.

**Significance statement:** Blood flow induces a force known as fluid shear stress (FSS) which is required for vascular development and for the health of the mature arterial system. One of the first endothelial responses to FSS is the induction of Krüppel-like transcription factors (KLFs). However, the mechanisms by which FSS activates *KLF* gene expression are incompletely understood. In this paper, we characterized all regulatory elements involved in driving FSS-induced expression of *KLF2*. This identified an essential MEF2-TBP double motif that was extremely rare across the genome, yet found within regulatory elements for multiple FSS-responsive KLF genes including *KLF2* and *KLF4*. This MEF2– and TBP-bound motif therefore enables blood flow to specifically activate the cascade of cardiovascular responses necessary for atheroprotective gene expression.

## Introduction

The mammalian blood vascular system is a hierarchically organised arterial-capillary-venous network transporting blood, metabolites, and waste products. Endothelial cells (EC) form the inner lining of the blood vascular system, and show significant heterogeneity downstream of cell-intrinsic, local environmental and haemodynamic stimuli (1, 2). Blood flow-induced fluid shear stress (FSS) is required for the proper differentiation and maintenance of arterial endothelium (3–5), for angiogenesis and for valve formation (6–8). FSS is also essential for the homeostasis and health of the mature artery, ensuring that the endothelium acts as an anti-inflammatory, anti-thrombotic barrier between blood and underlying vessel or tissue (9). Arterial regions that are not exposed to strong, unidirectional FSS (e.g. branch points, sites of high curvature) are prone to vascular dysfunction and to developing atherosclerotic lesions (5, 9).

Krüppel-like factors (KLFs) are a family of DNA-binding transcription factors (TFs). Both KLF2 and KLF4 play crucial roles in ECs, although they are also expressed in other cell types (10). In the developing vasculature, they influence arteriovenous formation, heart valve development and angiogenesis, while in mature vascular networks they are critical for vascular stability (10). In humans, alterations to KLF2 and KLF4 levels are linked to cardiovascular diseases that affect millions, including atherosclerosis, thrombosis, lymphedema, and cerebral cavernous malformations (CCMs) (9, 11, 12). Genetic ablation of *Klf2* or *Klf4* in mouse endothelium results in death during embryonic development or shortly after birth, while hemizygous deletion alters blood-brain barrier function and increases atherosclerosis and thrombosis (10, 12, 13). Compound deletion of *Klf2* with *Klf4* in adult endothelium results in rapid death alongside compromised vascular integrity and dysregulated coagulation (13).

*KLF2* and *KLF4* expression in ECs is specifically upregulated (at both mRNA and protein levels) by FSS, which in early development is strongest in the cardiac outflow tract, endocardial cushions and valves (14). KLF2 and KLF4 are also expressed at higher levels in arterial and capillary ECs, where FSS is greater (14). These patterns are recapitulated *in vitro*, with robust induction of *KLF2/4* mRNA and protein in cultured ECs when exposed to FSS (15, 16).

The MEKK3-MEK5-ERK5 kinase cascade is critical for FSS-induced expression of *KLF2* (17–19). In the current canonical model, ERK5 activates MEF2 TFs via phosphorylation, leading to increased *KLF2* transcription. Supporting this, genetic perturbation of *Erk5* leads to reduced *Klf2* mRNA and embryonic death, while dominant-negative MEK5 and MEF2 mutants prevent FSS-mediated induction of *KLF2* expression (17, 18). This model is assumed to extend to *KLF4*, with EC-specific compound deletion of *Mef2a; Mef2c; Mef2d* in adults strongly decreasing *Klf2/4* expression and recapitulating many features associated with *Klf2/4* deletion (13, 20). While this confirms a link between MEF2 factors and *KLF2/4*, this model does not explain the specificity of FSS-induced EC expression of *KLF2/4*. MEF2 factors are widely expressed and play key regulatory roles in many cell types, and their transcriptional activity is affected by multiple signalling pathways in addition to MAP kinases (e.g. HDACs, calcium signalling, VEGFA-VEGFR2 (19, 21, 22)). Additionally, MEF2 factors are broadly expressed throughout the endothelium, and regulate EC genes and signalling pathways unregulated and unaffected by FSS. In particular, MEF2 factors are key activators of angiogenic sprouting through direct binding and transcriptional activation of *Dll4*, *Hlx* and multiple *Ets* factors downstream of VEGFA-VEGFR2 (21). It is therefore highly likely that additional regulators work alongside MEF2 to specifically regulate *KLF2/4* expression and subsequently maintain endothelial homeostasis.

In this paper, we identify and characterize two enhancers for *KLF2,* demonstrating that both require FSS for activation. By investigating the TF motifs shared by both *KLF2* enhancers and the FSS-sensitive *KLF2* promoter, we uncovered an unexpected role for TBP alongside MEF2 in FSS-induced *KLF2* expression. This occurs via a distinct MEF2-TBP double motif, with FSS stimulating binding of TBP to these regulatory elements. The MEF2-TBP double motif was not found within other MEF2-target enhancers and promoters that lack FSS responsive, while genome-wide analysis demonstrates these motifs are restricted to regulatory elements for FSS-induced *KLF* genes. Our findings therefore describe a distinct transcriptional code that directly links blood flow to *KLF*-dependent endothelial homeostasis.

## Results

### Identification of two upstream enhancers for *KLF2*

Analysis of FSS induction of *KLF2* expression has previously focused on the proximal promoter region, which shows FSS-responsiveness and ERK5/MEK5/MEF2-dependence *in vitro*. To elucidate its role *in vivo*, we cloned the region between –358bp to +17bp relative to the transcriptional start site upstream of a promoterless β-galactosidase (*LacZ*) reporter plasmid (e.g. (23, 24)). This promoter element (termed *KLF2pr-0.4*) contained all key motifs identified in previous *in vitro* studies, including the MEF2 motif essential for the ERK5/MEK5 and FSS-responsiveness of this promoter (18, 19, 25–27). The activity of the *KLF2pr-0.4:LacZ* transgene was tested in transient transgenic mouse embryos at embryonic day (E)12.5, a timepoint at which *KLF2* mRNA is robustly expressed in some ECs (Figure 1A-D). However, only two of seven transgenic embryos showed any detectable expression, in both cases very faint (Figure 1C-D and S1A), suggesting that the promoter alone is not sufficient to recapitulate expression *in vivo*.

**Figure 1.**
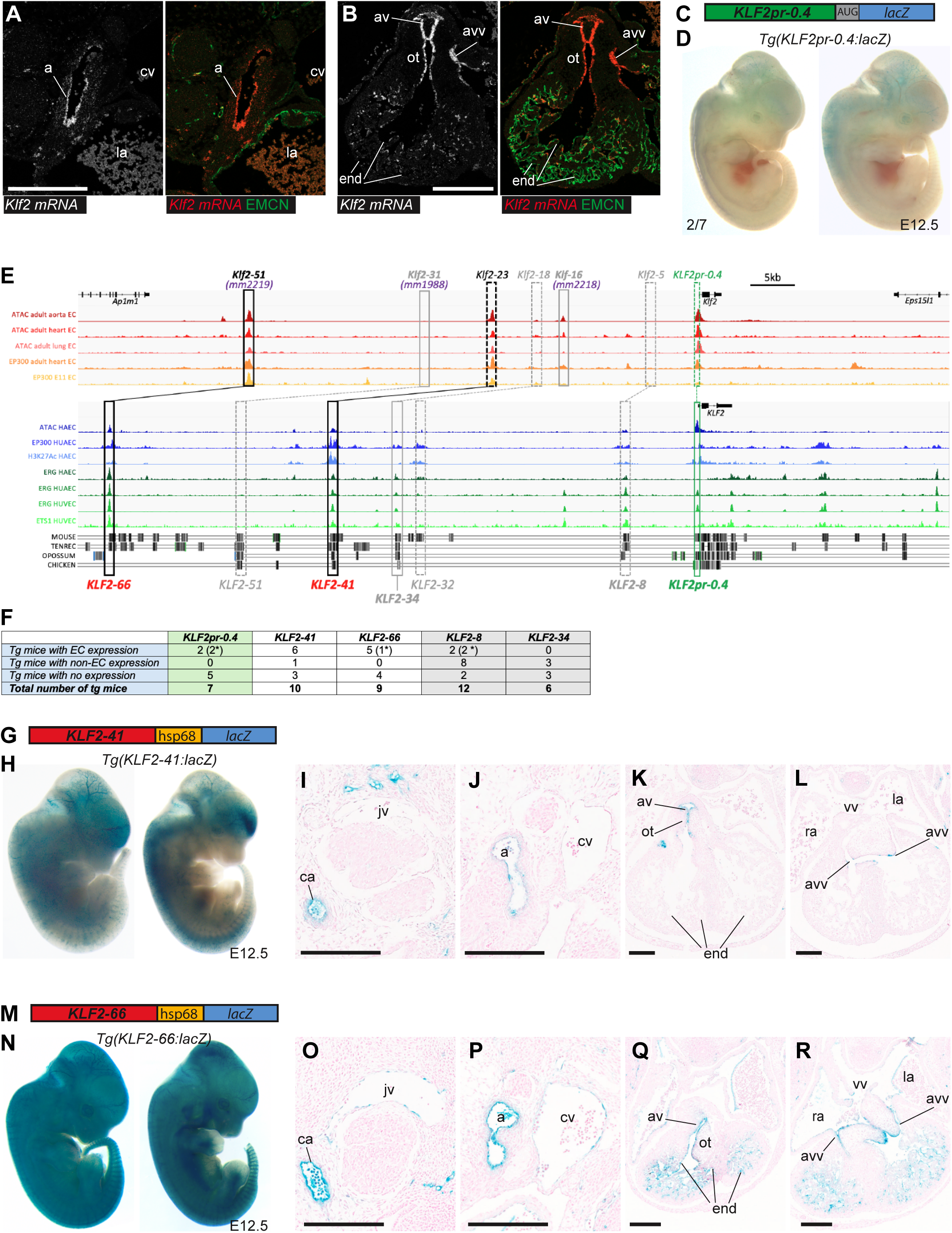
The *KLF2-41* and *KLF2-66* enhancers recapitulate endogenous *KLF2* expression patterns in E12.5 F0 mouse embryos. **A-B** Endogenous *Klf2* expression in ECs of arteries (**A**), endocardium and heart valves (**B**), assessed by RNA-scope in transverse sections through E12.5 embryos compared with the EC marker Emcn. **C-D** The *KLF2pr-0.4:lacZ* transgene (**C**) directs only very weak expression in a minority of E12.5 F0 transgenic mice (**D**), see also Figure S1A. **E** Analysis of enhancer marks in mouse (upper) and human (lower) ECs around the *Klf2/KLF2* locus. ATAC-seq (ATAC) adult aorta EC from (75), adult heart and lung EC ATAC-seq and EP300 binding from(76), EP300 binding in Tie2Cre+ve cells in E11.5 embryos (E11 EC) from (77); ATAC-seq HAEC (human aortic ECs) and H3K27Ac HAEC from (78); EP300 HUAEC (human umbilical arterial ECs) from (79). ERG and ETS ChIP-seq data: ERG HAEC from (78); ERG HUAEC and HUVEC (79); and ETS in HUVEC from (79). Black vertical lines indicate depth of conservation between human species with mouse, tenrec, opossum and chicken sequence. Black boxes mark validated enhancers (dashed lines indicate orthologous regions), grey boxes mark regions tested as enhancers (either mouse or human version) but not active, green box indicates promoter. **F** Table indicating expression patterns seen in all F0 transgenic embryos. * denotes weak expression only. **G-R** The *KLF2-41:hsp68:lacZ* transgene (**G**) and *KLF2-66:hsp68:lacZ* transgene (**M**) directs EC-specific expression of the *lacZ* reporter gene in F0 transgenic mice (**H-L, N-R**) in a pattern similar to endogenous *Klf2*. **H-L** Two representative wholemount E12 F0 transgenic embryos (**H**) and representative transverse sections from embryos expressing *KLF2-41:hsp68:lacZ* (**I-L**). **N-R** Two representative wholemount E12 F0 transgenic embryos (**N**) and representative transverse sections from embryos expressing *KLF2-66:hsp68:lacZ* (**O-R**). See also Figure S1B-E. Black scale bars are 200 μm, a = aorta, cv = cardinal vein, jv = jugular vein, ca = carotid artery, av = aortic valve, ot = outflow tract, vv = venous valve, avv = atrio-ventricular valve, ra/la = right/left atria, end = endocardium.

While promoters play an essential role in gene transcription, eukaryotic gene expression also involves additional cis-regulatory elements known as enhancers. Enhancers are short DNA sequences rich in TF binding motifs. Enhancer activation, and subsequent promoter contact and gene transcription, is typically dependent on the binding of multiple different TFs (e.g. lineage-specific TFs alongside signal pathway-specific TFs). This enables an array of different intrinsic and extrinsic cues to collectively influence gene expression patterns (28, 29), and research has identified at least one enhancer for all examined endothelial genes (30). Enhancers are capable of activating gene transcription from a distance, and are often located away from the immediate promoter region. Consequently, to better understand the manner in which FSS and other upstream inputs influence *KLF2* expression in ECs, we examined the *KLF2/Klf2* locus for enhancer marks, including open chromatin (via ATAC-seq), enriched H3K27Ac and p300 binding (31). Additionally, we looked for ETS1 and ERG binding, as enhancers active in ECs ubiquitously bind these EC-expressed ETS family TFs (30, 32). While our aim was to identify human *KLF2* enhancers, we examined enhancer marks in mouse ECs (from E11.5 embryos and adult aorta, heart and lung) alongside cultured human ECs, as this enabled us to study enhancer marks in ECs exposed to endogenous FSS (Figure 1E).

As expected, the *KLF2pm-0.4* promoter was marked by open chromatin, activating histone marks, p300, and ETS TF binding, and was well conserved across species (Figure 1E). In contrast, the region directly upstream of the promoter contained no further promoter/enhancer marks or human-mouse conservation. This analysis also identified two putative enhancers upstream of *KLF2,* both phylogenetically conserved with strong enhancer-associated marks in both human and mouse ECs (Figure 1E). These were named *KLF2-41* and *KLF2-66* in reference to the distance in kb from the human *KLF2* TSS. To assess their ability to independently drive expression, each putative enhancer was cloned upstream of the silent hsp68 minimal promoter-β-galactosidase (*lacZ*) reporter gene and used to generate transgenic mice. Enhancer activity was tested in F0 transgenic embryos examined at E12.5, at which stage blood flow is established and endogenous *Klf2* upregulated in FSS-exposed ECs (Figure 1A-B). Whole-mount F0 embryos transgenic for *KLF2-41:lacZ* and *KLF2-66:lacZ* both exhibited robust vascular-specific expression (Figure 1F-R, S1B-E). Examination of sections through these transgenic embryos indicated that the *KLF2-41* enhancer drove *lacZ* reporter gene expression in arterial and microvascular ECs but not venous ECs (Figure 1I-L and S1C). *KLF2-66* drove similar strong expression in arterial and microvascular ECs while also driving weak expression in some venous ECs (Figure 1N-R and S1E). In the heart, both enhancers were active in endothelium lining the outflow tract and endocardial cushion around the developing valves, while the *KLF2-66* enhancer also consistently drove strong expression in the endocardium (Figure 1K-L and Q-R). Collectively, these expression patterns correlated well with endogenous *Klf2* mRNA in mice at this timepoint (Figure 1A-B and (33)). Of note, the previously described *Klf2-50* mouse enhancer (*mm2219* on the Vista Enhancer Browser (34, 35)) implicated in directing HAND2-dependent gene expression in endocardial ECs, is the direct orthologue of human *KLF2-66* (33). Although the systemic vasculature was not a focus of George *et al*., (33), wholemount and sectional images of *Klf2-50:LacZ/mm2219* activity were similar to *KLF2-66*, with robust *Klf2-50:lacZ* expression in ECs of the dorsal aorta and endocardium and only weak expression in the cardinal vein (33).

Alongside *KLF2-41* and *KLF2-66*, we identified three additional genomic regions with weaker enhancer marks, named *KLF2-34, KLF2-32 and KLF2-8* (Figure 1E). *KLF2-34* and *KLF2-8* were also tested in F0 mouse transgenic models (Figure 1F and S1F-H). Neither directed consistent vascular expression patterns, although a single *KLF2-8:lacZ* embryo (1 of 10 transgenic embryos) had reporter gene activity in smooth muscle, ECs and outflow tract mesenchymal cells (Figure 1F and S1H). The mouse orthologue to KLF2-32 (*Klf2-16/mm2218)* was tested in F0 transgenic mice at E11.5 by George et al., (33) with similarly negative results. In this case, only a single transgenic embryo (1 of 8) showed consistent endocardial activity (33), and this enhancer (as *mm2218*) is recorded as a negative enhancer on the Vista Enhancer Browser (33–35). Consequently, these three elements were not studied further.

### Activity of the KLF2-41 and KLF2-66 enhancers closely mirrors endogenous *Klf2* during development

To further analyse the activity of the two robust *KLF2* enhancers, we generated stable transgenic lines and assessed expression throughout development (Figure 2). The *KLF2-41:lacZ* transgene was active in the vasculature from approximately E8.5, with expression detected (as X-gal staining) in ECs within the dorsal aorta adjacent to the heart (Figure 2A and S2A-B). Ectopic expression was also seen in the developing notochord, likely an off-target consequence of the *hsp68* minimal promoter. By mid-gestation, *KLF2-41:lacZ* expression could be seen throughout the ECs of the dorsal aorta, and in the neural vascular plexus (Figure 2A-C). In contrast, very little *KLF2-41:lacZ* activity was detected in the cardinal or head veins, closely mirroring the arterial– and microvascular-restricted activity pattern seen in the *tg*(*KLF2-41:lacZ)* F0 embryos (Figure 1). This continued through embryonic and early post-natal development (Figure 2A-C and S2D), although the *KLF2-41:lacZ* transgene was expressed only very weakly in adult tissues (a common occurrence with *enhancer:hsp68:lacZ* transgenes linked to epigenetic silencing e.g. (36)). In developing hearts, the *KLF2-41:lacZ* transgene was expressed in the sinus venosus-derived coronary plexus on the dorsal surface of the heart from E12.5 (Figure 2B-C), agreeing with scRNA-seq analysis in SV-derived coronary ECs at similar stages (37), shown in Figure S2E. *KLF2-41* activity in coronary vessels was maintained throughout embryonic development, although it slowly became restricted to coronary vessels within the deeper myocardium, with strongest expression in coronary arterial ECs (Figure 2C). This expression pattern suggests a continuation of the arterial and microvascular-restricted expression associated with *KLF2-41:lacZ* in the systemic vasculature. However, little transgene activity was seen in the coronary vessels in the septum and inner myocardial layer, suggesting that *KLF2-41* was preferentially active in sinus-venosus versus endocardial-derived coronary vessels (Figure 2C). Additionally, *KLF2-41:lacZ* activity in the endocardium was restricted to early embryonic stages, although expression was seen in the ECs of the heart valves at later embryonic and early post-natal stages (Figure 2C and S2C).

**Figure 2.**
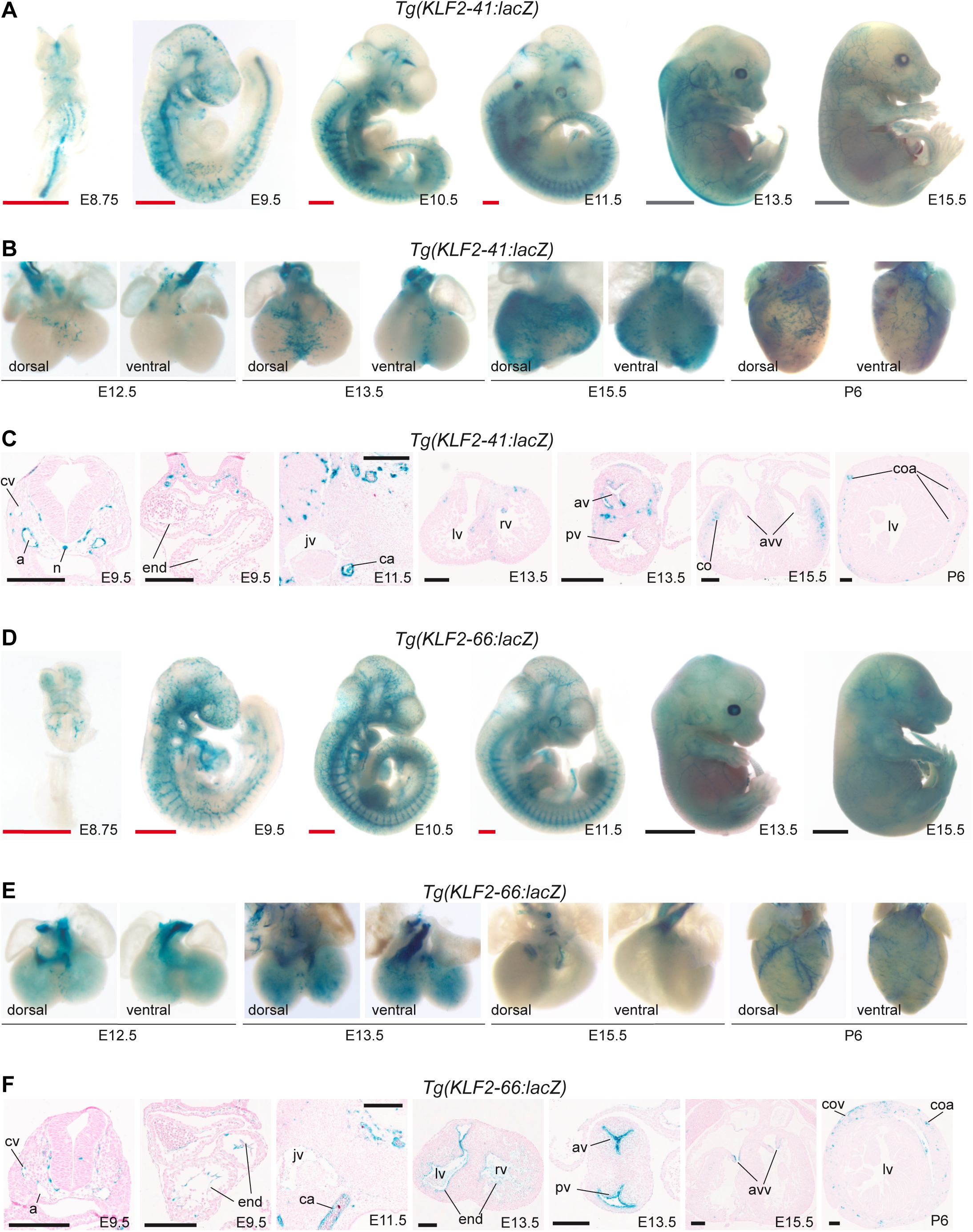
Reporter gene expression throughout development in stable *KLF2-41:lacZ* and *KLF2-66:lacZ* transgenic mice lines recapitulates endogenous *Klf2* expression. **A-C** Reporter gene expression in a stable line expressing *KLF2-41:lacZ* is seen in arterial ECs throughout development. **A** Representative wholemount stable *KLF2-41:lacZ* transgenic embryos. **B** Representative hearts from stable *KLF2-41:lacZ* transgenic embryos and postnatal pups. **C** Transverse sections though stable *KLF2-41:lacZ* embryos and postnatal pups. **D-F** Reporter gene expression in a stable line expressing *KLF2-66:lacZ* is seen in arterial ECs throughout development. **D** Representative wholemount stable *KLF2-66:lacZ* transgenic embryos. **E** Representative hearts from stable *KLF2-66:lacZ* transgenic embryos and postnatal pups. **F** Transverse sections though stable *KLF2-66:lacZ* embryos and postnatal pups. Red scale bars represent 500μm, grey scale bars represent 2mm, black scale bars represent 200μm. a = aorta, cv = cardinal vein, n = notochord, jv = jugular vein, ca = carotid artery, av = aortic valve, pv = pulmonary valve, co = coronary vessels, coa = coronary arteries, cov = coronary veins, vv = venous valve, avv = atrio-ventricular valve, rv/lv = right/left ventricles, end = endocardium. See also Figure S2.

Analysis of the *KLF2-66* enhancer during development also correlated with the F0 results, with strong transgene activity in arterial ECs alongside weaker venous EC expression (Figure 2D-F and S2F-G). Similar to *KLF2-41*, the *KLF2-66:lacZ* transgene was active by E8.5, with expression restricted to the heart-adjacent region of the dorsal aorta and the microvasculature of the head (Figure 2D). While transgene expression was equally detected in both dorsal aorta and cardinal/head veins at E9.5, by E11.5 the transgene was more robustly expressed in arterial compared to venous ECs (Figure 2F). This arterial-enriched expression pattern continued through embryogenesis and into the post-natal stage, although reduced in adult tissues (Figure 2D-F and S2F-G). However, the activity of the *KLF2-66* enhancer in the embryonic heart was significantly different from *KLF2-41*: *KLF2-66:lacZ* activity was not detected in coronary vessels until P0-P6, where expression was seen in both coronary veins and arteries (Figure 2E-F and S2F). Conversely, the *KLF2-66:lacZ* transgene was more active in the endocardium during embryonic stages, although it also became slowly restricted to the ECs of the cardiac valves after E13.5, where it remained robustly active through post-natal stages (Figure 2E-F and S2F).

### Both *KLF2-66* and *KLF2-41* enhancers require blood flow for endothelial activity

Our results so far indicate that the *KLF2-41* and *KLF2-66* enhancers, separated from their native promoter, can still collectively drive reporter gene expression in a pattern similar to endogenous *Klf2* during development. The activity of both the *KLF2-41* and *KLF2-66* enhancers also tracks with the onset of blood flow and the magnitude of FSS, as the mouse embryonic heartbeat is first detected around the 5 somite (E8.25) stage and FSS forces are generally stronger in arteries (38, 39). To directly test their requirement for FSS, we examined enhancer activity *in vivo* in *Ncx1^−/−^* mice, which lack a heartbeat (40). All *Ncx1^−/−^* embryos express *lacZ* throughout the myocardium (due to the *lacZ* transgene inserted into the *Ncx1/Slc8a1* gene to disrupt expression) and are developmentally normal at E9.0, significantly smaller than controls by E10.0 and not viable beyond E12.5 (40). Blood vessels clearly develop in these embryos between E7.5-E9.5 (Figure S3A-B and (4)), providing a window of time to analyse enhancer activity prior to significant growth retardation. We crossed mice transgenic for *KLF2-41:lacZ* and *KLF2-66:lacZ* into the *Ncx1^−/−^* background, directly comparing *lacZ* expression patterns in ≈E9.25 *Ncx1^+/−^* embryos (heartbeat) with littermate *Ncx1^−/−^*embryos (no heartbeat) (Figure 3A-D, S3C-D). *Tg(KLF2-41;lacZ);Ncx1^+/+^* and *tg(KLF2-41:lacZ);Ncx1^+/−^*control embryos (with heartbeat) showed expression in arterial ECs as previously observed (Figure 3A-B and S3C). In contrast, tg(*KLF2-41:lacZ);Ncx1^−/−^*embryos consistently lacked any EC enhancer activity, although ectopic activity of the enhancer in the neural crest was maintained (Figure 3A-B and S3C). Activity of the KLF2-66 enhancer was also drastically reduced in *tg(KLF2-66:lacZ);Ncx1^−/−^*embryos comparative to controls, although some weak residual *lacZ* staining was consistently seen in the microvasculature around the branchial arches (Figure 3C-D and S3D). Interestingly, *KLF2-41:lacZ* and *KLF2-66:lacZ* expression was still seen in some endocardial cells in *Ncx1^−/−^* embryos (Figure 3B and D). This is unlikely to be a consequence of the *lacZ* inserted into the *Ncx1* locus, as despite high X-gal staining in the myocardium, *Ncx1^−/−^* embryos without enhancer transgenes did not have endocardial *lacZ* expression (Figure S3B).

**Figure 3.**
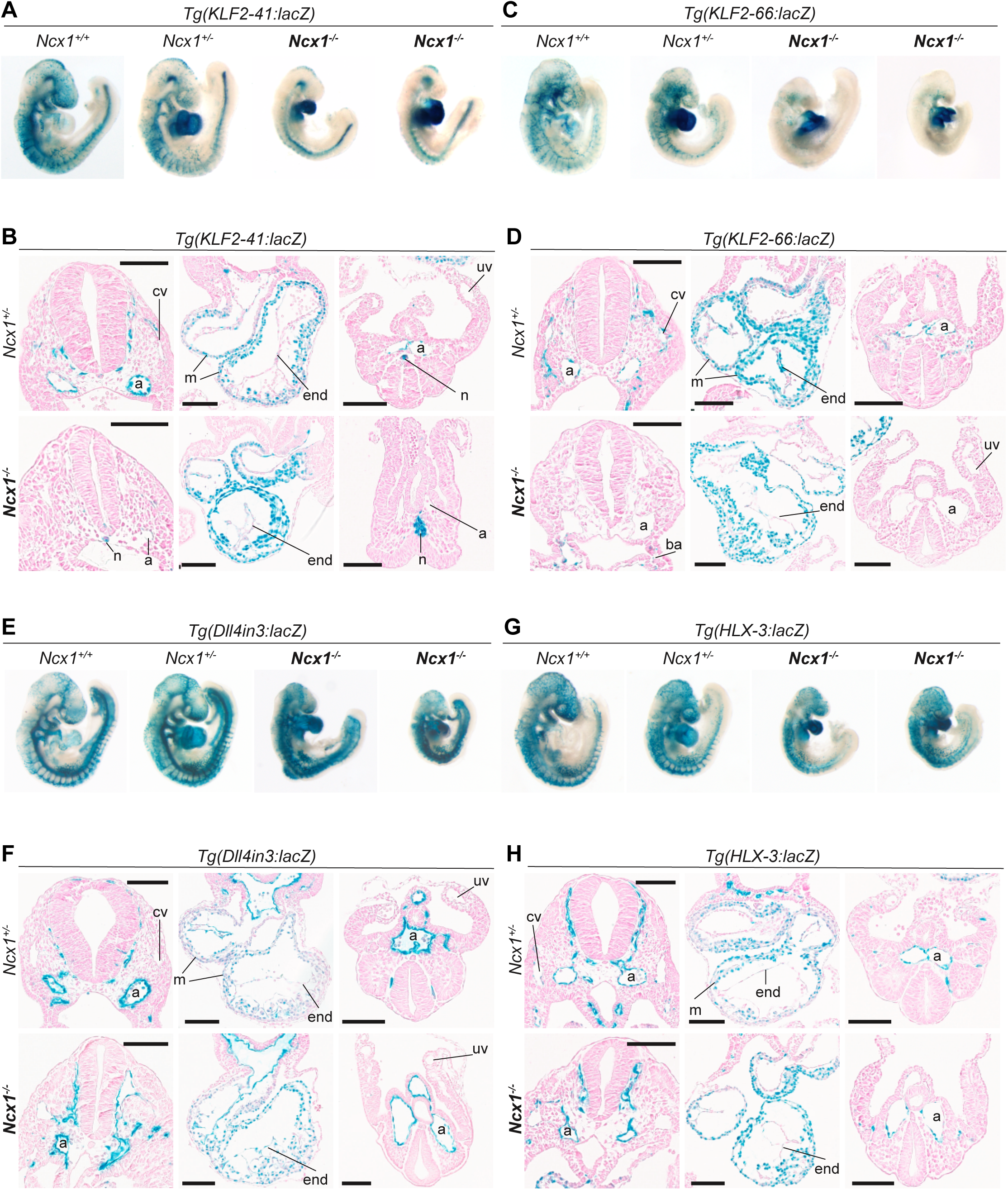
The KLF2-41 and KLF2-66 enhancers require a heartbeat for full activity. Representative E9.25 wholemount embryos (**A**, **C**, **E**, **G**) and transverse embryos sections (**B**, **D**, **F**, **H**) expressing the *KLF2-41:lacZ* (**A-B**), *KLF2-66:lacZ* (**C-D**), *Dll4in3:lacZ* (**E-F**) and *HLX-3:lacZ* (**G-H**) transgenes on a *Ncx1^+/+^* (wildtype), *Ncx1^+/−^*(blue heart and heartbeat) and ***Ncx1^−/−^*** (blue heart no heartbeat) background. The *KLF2* enhancers are not expressed in most ECs in the absence of blood flow while the Dll4in3 and HLX-3 enhancers are unaffected. Black scale bars represent 100μm. a = aorta cv = cardinal vein, n = notochord, m = myocardium, end = endocardium, uv = umbilical vein, ba = branchial arches. See also Figure S3-4.

To determine if these results were specific to *KLF2* or more general across EC enhancers, we also examined the activity of two other EC enhancers on the *Ncx1^−/−^*background: *Dll4in3* and *HLX-3*. These had similar expression patterns to the *KLF2* enhancers, with the *Dll4in3:lacZ* transgene active in arterial and angiogenic ECs, and *HLX-3:lacZ* transgene enriched in arterial ECs at early embryonic stages but also weakly expressed in veins and strongly expressed in angiogenic microvasculature (21, 41). In contrast to the results seen with the *KLF2* enhancers, both the *HLX-3:lacZ* and *Dll4in3:lacZ* transgenes remained active in ECs when crossed into *Ncx1^−/−^* embryos (Figure 3E-H and S4A-B). This agrees with observations in zebrafish models, where expression of neither *Dll4in3* nor *HLX-3* is reduced when flow was perturbed (Figure S4C-D), and previous studies which found increased *Dll4* expression in no flow or loss of flow situations (42, 43). Thus, loss of *KLF2* enhancer is not a general EC enhancer phenomenon.

### TF analysis of KLF2 enhancers

Given the FSS-dependence of the *KLF2-41* and *KLF2-66* enhancers in ECs, we examined their DNA sequences to identify TFs involved in this response. MEF2 factors are already linked to *KLF2* expression, initially from observations of MEF2 proteins binding the *KLF2* promoter *in vitro* (17–19, 27) and reduction of *Klf2* and *Klf4* mRNA in compound *Mef2a;Mef2c;Mef2d* null mice (20). Both *KLF2-41* and *KLF2-66* enhancers contained two A/T-rich sequences that were potential MEF2 binding motifs, each conserved between species to the same extent as the surrounding enhancer region (Figure 4A-B). Two similar motifs were also found in the *KLF2* promoter sequence (Figure 4C). ChIP-seq of MEF2A and MEF2C binding in mouse adult ECs isolated from heart and lung, and in adult heart ventricles found significant MEF2A and MEF2C binding peaks at both *KLF2-41* and *KLF2-66* enhancers in addition to the *KLF2* promoter (Figure 5A).

**Figure 4.**
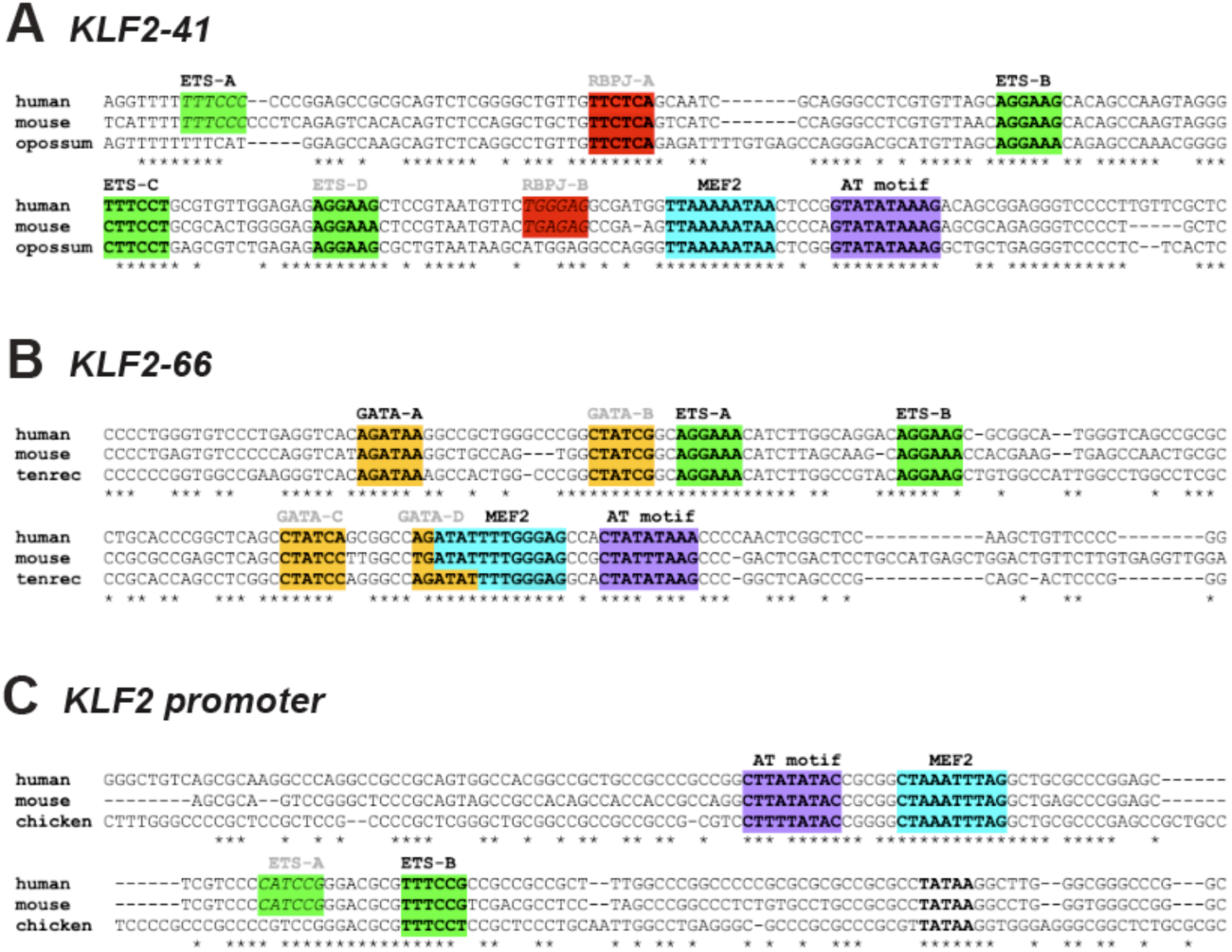
The *KLF2* enhancers and promoter contain well conserved binding sites for multiple TFs. Sequences of the core conserved sequences of the *KLF2-41* enhancer (**A**), *KLF2-66* enhancer (**B**) and *KLF2pr-0.4* promoter (**C**) alongside annotated TF binding motifs. ETS motifs highlighted in green, RBPJ motifs in red, MEF2 in bright light blue, AT motif in purple denotes an A/T rich motif that did not bind MEF2 TFs. Grey text indicates motif not confirmed by ChIP-seq or EMSA, black bold coloured text indicates verified TF binding, bold TATAA indicates TBP motif (TATA box) 31 bp upstream of TSS sites as noted by (26), * indicates conserved nucleotides.

**Figure 5.**
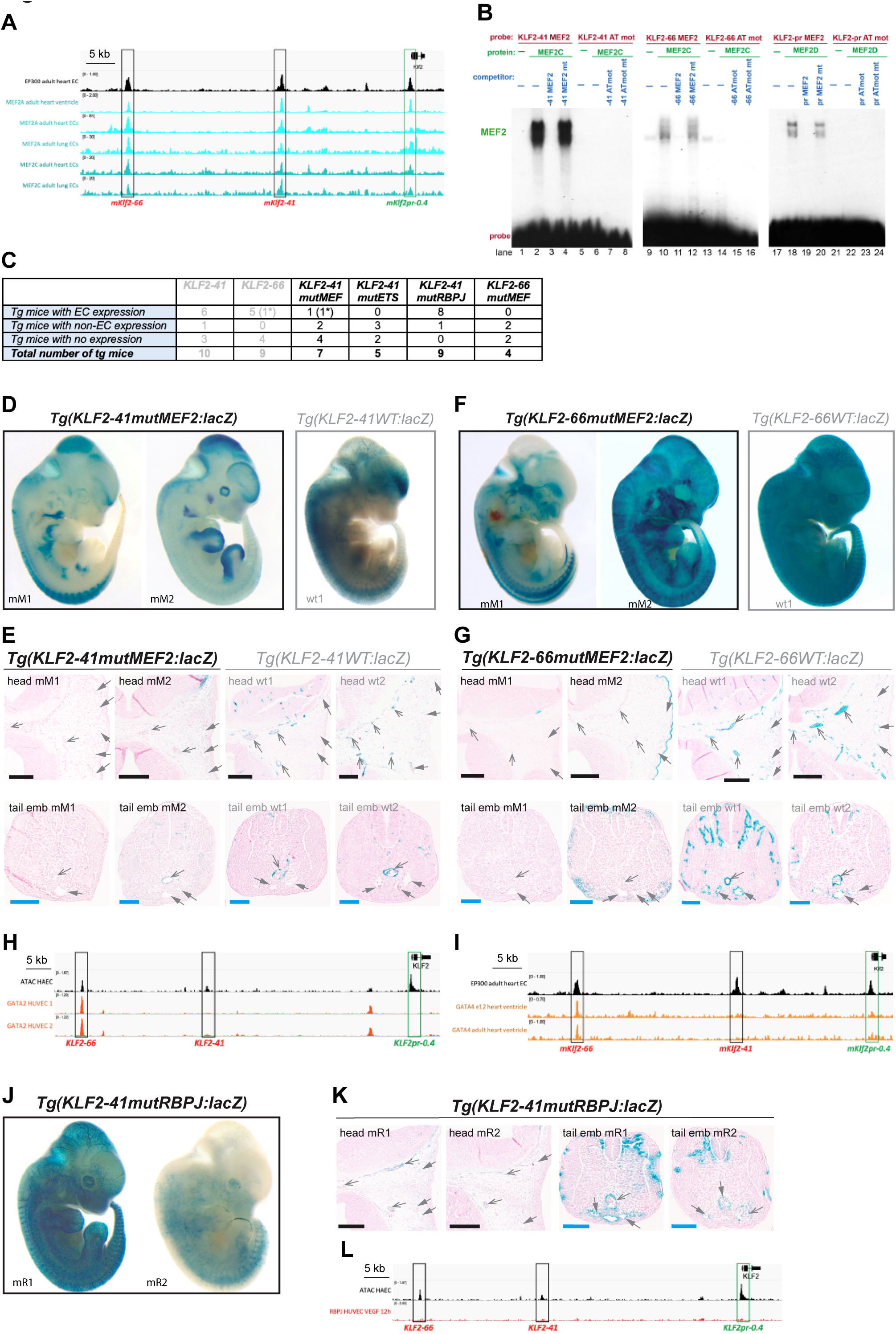
Expression of the *KLF2* enhancers is influenced by MEF2, GATA and RBPJ TFs. **A** MEF2A and MEF2C binding at the *KLF2* locus as assessed by ChIP-seq in adult heart ventricles (from (58)) and adult heart and lung ECs, alongside EP300 from adult heart ECs (from (76)). Binding peaks are seen at the mouse orthologue *Klf2-66*, *Klf2-41* and *Klf2* promoter (black and green boxes). **B** EMSA analysis showing direct binding of MEF2 factors to the MEF2 motifs within each *KLF2* enhancer/promoter element. In each case, the similar A/T rich motif (ATmot) cannot bind MEF2 factors. See also Figure S5. **C** Table indicating expression patterns seen in all F0 mutant transgenic embryos. * denotes weak expression only. **D-G** Mutations to the MEF2 motifs ablate most EC expression of the *KLF2-41:lacZ* transgene (*KLF2-41mutMEF2*, **D** wholemount, **E** transverse sections) and *KLF2-66:lacZ* transgene (*KLF2-66 mutMEF2* **F** wholemount, **G** transverse sections). All embryos at E12.5. See also Figure S5. **H-I** GATA TF factors robustly bind the KLF2-66 enhancer region but not the *KLF2-41* or KLF2-promoter region. GATA2 in HUVECs (**H**) from (84), GATA4 in E12.5 mouse heart ventricles (**I**) (including endocardium) from (86). See also Figure S6. **J-K** Mutations to the RBPJ motifs result in expanded expression of the *KLF2-41:lacZ* transgene to venous ECs (*KLF2-41mutRBPJ*, **J** wholemount, **K** transverse sections), see also Figure S6. **L** RBPJ ChIP-seq in HUVECs after VEGFA stimulation (from (49)) does not show binding to the *KLF2* locus, see also Figure S6 for controls. Black scale bar is 200μm, blue scale bar is 100μm, grey open arrow is artery, grey closed arrow is vein. See also Figure S5 and S6.

Electrophoretic mobility shift assays (EMSAs) were used to determine the ability of each motifs to bind MEF2 proteins. At each enhancer, this identified a single A/T rich motif able to directly bind MEF2 factors (Figure 5B and S5A). Lastly, we constructed transgenes in which these MEF2 binding motifs were mutated (*KLF2-41mutMEF:lacZ* and *KLF2-66mutMEF:lacZ*). *KLF2* enhancers without functional MEF2 motifs drove very little vascular transgene expression, indicating an essential role for MEF2 factors in both *KLF2* enhancer and promoter activity.

MEF2 proteins are widely expressed in numerous non-endothelial cell types (22), and the role of MEF2 factors in *KLF2* promoter activation was first established in non-EC cells (27). MEF2 factors are therefore unlikely to direct KLF2 expression alone in ECs. Consequently, we next investigated potential roles for a range of other EC-associated TFs in the endothelial activity of the *KLF2* enhancers. This analysis combined motif analysis with ChIP-seq/Cut&RUN and EMSA verification (as described in (44)). Our initial enhancer identification already established that both the *KLF2-41* and *KLF2-66* enhancers bound ETS factors (Figure 1E). Sequence and EMSA analysis confirmed that each enhancer contained multiple functional ETS motifs (Figure 4 and S5D-E), while transgenic mouse models demonstrated that a mutant KLF2-41 transgene without functional ETS motifs (*KLF2-41mutETS:lacZ*) was not active in the endothelium (Figure S5F-G). These results align with the known essential role of ETS factors for EC gene expression, and suggest that ETS factors contribute to the EC-restricted pattern of expression of *KLF2*.

In addition to ETS, the *KLF2-66* enhancer contained multiple GATA motifs (Figure 4B), two of which strongly bound GATA2 in EMSA analysis (Figure S6A). Binding of GATA2 and GATA4 to the *KLF2-66* enhancer was also confirmed by ChIP-seq, the latter in cells extracted from adult heart ventricles (Figure 5H-I). However, no GATA binding or motifs were seen at the KLF2-41 enhancer or KLF2 promoter (Figure 4A and C, Figure 5H-I), suggesting that GATA binding to *KLF2* regulatory regions is not a prerequisite for either FSS-responsiveness or EC expression. Instead, the role of GATA in *KLF2* regulation may be specific to endocardial expression: GATA4 has been shown to drive gene expression in endocardium (45), and the GATA4-bound KLF2-66 enhancer was strongly active in the developing endocardium while the KLF2-41 enhancer (no GATA) was not (Figure 2). Previous analysis links the HAND2 TF to endocardial KLF2 expression but the loss of HAND2 did not fully ablate *Klf2* endocardial expression (33), indicating that other TFs such as GATA4 may also contribute to endocardial *KLF2* expression.

As the KLF2-41 and KLF2-66 enhancers are more active in arteries, we also investigated potential roles for the arterial-associated RBPJ/NOTCH and SOXF TFs. Despite the close links between SOXF and arterial transcription in other settings (30, 32), we found no SOX consensus or near-consensus motifs within the core regions of the *KLF2* enhancers or promoter (Figure 4). In contrast, the *KLF2-41* enhancer contained multiple potential RBPJ motifs (Figure 4A). Mutation of these motifs (generating the *KLF2-41mutRBPJ:lacZ* transgene) did not significantly affect transgene expression in arterial ECs (Figure 5J-K, Figure S6B-C). This agrees with previous observations that FSS-induced *KLF2/4* expression was unaffected by NOTCH blockade (43, 46). However, the mutated *KLF2-41mutRBPJ* enhancer was no longer arterial specific in these mice, with reporter gene expression consistently seen in many venous ECs (Figure 5K and S6C). This suggests that these motifs may play a role in limiting enhancer activity in venous ECs rather than activating it in arterial ECs. RBPJ is known to act as a repressor in the absence of Notch signalling in other cell types (47), and a similar vein-expansion phenotype was observed in an arterial-specific enhancer for *Flk1/VEGFR2* after RBPJ motif mutation(48). However, we could not fully establish whether the RBPJ motifs within *KLF2-41* directly bound RBPJ: although they competed for RBPJ binding in competition EMSA analysis, they were unable to directly bind RBPJ in EMSA (Figure S6D). Further, ChIP-seq data from HUVECs exposed to VEGF (from (49)) did not find RBPJ binding at any *KLF2* element (Figure 5L), although this assay also failed to show binding at the RBPJ-repressed *Flk1* enhancer, suggesting that the culture conditions may not be appropriate (Figure S6E). Therefore, while it is clear that these motifs restrict activity of the *KLF2-41* enhancer in venous ECs, it is still undefined whether this occurs via RBPJ, a combination of RBPJ with other TFs, or undefined TFs that recognize the same motif.

### All *KLF2* regulatory elements contain a common and essential MEF2-TBP double motif

Our analysis so far shows that ETS and MEF2 factors bind to all three *KLF2* regulatory elements. However, it is unlikely that these TFs alone control *KLF2* enhancer and promoter responsiveness to FSS. ETS factors regulate many EC enhancers with no association with FSS induction, including some specifically active in vein and lymphatic ECs exposed to lower FSS (30, 50). Similarly, while previous studies associated MEF2 factors with FSS-responsiveness, MEF2 TFs are also linked to angiogenic gene activation (21, 44). Crucially, a number of angiogenic enhancers (including *Dll4in3*, *HLX-3* and *ETS1+195*) require both MEF2 and ETS factors (21), yet there activity is not affected by loss of flow (Figure 3E-H, S4 and S7A). We therefore used HOMER analysis to identify additional TFs that may bind both *KLF2* enhancers and promoter to influence FSS-responsiveness. In addition to MEF2 and ETS, this analysis identified motifs for TATA-binding protein (TBP) within all three regulatory elements (Figure 6A). Similar analysis using JASPAR (51) and TRANSFAC (52) also found TBP motifs in each of the *KLF2* regulatory elements (Figure 6B and S7B). TBP is best known as a component of the transcriptional machinery at the core promoter, where it binds alongside RNA polymerase II and other general TFs (GTFs) (53). However, a role for TBP alongside MEF2 in gene transcription has some precedent in non-mammalian models: in *Drosophila,* differential co-binding of TBP with MEF2 downstream of phosphorylation drives the ability of MEF2 to switch from activating metabolic genes to activating immune genes (54).

**Figure 6.**
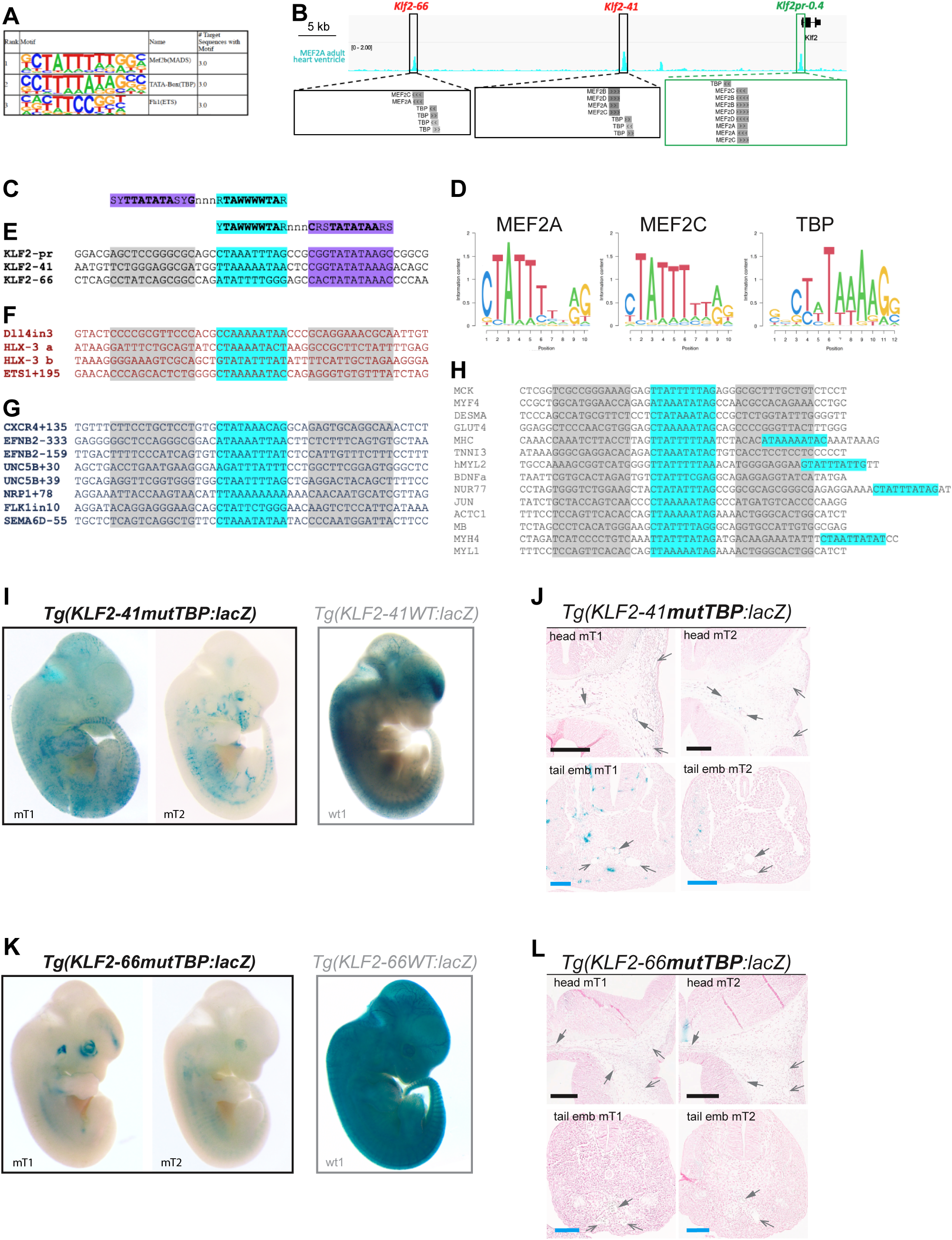
A shared and required MEF2-TBP double motif found within *KLF2-41*, *KLF2-66* and the *KLF2* promoter. **A-B** Homer analysis (**A**) and JASPAR analysis (**B,** via the UCSC browser) both identifies MEF2 and TBP motifs within each of the *KLF2* enhancers and promoter. Each double motif aligns with the MEF2A ChIP-seq (from (58)). **C-H** Consensus motifs for MEF2 and TBP factors (**C**, defined by HOMER, and consensus double motif (**D**)) found within the *KLF2-41*, *KLF2-66* and *KLF2* promoter (**E**). In each case, the MEF2 motif (highlighted in cyan) is separated by a 3 bp spacer sequence from the TBP motif (purple). Similar double motifs are not found in other known MEF2-bound EC enhancers (**F**), other known MEF2-motif containing arterial enhancers (**G**) or other known MEF2-bound non-EC enhancers (**H**). Grey shading indicates locations where TBP motifs would be expected. S indicates A, T or C nucleotides, Y indicates G or C, n indicates A, G, C or T, R indicates G or A. **I-L** Mutations to the TBP component of the MEF2-TBP double motif ablates most EC activity of the *KLF2-41:lacZ* transgene (*KLF2-41mutTBP*, **I** wholemount, **J** transverse sections) and *KLF2-66:lacZ* transgene (*KLF2-66 mutMEF2,* **K** wholemount, **L** transverse sections), all embryos are E12.5. * denotes weak expression only. Black scale bar is 200μm, blue scale bar is 100μm, grey open arrow is artery, grey closed arrow is vein. See also Figure S7.

Intriguingly, our sequence analysis also highlighted the repeated and precise spatial arrangement of the TBP motif alongside MEF2 motifs (Figure 6B and S7B): in each *KLF2* regulatory element, a TBP motif is directly adjacent to validated MEF2 binding motifs, forming a MEF2-TBP double motif with a precise 3 bp spacer in between (Figure 6C-E). The MEF2 and TBP motifs were also deeply conserved across species within each element alongside the precise spacing, with the sole variation being a switch to a 2 bp spacer in the zebrafish version of *KLF2-41* (Figure S7C).

Analysis of other MEF2-bound regulatory elements suggests that MEF2-TBP motifs are not a shared feature. The FSS-insensitive *Dll4in3*, *HLX-3* and *ETS1+195* EC enhancers lack TBP motifs near their MEF2 sites; and double MEF2-TBP motifs were not seen in eight other MEF2-bound EC enhancers described by Nornes *et al*., (44) (Figure 6F-G). Further, no MEF2-TBP double motif was found within 14 MEF2-driven promoters and enhancers previously defined in other tissue types, although a number of these contained a spatially and sequentially distinct MEF2-MEF2 double motif associated with activation in some cell types (Figure 6H and S7D) (55, 56).

Our earlier analysis already demonstrated the importance of the MEF2 motifs to *KLF2* enhancer activity (Figure 5). We therefore next examined whether the TBP component of these MEF2-TBP motifs was required for enhancer activity. We generated *KLF2-41mutTBP* and *KLF2-66mutTBP* transgenes, each bearing small mutations specifically within the TBP motif. In transgenic embryos, mutation of the TBP motif resulted in loss of reliable enhancer:reporter gene expression in the vasculature for both enhancers, although a minority of embryos transgenic for the *KLF2-41mutTBP:lacZ* transgene had some reporter gene expression in ECs (Figure 6I-L and S7E-H). Intriguingly, Sohn *et al.* (18) also identified an essential role for the TBP component of the MEF2-TBP double motif within the KLF2 promoter, although at the time it was identified as “A/T-rich motif A” with no known binding TF. This analysis found that selective mutation of the TBP motif ablated activation of the promoter by ERK5-MEK5 signalling, and consequently, FSS (18). These combined results strongly suggest that both MEF2 and TBP components of the MEF2-TBP double motif located within all *KLF2* regulatory elements play important roles in endothelial FSS-driven gene expression.

### TBP differentially binds the KLF2 regulatory elements in response to FSS

To confirm the results from sequence analysis, we next assessed binding of TBP at the *KLF2* locus, first using a TBP ChIP-seq dataset generated in WT postnatal mouse liver, where approximately 20% of cells are ECs (57). Significant TBP binding was observed at both *KLF2* enhancers in addition to the *KLF2* promoter, consistent with the motif analysis (Figure 7A). Other components of the transcription complex examined in the same assay (RNA pol II, TFIIB and TFIIE) did not show similar levels of binding at the *KLF2* enhancers, suggesting that TBP binding at these elements was not the sole consequence of intergenic transcription (Figure 7A). TBP binding to the *KLF2* enhancers was also not seen in ChIP-seq datasets from other cell types, suggesting EC specificity (Figure S8A). To study the effects of FSS on TBP binding, we analysed TBP binding in immortalized human aortic ECs (telo-HAECs) without flow or under physiological FSS using CUT&RUN for endogenous TBP. TBP binding at the *KLF2* regulatory elements was assessed by both qRT-PCR and whole genome sequencing. In static ECs, we found significant binding for TBP at both *KLF2* enhancers and the *KLF2* promoter, agreeing with the earlier liver TBP ChIP-seq (Figure 7B). Notably, FSS consistently increased TBP binding at the KLF2 regulatory elements (Figure 7B-C). This was less pronounced at the promoter, which will also bind TBP as part of the transcriptional preinitiation complex. In keeping with the specificity of the MEF2-TBP motif, TBP binding was not common to all EC enhancers (Figure S8B-C). In conclusion, TBP binds alongside MEF2 factors at all regulatory elements within the *KLF2* locus in a FSS-sensitive pattern.

**Figure 7.**
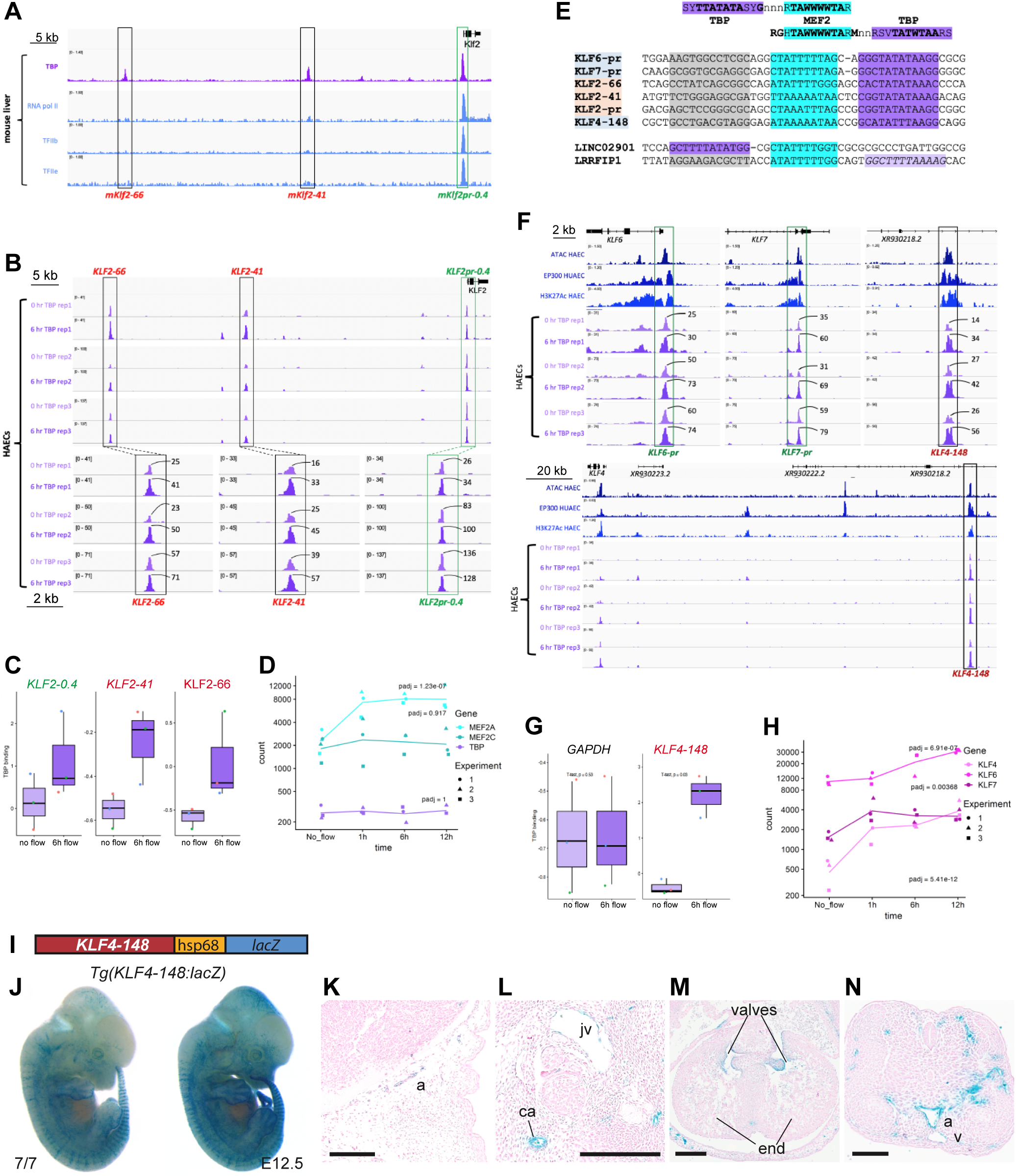
TBP binding to MEF2-TBP elements increases with flow and is predictive of regulatory elements within other *KLF* genes. **A** TBP selectively binds all three regulatory elements (black and green boxes) within the *Klf2* locus as assessed by ChIP-seq in postnatal whole livers although other components of the basal transcriptional machinery (RNA polymerase II, TFIIb and TFIIE) only significantly bind the *Klf2* promoter. Data from (57). **B-C** TBP binding at the *KLF2* regulatory elements in telo-HAECs increases after 6 hours of FSS assessed by CUT&RUN followed by whole genome sequencing (**B**) or qRT-PCR (**C**). **D** *TBP* and *MEF2C* mRNA levels do not increase after FSS, MEF2A mRNA increases. **E** Whole genome analysis of MEF2– and TBP-bound regions identified MEF2-TBP motifs within other *KLF* regulatory elements. These MEF2-TBP motifs shared sequence orthology with the *KLF2* motifs. See also Table 1 and Table S1-2. **F** MEF2-TBP-containing elements within the *KLF6*, *KLF7* and *KLF4* loci contained enhancer/promoter marks (ATAC-seq and H3K27Ac in HAECs (78) and EP300 HUAEC from (79)) and bound TBP, with greater binding seen after 6 hrs of FSS. **G** TBP binding at the *KLF4-148* MEF2-TBP containing enhancer increased after FSS as assessed by CUT&RUN followed by qRT-PCR. **H** *KLF4, 6* and *7* mRNA levels increase after FSS. **I-N** The *KLF4-148:lacZ* transgene (**I**) drives strong EC specific expression in the arteries and microvasculature of F0 E12.5 transgenic embryos as depicted in representative wholemount images (**J**) and transverse sections (**K-N**). See also Figure S10. Numbers next to each peak indicate peak height as assessed on IGV, black boxes denote enhancers, green boxes denote promoters, experimental repeats were biological replicates run on separate days. Black scale bars are 200 μm, a = artery cv = jv = jugular vein, ca = carotid artery, v = tail vein, end = endocardium.

**Table 1.** List of genomic locations bound by both MEF2A and TBP and containing MEF2A and TBP motifs separated by 2-4 bp in mouse and/or human. Light pink shading indicates known MEF2-TBP motifs within the *KLF2* regulatory elements, light blue shading indicates novel motifs within other *KLF* genes. See also Table S3. Bold numbers indicate evolutionary conserved distances between the TBP and MEF2 motifs in human and mouse.

| Nearest gene hg38 | Neighbouring genes hg38 | Distance btw motifs hg38 | Distance btw motifs mm10 | Annotated location hg38 | TBP peak location hg38 | MEF2A peak location mm10 |
| --- | --- | --- | --- | --- | --- | --- |
| LINC02901 | OSTCP1; RSPH3 | 2 | 2 | Promoter (<=1kb) | chr6 158869701 158870000 | chr17 6827841 6828061 |
| **KLF6** | LINC02660; LINC02669 | 2 | 2 | Promoter (<=1kb) | chr10 3784934 3786509 | chr13 5861130 5861331 |
| **KLF7** | CPO; MYOSLID | 2 | 2 | Promoter (<=1kb) | chr2 207165508 207166950 | chr1 64121704 64121931 |
| AP1M1 | **KLF2**; FAM32A | 3 | 3 | Distal Intergenic | chr19 16258401 16258975 | chr8 72268461 72268713 |
| **KLF2-dt** | **KLF2**; AP1M1 | 3 | 3 | Exon | chr19 16283151 16284100 | chr8 72295792 72295994 |
| **KLF2** | AP1M1; EPS15L1 | 3 | 3 | Promoter (<=1kb) | chr19 16324451 16325050 | chr8 72318832 72319054 |
| LOC105376207 | **KLF4** | 3 | 3 | Exon | chr9 107637351 107639043 | chr4 55651158 55651370 |
| LRRFIP1 | RAB17; RBM44 | 4 | 4 | Intron | chr2 237616851 237617250 | chr1 90980737 90980960 |

The increased TBP binding at *KLF2* elements after FSS was not explained by a concurrent increase in TBP transcription, which was unaltered by FSS (Figure 7D). This indicates that FSS alters the ability of the TBP protein to bind the MEF2-TBP motif. *MEF2A* expression was increased by FSS (Figure 7D), but the MEF2A promoter neither contains a MEF2-TBP double motif or binds significant TBP in either conditions, nor could TBP binding be seen elsewhere at the locus (Figure S8D-E). However, the MEF2A promoter contains multiple KLF motifs and bound KLF4 in a FSS-dependent manner, suggesting that increased *MEF2A* mRNA is secondary and occurs indirectly via MEF2/TBP induction of *KLF2/4* (Figure S8E). *MEF2C* levels show no change after FSS, and the *MEF2C* promoter and enhancers bind no KLF factors, contained no MEF2-TBP motifs nor differentially bound TBP (Figure S8F).

### The MEF2-TBP double motif is rare within the genome yet found within multiple KLF regulatory elements

One of the striking aspects of FSS-induced *KLF2* expression is the specificity of this regulation despite the widespread expression of TBP, MEF2 and ETS factors. Our results so far indicate that the MEF2-TBP double motif may be the driver of this specificity, since these motifs are found within all three FSS-sensitive *KLF2* regulatory elements but not in other unrelated EC and/or MEF2-driven enhancers (Figure 6C-H). To further investigate, we next determined how often this functional motif occurs across regulatory elements in the genome. We first examined all called peaks for TBP binding in HAECs, looking for peaks in which motifs for both TBP and MEF2 were called (Table S1). This analysis identified 16480 TBP-bound peaks, 3295 of which contain TBP motifs. 829 of these peaks also contained called MEF2 motifs, of which only 101 peaks contained MEF2-TBP motifs with a spacer of 10 bp or less. This analysis was repeated for called peaks for MEF2A binding in mouse heart ventricles (58). This ChIP-seq dataset identified 6725 peaks, of which 5881 contained called MEF2 motifs, 861 contained both MEF2 and TBP called motifs, and 167 contained MEF2-TBP double motifs with a spacer of 10 bp or less (Table S2). Our previous analysis of the MEF2-TBP double motif in the *KLF2* locus found that the spacer length was strongly conserved across species, with only one single bp change (Figure S7C). We therefore focused on MEF2-TBP motifs with spacers of 2-4 bp. This analysis identified 24 MEF2-TBP motifs within TBP peaks and 41 MEF2-TBP motifs within MEF2A peaks, with only eight MEF2-TBP motifs with a 2-4 bp spacer bound by both TBP and MEF2 (Table 1 and Tables S1-3). Of these, three were the known MEF2-TBP motifs from *KLF2-66*, *KLF-41* and *KLF2pr0.4*. Strikingly, of the remaining five MEF2-TBP double motifs, three were associated with other *KLF* genes (Table 1). In each case, these MEF2-TBP motifs showed strong similarity to those within the *KLF2* elements (Figure 7E). They were also highly conserved across species, with the MEF2-TBP double motif showing deeper conservation that the surrounding region (Figure S9A).

We next sought to verify these results using genomic datasets not included in the original analysis: TBP binding in mouse liver; enhancer/promoter marks in mouse ECs; MEF2A/C binding in adult mouse ECs; and enhancer/promoter marks in human ECs. The three newly identified *KLF*-associated MEF2-TBP elements were bound by MEF2 and TBP in each assay, and were strongly correlated with enhancer/promoter marks (Figure 7F and Figure S9B). Analysis of TBP binding also demonstrated increased binding peaks at each of these elements after FSS (Figure 7F-G). In agreement with our hypothesized link between binding at the MEF2-TBP motif and FSS-sensitivity, transcription of *KLF4* increases after FSS (Figure 7H and (16, 59)). Although less is known about *KLF6* and *KLF7* in ECs, both genes were also expressed in cultured HAECs, with their expression increasing after FSS alongside *KLF4* (Figure 7H).

In addition to the MEF2-TBP motifs associated with the *KLF6*, *KLF7* and *KLF4* genes, we also identified two other regions with bound MEF2-TBP motifs with 2-4 bp spacers (*LINC02901-pr* and *LRRFIP1-10*, Figure 7E and Table 1). These motifs shared fewer similarities to the *KLF2* MEF2-TBP motifs: the *LINC02901*-associated motif was in the opposite orientation, and the TBP motif within *LRRFIP1*-*10* was divergent (Figure 7E). Both were also poorly conserved (Figure S9C-D), had fewer enhancer/promoter marks and less MEF2 binding (Figure S9B). TBP binding at both elements was also weaker than for *KLF*-associated elements, and did not increase with FSS (Figure S9E). The *LRRFIP1*-10 element was within the *hs1945* enhancer previously identified by the Vista Enhancer Browser consortium (34, 35) (Figure S9F). As *hs1945* was active only in the outflow tract of these embryos, it is possible that the *hs1945* MEF2-TBP motif within this enhancer functions as a flow-dependent element in a different cell type.

Our analysis so far investigated the genomic distribution of MEF2-TBP motifs within called binding peaks for MEF2 and TBP. Next, we investigated the presence of MEF2-TBP motifs within EC-associated enhancer and promoter elements genome-wide in a manner that was agnostic to MEF2 or TBP binding. This analysis utilized enhancer/promoter-associated chromatin marks (as in the initial identification of KLF2 regulatory elements, Figure 1E) to define EC regulatory elements, and the GLAM2SCAN tool to identify sequences with high similarity to the MEF2-TBP double motifs within the six *KLF*-associated elements. In agreement with our earlier results, very few regions with similarity to the MEF2-TBP double motif were found (Table S4-S5), further supporting a specific link between the presence of MEF2-TBP double motifs and FSS-induced *KLF* gene expression.

### The MEF2-TBP double motif-containing KLF4-148 sequence functions as an EC-specific enhancer in mouse models

While the MEF2-TBP double motifs for *KLF6* and *KLF7* were located in promoter regions, the MEF2-TBP double motif associated with *KLF4* was located 148 kb upstream of the TSS. This distal region was implicated in *KLF4* regulation by Maejima *et al.* due to observed binding of MEF2C and looping with the *KLF4* promoter (60). However, *KLF4* is expressed in multiple tissue types and this element was never tested *in vivo*. To verify whether our MEF2-TBP motif analysis was sufficient to predict an endothelial-specific regulatory element, we cloned the conserved region around the *KLF4-148* MEF2-TBP binding motif upstream of *hsp68:lacZ* and tested in transgenic mice as before. Analysis in E12.5 mouse embryos demonstrated that this KLF4-148 element functions as a robust EC-specific enhancer *in vivo*, with EC expression seen in all seven F0 transgenic embryos (Figure 7I-N and S10). Expression was detected in both arterial and microvascular beds, the latter correlating with the known role of KLF4 in angiogenesis (59). This result therefore strongly supports an important and specific role for the MEF2-TBP double motif in transcriptional regulation of *KLF* genes downstream of FSS.

## Discussion

Despite a wealth of research linking FSS with increased *KLF2*/*KLF4* expression, and the subsequent cascade of cardiovascular events triggered by these factors, the transcriptional mechanisms through which this occurs remain incompletely understood. Here, our detailed analysis of *KLF2* regulatory elements implicates TBP as an essential transcriptional regulator of FSS-induced *KLF* gene regulation, working alongside MEF2 factors at a distinct and specific double TF motif. The limited genomic distribution of the MEF2-TBP double motif also explains how FSS selectively activates *KLF* gene expression despite the widespread expression of both MEF2 and TBP.

Evidence linking FSS-responsiveness of the *KLF2* promoter with a nucleotide sequence encompassing our MEF2-TBP double motif first emerged in 2004 (25, 26). Subsequent research found that this sequence was required for FSS-induced MEK5/ERK5 modulation of *KLF2* promoter activity (18, 61), showed that MEF2 factors bound one component of this sequence (18, 19) and were required for FSS-induced promoter activation (61). Crucially, these studies observed that the entire sequence was required for full FSS induction, with neither the MEF2 motif nor the neighbouring AT-rich motif was alone sufficient for MEK5/ERK5 responsiveness (18, 26, 27). However, the protein bound adjacent to MEF2 was never defined, and the canonical MEK5/ERK5/MEF2 model for FSS-induced expression of *KLF2* omits the fact that additional TFs are involved. There was also a disconnect between the high transcriptional specificity assumed in this model, and the widespread expression and transactivation of MEF2 factors both within and beyond the vasculature (22). Here, the discovery of near-identical and highly conserved MEF2-TBP double motifs within regulatory elements for the FSS-responsive *KLF2*, *KLF4*, *KLF6* and *KLF7* genes emphasizes the essential role of both components of this element, and the necessity to better understanding the TFs that bind it.

The similarity of the *KLF2* promoter FSS-responsive sequence to a TBP motif (then referred to as a TATA box) was noted in previous studies but dismissed. This was influenced by the inability of this motif to bind other components of the transcriptional preinitiation complex (62), and its distal location relative to the TSS. Additionally, research linking TBP to differential MEF2 binding and resultant transcriptional switching in *Drosophila* was only published in 2013 (54). In this paper, Clark *et al*., show that the ability of MEF2 factors to drive two distinct gene expression states is regulated by changes in phosphorylation and subsequent altered ability to bind with TBP at a single shared motif (54). Although the *KLF*-associated MEF2-TBP double motif is distinct from those in the *Drosophila* model, there are some key similarities. MEF2 factors also drive two different gene programmes in the endothelium via distinct motifs, transactivating angiogenic genes via a single consensus MEF2 motif (to contribute to an activated endothelial state), while alternatively transactivating *KLF* genes via the MEF2-TBP double motif (to contribute to a homeostatic state). Additionally, the link between FSS, MEK5/ERK5 phosphorylation and MEF2 transactivation suggests that MEF2-TBP-driven gene activation may similarly be influenced by phosphorylation. Given the increased TBP binding to MEF2-TBP double motifs after FSS, one potential model may be that ERK5-driven MEF2 phosphorylation increases its ability to interact with TBP at the MEF2-TBP double motif. However, the influence of ERK5 on MEF2 is multifaceted and still incompletely understood. While ERK5 can phosphorylate MEF2 factors as part of EGF-induced ERK5-MEF2-*JUN* and ERK5-MEF2A-*PFKFB3* transcriptional pathways (63, 64), ERK5 kinase inhibitors paradoxically stimulate MEF2 reporter and *KLF2* promoter activity by enhancing MEF2 nuclear translocation(65). Additionally, there are numerous annotated but uncharacterized MEF2 phosphorylation sites that may also contribute to altered MEF2-TBP complex binding downstream of FSS, and TBP itself may be phosphorylated. Local environmental stimuli can also influence MEF2 transactivation in other ways. In particular, HDACs can inhibit MEF2 activation of *KLF2* expression downstream of NF-κB signaling, and a link between HDACs and MEF2 is established in angiogenic ECs (19, 21). Although transcription of *MEF2A* also increases under FSS, this appears to occur indirectly via the increased transcription of *KLF* factors, therefore reinforcing rather than activating this pathway. Additionally, we found no evidence of increased levels of MEF2C, which also binds and activates *KLF* genes and compensates for MEF2A in gene ablation studies (20).

Although its binding is not restricted to promoter regions (66), TBP is primarily known as a component of the transcription preinitiation complex (PIC) at core promoters. This link was first established by the discovery of TBP motifs (also known as TATA boxes) adjacent to transcriptional start sites, although TBP binding at core promoters does not require a consensus TBP motif (67). However, the locations of the MEF2-TBP motifs within the *KLF2, KLF6* and *KLF7* promoters suggest that they are not directly involved in the PIC: core promoter TBP motifs must be located ≈30 bp upstream of the TSS (68), whereas the MEF2-TBP motif is 110-134 bp upstream of the *KLF2* TSS, 322-345 bp upstream of the *KLF6* TSS, and 389-412 bp upstream of the *KLF7* TSS. Further, all three genes have additional TBP-like motifs located correctly within the core promoter (e.g. *KLF2*, see Figure 4 and (26)). Consequently, the promoter MEF2-TBP motifs appear to act as an additional element rather than a misplaced TATA box. Potential involvement of the enhancer MEF2-TBP motifs in transcriptional initiation is harder to rule out, as enhancers also initiate RNA transcription to generate enhancer RNAs (eRNAs), a process which can involve TBP proteins and motifs (69). Unlike at core promoters, the site of enhancer transcriptional initiation can be challenging to establish due to bidirectional transcription, instability of eRNA transcripts, and transient occupancy of components of the PIC (70, 71). However, the *KLF2* and *KLF4* enhancers display much stronger TBP binding than other non-FSS-sensitive endothelial enhancers, despite TBP and RNA pol II binding at the promoters of all. We also detected only very weak RNA pol II and TFIIb/TFIIe binding at the *KLF2* enhancers compared to the robust TBP binding. These results further suggest that TBP bound alongside MEF2 at the KLF-associated double motif acts as a traditional TF rather than a component of the PIC.

In addition to the MEF2-TBP double motif, our analysis also implicates other TFs in aspects of *KLF2* expression within the vasculature. This was not unexpected, as precise spatio-temporal gene expression often involves combinations of TFs (72). In particular, our results strongly indicate a key role for ETS factors in directing the EC-specificity of the KLF enhancers, consistent with extensive previous evidence of essential roles for ETS in endothelial-specific gene expression (30). This also explains the residual EC expression seen in a small number of transgenic mice after mutation of either MEF2 or TBP motifs. In addition, our data indicates a potentially crucial role for GATA factors in driving *KLF2* expression in heart valves. GATA4 is known to be strongly expressed in the endocardium, and was able to directly bind the valve-expressed KLF2-66 enhancer. This may occur alongside HAND2, which is itself essential for valve morphogenesis and also directly linked to the KLF2-66 enhancer (33). Such an interaction has precedent, as GATA4 and HAND2 are already known to synergistically activate gene transcription in other cell types, including myocardium (73).

One of the most striking aspects of our results was the strength of association between the MEF2-TBP double motif and FSS-responsive *KLF* gene regulatory elements. This provides a mechanistic explanation by which the widely-expressed MEF2 factors are able to specifically target KLF gene expression, and explains the marked similarities between *Mef2a/c/d* and *Klf2/4* EC null mouse models. Additionally, given the paramount importance of *KLF2/4* expression in suppressing atherosclerosis, the uniqueness of the MEF2-TBP double motif presents a promising target for therapeutic modulation of *KLF* levels without off-target effects.

## Methods

### Mice and zebrafish models

All animal procedures were approved by local ethical review and licensed by the UK Home Office. E12.5 F0 transgenic embryos were generated, dissected and stained in X-gal by Cyagen Biosciences. *Ncx1^−/−^* mice were from (40), *Tg(Dll4in3:lacZ)* from (41), *Tg(HLX-3:lacZ)* from (21). Stable *Tg(KLF2-41:lacZ)* and *Tg(KLF2-66:lacZ)* transgenic mouse lines were generated on a C57BL/6 background by pro-nuclear oocyte microinjection of linearized DNA. Zebrafish lines *tg(Dll4in3:GFP)*, *tg(hlx-3:GFP)* and *tg(ETS1+194:GFP)* were from (21, 41).

For analysis, mouse samples were collected along with yolk sac/tail tip, and fixed in 2% paraformaldehyde, 0.2% glutaraldehyde, 1X PBS for 10-120 mins according to sample size. After fixation, samples were rinsed in 0.1% sodium deoxycholate, 0.2% Nonidet P-40, 2 mM MgCl2 in 1 X PBS, then stained for 2-24 hours in 1 mg/ml 5-bromo-4-chloro-3-indolyo-β-D-galactoside solution (X-gal) containing 5 mM potassium ferrocyanide, 5 mM ferricyanide, 0.1% sodium deoxycholate, 0.2% Nonidet P-40, 2 mM MgCl_2_ and 1 X PBS. After staining, embryos were rinsed through a series of 1 X PBS washes, then fixed overnight in 4% paraformaldehyde at 4°C. Embryos were imaged using a Leica M165C stereo microscope equipped with a ProGres CF Scan camera and CapturePro software (Jenoptik). For histological analysis of transgenic mice, samples were dehydrated through a series of ethanol washes, cleared by xylene and paraffin wax-embedded. 5 or 6-μm sections were prepared and de-waxed. For imaging of X-gal staining, slides were counterstained with nuclear fast red (Electron Microscopy Sciences). Analysis was qualitative not quantitative, therefore no statistical analysis was applied to the observations of staining intensity and pattern. Numbers of transgenic mice used followed precedent set by similar published papers. For F0 transgenic analysis, representative wholemount images are shown in the main figure but images of every X-gal-positive embryo is included in the Supplementary Figures. No experimental randomization or blinding was used as this was not considered necessary.

The yolk sac or tail tip was used for genotyping. In cases where genotyping results from yolk sac were ambiguous (e.g. due to contamination), tissue samples from the embryo were removed for genotyping. Each tissue sample was digested in 100μL of 25mM NaOH/0.2mM EDTA lysis buffer at 98°C for 1 hour and vortexed prior to the addition of 100μL of 40mM Tris-HCl (pH 5.5). For each PCR reaction, 2μL of the crude lysate was added to GoTaq G2 Green master mix (Promega) and lacZ primers (see Supplemental Methods).

Zebrafish embryos were maintained in E3 medium (5 mM NaCl; 0.17 mM KCl; 0.33 mM CaCl2; 0.33 mM MgSO4) at 28.5°C. Morpholinos were dissolved in ultrapure water and injected into 1– to 2-cell stage zebrafish embryos alongside control morpholino as previously described (41). *Tnnt2* (ATG/start) blocking morpholino sequence was 5’ CATGTTTGCTCTGACTTGACACGCA 3′ as published in (74). Cessation of heartbeat was confirmed visually prior to analysis. To image, zebrafish embryos were dechorionated and anesthetized with 0.1% tricaine mesylate, GFP reporter gene expression screened with a Zeiss LSM 710 confocal microscope. Whole fish were imaged using the “tile scan” command, combined with Z-stack collection under a confocal microscope Zeiss LSM 710 MP (Carl Zeiss) at 488nm excitation and 509nm emission (EGFP). Embryos were embedded in 0.4% TopVision low melting point agarose (R0801, Thermo) in 0.14mg/ml Tricaine (Ethyl 3-aminobenzoate methanesulfonate, A5040, Sigma) in glass bottom multi-well culture plates (MatTek). Analysis of transgene expression was qualitative not quantitative, therefore no statistical analysis was applied to the observations of reporter gene intensity and pattern. No experimental randomization or blinding was used as we did not consider this necessary.

### RNAScope & Immunohistochemistry

Formalin-fixed, paraffin-embedded sections were analysed using the RNAscope® Multiplex Fluorescent Reagent Kit v2 (RNAScope, 323100) according to the manufacturer’s instructions. Briefly, slides were baked for 1 hour at 60°C, followed by three 5 minute incubations in Histoclear (deparaffinization) and two 2 minute incubations in 100% ethanol (dehydration). Slides were then treated with hydrogen peroxide (RNAScope, 322330) for 10 minutes and rinsed with distilled water. Afterwards, target retrieval was performed by incubating slides in 1× RNAScope Target Retrieval buffer (RNAscope, 322000) for 15 minutes in a steamer, followed by transfer to distilled water then 100% ethanol for 2 minutes. Slides were dried again at 60°C, treated with RNAScope protease plus (RNAScope, 322330) for 30 minutes at room temperature (RT), rinsed with distilled water then incubated with RNAscope® Probe Mm-Klf2-O1 (RNAScope, 510671) for 2 hours at 40°C. Probe amplification was performed by treating each section with the RNAScope Multiplex FL V2 Amp1 solution for 30 minutes at 40°C, repeated with Amp2 and Amp3 solutions. Probe signal was developed by incubation with RNAScope Multiplex FL V2 HRP-C1 for 15 min at 40°C, washing twice with 1x RNAScope Wash Buffer (RNAScope, 310091), followed by incubation with Opal 690 dye (1:1000, Akoya Biosciences, FP1497001KT) for 30 min at 40°C, washing, blocking with RNAScope Multiplex FL V2 HRP Blocker for 15 min at 40°C, and further washing. Immunohistochemistry was then performed to detect Endomucin protein expression. Briefly, the slides were washed twice in 1 x PBS and blocked with 4% FBS, 10% goat serum, 0.2% Triton X-100 in PBS for 1 hour. The sections were then incubated with primary anti-Endomucin antibody overnight at 4°C. The day after samples were re-washed and incubated with anti-rat Alexa 488 for 1 hour at RT. After re-washing, the slides were counterstained with DAPI (1:500,) and mounted. Sections were imaged using a Zeiss LSM 710 confocal microscope.

### Datasets used in enhancer identification

ATAC-seq in primary mouse adult aortic ECs (MAECs) (SRX7016285 GSM4128624) came from (75); ATAC-seq in ECs from adult mouse heart and lung (SRX14009316 GSM5856462; SRX14009317 GSM5856463), and EP300 ChIP-seq in ECs from adult mouse heart (SRX14009291 GSM5856486) came from (76); EP300 binding in Tie2Cre+ve cells from E11.5 mouse embryos (SRX2246377 GSM2346453) came from (77); ATAC-seq, H3K27Ac and ERG ChIP-seq in cultured human aortic ECs (HAECs) (SRX2355049 GSM2394391; SRX2355060 GSM2394402; SRX2355058 GSM2394400) came from (78); EP300 and ERG ChIP seq in human umbilical arterial ECs (HUAECs) (SRX5527609 GSM3673413; SRX5527654 GSM3673458) came from (79); and ETS1 ChIP-seq in HUVECs (SRX2452430 GSM2442778) came from (80).

### Tissue culture and flow shear stress application

An hTERT-immortalised aortic endothelial cell line (TeloHAEC, CRL4052) was acquired from ATCC and cultured in Vascular Cell Basal Medium (PCS-100-030) supplemented with Endothelial Cell Growth Kit-VEGF (PCS-100-041) at 37°C and 5% CO_2_. For FSS experiments, glass slides were coated with fibronectin before seeding cells. Confluent slides were mounted with a rubber gasket spacer on manifolds which allow medium to flow over cells via channels on either end of it. The manifolds were mounted into a custom parallel-plate flow chamber setup with a peristaltic pump and cells were exposed to shear stress at a rate of 15 dynes/cm^2^ for 6-12h (81, 82). Cells seeded on glass slides but kept under static conditions were used as control.

### CUT&RUN, ChIP-seq and RNA-seq

For RNA-seq, control cells and cells exposed to shear stress for 6h were harvested and RNA isolated using Direct-zol(TM) RNA MiniPrep kit (ZymoReseach, R2051). RNA quality was checked on a TapeStation (Agilent). RNA was converted to RNA-seq libraries using the Zymo-Seq RiboFree Total RNA Library Kit (ZymoResearch, R3000). Libraries were quality controlled using an Agilent Bioanalyzer with a High Sensitivity DNA Assay. Sequencing was performed on an Illumina NextSeq 2000 with paired-end reads of 61bp length. Data was analysed using the nf-core RNA-seq pipeline v3.12.0 (10.5281/zenodo.1400710) and DESeq2 package (83).

For CUT&RUN experiments, cells under static or FSS treatment were harvested with trypsin, washed, counted, and subsequently lightly cross-linked with 0.1% PFA (methanol-free, CST, 12606S) for 2 min. Crosslinking was stopped by adding glycine (CST, 7005S) and incubating for 5 min. Cells were pelleted and processed for CUT&RUN using the assay kit from Cell Signaling Technology (CST, 86652S). The following antibodies were used for CUT&RUN: IgG (CST, 66362S), TBP (CST, 12578S), KLF4 (R&D, AF3158). Buffers were modified to account for crosslinked samples by adding 1% Triton X-100 and 0.1% SDS to all steps before eluting the digested fragments. Crosslinking was reversed before purifying fragments by incubating samples at 65°C overnight in the presence of 0.1% SDS and 20 mg/ml proteinase K. Fragments were isolated with ChIP DNA Clean & Concentrator kit (ZymoResearch, D5205) and either used for qPCR or library preparation. CUT&RUN-qPCR was done on a Viia7 Real-Time PCR instrument (Applied Biosystems) using Fast SYBR Green (Life Technologies, #4385617), primer sequences available in Supplemental Methods. Conversion to libraries was done with the NEBNext® Ultra™ II DNA Library Prep kit for Illumina® (NEB, E7645L) and NEBNext® Multiplex Oligos for Illumina® (NEB, E7600S) following the protocol adapted for TFs (dx.doi.org/10.17504/protocols.io.bagaibse). Libraries were repeatedly cleaned as necessary with AmpureXP beads and libraries quality controlled on a TapeStation. Sequencing was performed by Azenta on an Illumina NovaSeq™ 6000.

GATA2 ChIP-seq in HUVECs (SRX070876 GSM730701, SRX150427 GSM935347) came from (84, 85); GATA4 in E12.5 heart ventricles (GSM1260024 SRX373594; GSM1260025 SRX373595) came from (86). RBPJ ChIP-seq in HUVECs after VEGFA stimulation (GSM2947453 SRX3599308) came from (49). TBP, RNA polymerase II (RNApolII), TFIIb and TFIIe ChIP-seq in mouse post-natal day 12 WT whole liver (GSM1390699 SRX547082; GSM1390701 SRX547084; GSM1390703 SRX547086; GSM1390705 SRX547088) came from (57), TBP ChIP-seq in WT testis (GSM2100637 SRX1667524) came from (87); and TBP ChIP-seq in WT cultured trophoblast stem cells (GSM967657 SRX160415) is from (88).

MEF2A and MEF2C ChIP-seq (tagmentation) and analysis used the protocol described in full in (76). In brief, the mouse heart and lungs were perfused, excised, minced, digested, mechanically disrupted, and filtered to obtain a single-cell suspension. CD31+/CD45– cells were isolated as ECs for downstream analyses. Flow-sorted ECs were fixed with formaldehyde, lysed, and frozen until use. Anti-Mef2A (Abcam, ab109420) and anti-Mef2C (Cell Signaling, 5030S) antibodies were conjugated to Protein A/G-coupled Dynabeads (Bimake, B23202) and incubated. Nuclear extract was sonicated, neutralized, and 5% of aliquots were saved for input controls. Antibody-coated Dynabeads were washed with PBS, mixed with nuclear lysate, incubated overnight at 4°C, and then washed (salt washes). For tagmentation, bead-bound chromatin was resuspended in 30 μL of tagmentation buffer, and one μL of transposase (Nextera, Illumina). The samples were incubated at 37°C for 10 minutes, followed by two washes with low-salt buffer. ChIP complexes were crosslinked, then diluted with ChIP elution buffer, vortexed, and incubated with proteinase K and RNase overnight at 65°C. Reverse crosslinked DNA purification was carried out using the Qiagen MinElute PCR Purification Kit. The libraries were amplified using 15 μL of PCR master mix and five μL of primer mix (Nextera, Illumina). Library clean-up was performed using AMPure XPeads (Beckman Coulter) at a 1:1 bead-to-sample ratio. Libraries were quantified and pooled. The libraries were single-end sequenced (50 cycles) on the NextSeq 500 platform (Illumina). All sequencing was performed with Psomagen Inc.

### Electrophoretic mobility shift assay

Recombinant ETS1-DNA binding domain (DBD), MEF2A, MEF2C, MEF2D, RBPJ, and GATA2 proteins for EMSA were *in vitro* transcribed and translated from pCITE2 (ETS1 DBD, MEF2C, and GATA2) or pcDNA3 (RBPJ, MEF2A, MEF2D) vectors using the TNT Quick Coupled Transcription/Translation System (Promega). Oligonucleotides (see Supplemental methods) were annealed, then labelled with ^32^P-dCTP using Klenow fragment (Promega) to fill in overhanging 5’ ends, and purified on a non-denaturing polyacrylamide-TBE gel (10%). Probe concentration was measured and diluted to 20-40000 counts per minute/μL. For the gel shift, binding reactions containing recombinant protein were pre-incubated at room temperature in 1x binding buffer (40 mM KCl, 15 mM HEPES pH 7.9, 1 mM EDTA, 0.5 mM DTT, 5% glycerol), 0.5-2 μL poly dI-dC and 100x excess competitor DNA where indicated for 10 min before the addition of radiolabelled probe. Reactions were incubated for an additional 25 min at room temperature after probe addition and electrophoresed on a 10% (ETS-DBD) or 6% (MEF2, RBPJ, GATA2) non-denaturing polyacrylamide gel for 1-2h. Gels were dried at 80°C and visualised by autoradiography.

### Bioinformatic analysis of MEF2-TBP motifs across genome

Motif analysis on the KLF2 promoter and enhancer sequences was performed using Homer’s findMotifsGenome.pl (89) tool with settings –size given –mask.

CUT&RUN data was processed with the nf-core cutandrun pipeline (v3.1 or 3.2.2) (10.5281/zenodo.5653535) using Bowtie2 alignment to hg38 genome, the hg38-CnR-blacklist.bed from Nordin *et al*., and CPM normalisation mode (90, 91). Peak calling was performed through the pipeline both with SEACR with stringent settings and MACS2(92, 93). Additionally, peaks were called with GoPeaks (94) and Homer (89). Homer peak calling was performed with setting –style factor and –size 200 and using BED files generated from BAM files with bedtools-bamtobed (95). Homer output was sorted with bedtools-sortBed, before overlap of the 4 peak callers was calculated with bedtools-multiinter. Only peaks recognised by at least 3 peak callers were considered for further analysis. Since some peaks were fragmented during overlap calculation due to varying peak sizes detected by the different peak callers, neighbouring peaks were merged with bedtools-merge to generate the final peak list per sample. Because some peaks are only present in static flow, while other sites are only bound by TBP under flow, the consensus peak list of TBP binding sites per experiment was generated by merging the peak list of the static and 6h flow samples for each experiment with bedtools-merge. The final high confidence TBP peak list was calculated by overlapping all three repeats using bedtools-multiinter and restricting the final high confidence peak list to peaks detected in at least 2 repeats.

For MEF2A peak analysis, previously published data SRR8335343, SRR8335344 and SRR8335345 (58) were downloaded from NIH Sequence Read Archive and processed with the nf-core ChIP-seq pipeline v2.1.0 (10.5281/zenodo.3240506) aligning to mm10 genome with Bowtie2. The pipeline calls peaks using MACS2 with narrow peak setting. Additionally, peaks were called with Homer as described above. Consensus peaks were calculated by overlapping the called peaks from both peak callers and both biological repeats and only considering peaks found in at least 3 of the 4 peak sets.

To identify the subset of consensus peaks containing motifs for both TBP and MEF2, the Homer tool annotatePeaks.pl was employed setting the motif option –m to the Homer motifs tata.motif and mef2a.motif or mef2b.motif or mef2c.motif or mef2d.motif (89). The outputs were combined and processed in R studio to filter peaks containing both a TBP and a MEF2 motif (96). The closest pair of TBP and MEF2 motifs and their distance in each peak containing both types of motifs was calculated and the final peak list annotated with Homer annotatePeak.pl, and in the case of the TBP peaks, also with the hg38 Genehancer.gtf annotation file in R studio (97). For comparison of motif instances in peaks found both in our TBP CUT&RUN and the previously published MEF2A ChIP-seq, MEF2A high confidence peaks were lifted over to hg38 genome coordinates using UCSC’s LiftOver tool before calculating overlapping peaks with the GenomicRanges package in R studio and annotating the peaks with Homer (98, 99).

The TBP-MEF2 double motif consensus was calculated from a sequence alignment containing the sequences of the KLF2 promoter, KLF2-41 and –66 enhancer, KLF4-148 enhancer and KLF6 and KLF7 promoter in human, mouse, opossum, chicken, xenopus and zebrafish using the tool Glam2 to calculate a gap-aware motif and alignment (100). The Glam2 motif alignment was shortened to remove edge bases with reduced conservation starting from the point where alignment was achieved by Glam2 introducing additional gaps (8 and 5 edge bases, respectively). The resulting shortened alignment of 26 nt was used as motif input for Glam2scan to search for additional instances of this double motif in open chromatin regions in ECs (100). These regions were identified by combining previously published ATAC-seq and p300 ChIP-seq in human and mouse ECs (SRX14009316-SRX14009317 (76), SRX7016285 (75), SRX14009291 (76), SRX5527609-10 (79)). Raw data was downloaded from the NIH Sequence Read Archive and processed with the nf-core ATAC-seq pipeline v2.1.2 (10.5281/zenodo.2634132) or ChIP-seq pipeline v2.1.0 (10.5281/zenodo.3240506) using Bowtie2 as aligner to either mm10 or hg38 genome. Since SRX14009291 did not have a matching input control it could not be processed with the ChIP-seq pipeline, but was instead processed with the ATAC-seq pipeline with the additional parameters –-mito_name false –-skip_merge_replicates. MACS2-called peaks were processed with bedtools as described above to calculate high confidence peaks considering true peaks as locations called in at least 3 data sets (mm10) or both data sets (hg38), respectively. The sequences corresponding to the high confidence peak locations were extracted using bedtools-getfasta and the corresponding fasta file used as input for the motif search with Glam2scan and setting flags –n 200 –2.

## Data Availability

All original CUT&RUN data has been uploaded onto GEO as GSE313645.

## Supporting information

All Supplemental Data

## Acknowledgements and funding sources

We thank Nadav Ahituv for providing the GW vectors. L.F., S.B., M.A.S., M.K.J., B.G.C and S.D., conceived and designed the study. L.F., V.V. M.K.J. and S.D. designed in vivo experiments, L.F., V.V. H.R-C., A.G., S.N., D.S., K.C., S.J.C, A.N., S.P. and S.D. implemented and analysed in vivo experiments, S.B. and B.G.C. designed the in vitro experiments, S.B., H.L., A.R., D.S.G, M.G. and B.G.C. implemented and analysed the in vitro experiments. L.F., S.B., and S.D. wrote the original manuscript, M.A.S., V.V., and B.G.C. edited the original manuscript, all authors provided editorial comments.

This work was supported by a Leducq Foundation Transatlantic Network of Excellence (18CVD03, to S.B., M.A.S., V.V., M.K.J., B.G.C. and S.D.), the British Heart Foundation (BHF) (FS/SBSRF/22/31037 to S.D.V. and S.N; FS/17/68/33478 to L.F.; PG/24/11910 to S.B.), the Oxford BHF Centre of Research Excellence (RE/18/3/34214 to A.G.), and the Medical Research Council (MR/S01019X/1 to H.R.C).

