## Supplementary material for "A transcriptional code controlling fluid shear stress-induced gene expression": All Supplemental Data

#### Supplemental methods

##### KLF enhancer and promoter sequences

**KLF2pr-0.4 human**

CAGGCGGGAGCCCGGGGCGCCCGGGGGATCCCTGAGCGTCACGCCGCTGTTGTGGAGCGCGTGTTGAC
AACGTCGCCGGGGAGACGGGCGGGGGCGGGGCCCGGGAGAGGGGGAGGCGCGGCCCTGGCGGCGCGCG
AGGGGCCGGGCTGTCAGCGCAAGGCCCAGGCCGCCGCAGTGGCCACGGCCGCTGCCGCCCGCCGGCTT
ATATACCGCGGCTAAATTTAGGCTGCGCCCGGAGCTCGTCCCCATCCGGGACGCGTTTCCGCCGCCGC
CGCTTTGGCCCGGCCCCCGCGCGCGCCGCGCCTATAAGGCTTGGGCGGGCCCGGCCGCGGCCCACAGA
GCCGTCCCCGCC

**KLF2-41 human**

GGTTGCACAACACTGCATGTATAAAATGCCACTACTTTAAAAATGGTTAATTTCATGTGAATTGCACT
TTAATAAAAATAAACAAGTATTTAAAAATCCACCAATGCAAACACTTCTCTGTTCCTCTGTGGTGGAG
GAAGGGACACGCGCTTTTTTTTTCCGACCTTAGGAAGGAACAAGGGAGCCGGGGTCCCCTCCCAGCCTG
GGAGCCCTGGGCACAGTCCCGGCTCATTTGTCAGAGCTATCGGAGCCGTCCTCGGGCTGGTGGGAGTT
CAGGGCTCTGAAAGGTTTTCTGTCAAGGCTTGAAAGGGGGCCAGGTTTTTTTCCCCCCGGAGCCGCGC
AGTCTCGGGGCTGTTGTTCTCAGCAATCGCAGGGCCTCGTGTTAGCAGGAAGCACAGCCAAGTAGGGT
TTCCTGCGTGTTGGAGAGAGGAAGCTCCGTAATGTTCTGGGAGGCGATGGTTAAAAATAACTCCGGTA
TATAAAGACAGCGGAGGGTCCCCTTGTTCGCTCACTCGGGCGCCGGCCGGCTGGACGCAGGGCCGAGC
AGGTGGTTTGGGGCCTCGGGAAGGCCAAACCCCCGCCTCTGGGCCCCTGGCTGGGGAAGACACCAGCC
AAGTTCAGAGCCCCAAGTCGGCCTCACTTCCACAACTCAGCGTCAGGGACACCGTGGGCGTTTCTGTT
TCAAAACGCTTTTCTCCAGCAAAGAACGTAACCTCAAGCTGCTGTCAGGGTAGAGGAATCCCTGCCCC
CCGCCGCCACCCAGATAGGTTGGGCCGTGATTTTGGCTGTGCCTGGCAGTGACGATGTAGCGCACCCA
GGCTGGGGGGCGCTGGCAGCCAGGGTTCAAATCCCGGCTCTGCCACCTAACGCTGGGACTCAGTTTC

**KLF2-66 human**

GATCGCAACATTGCACTCCAGCCTGGGCAAGAAGAGCAAAACTCCATCTCGAAATAAAAATAAAAATA
AAAAATAAATGAAATGGAGACCCCAGTACCCCCCGTTGATAGATAAGAGACTGAGATTTCTAGAGGGG
CTCAGCTTCCCTGGGGGACAGCTGTTTAGACCCAGAGCCAATTTTCAACTCTTGGGCAGCCAGACTGC
AAGTCCAGTGCTCCTTCCAAAACTTGTGTGTATGGAGAACCCACCACCTGTGCCCTGACCCCCTCCTG
GTTCCCCACAGCCCCTGCCCAGGCCAGGACACCCTGTCCCCTGGGTGTCCCTGAGGTCACAGATAAGG
CCGCTGGGCCCGGCTATCGGCAGGAAACATCTTGGCAGGACAGGAAGCGCGGCATGGGTCAGCCGCGC
CTGCACCCGGCTCAGCCTATCAGCGGCCAGATATTTTGGGAGCCACTATATAAACCCCAACTCGGCTC
CAAGCTGTTCCCCGGGGAAGCCTCAGGCTTCATGTTCTGCGGCAGCCCTGAGGCTGCTGGAACACGGG
GCCCGGGAGGAGCTCAGGAGCTGAGCGGGACAGAGTGACAGGGGTGGACTCCGTCCGGGCATCACTGC
AGGCCAGAGGGTGCTGGCTCTGAATCGAGCTAGCCCCTATTCAGTCCTGGTGCAGACACTCTCACTTG
CTGGGCAAGTGGCTTCACCGCCCTGTGCCTCAGTTTCCCCCTCCCCACATCCCTCCTTTGATCCTGTC
CCAGGGAAAAACACACCTCCAGGTCCTGGAGCCTTTTCTGAAGTTTCTGGGCCGACATCTGAGGGCCA
ACCCACTTTCTCCCAATGCAAAGCAGGGCCCAGGAATCTGCATCTTAAAACCTGCCTTGGCCGGGCAC
AGTGGCTCACACCTGTAATCCCAGCACTTTGGGAGGCTGAGGCGGGTGGATCATGAGGTCAGGCATTC
GAGACTAGCCTTG

**KLF2-34 human**

GCATGCTCTTATGCCTGGCTAATTTTTGTATTTTTTTGTAGAGATGAGCTCTTGCTATGTTG
CCCACCTGCTGGGACTCAATCCACCAAGTAGATGTCTCCAATTCTCCACCATTCCCCAAGAC
AAGTCTGGTGACAAAGAATTGGTTCAGAACCTGGGAAGTCTTTGAAGGTCCACTTTTGTCAG
ATGACACATGATCTAAATTGGTCCCCAACCCCAGGCCGCCGGAAAGGCTAGTGAGCATGGGG
TGAGCATAGAATCACCGCTCTATGAAAACATGCCTCCCTCAGCCCTGCCTCCTGGGCCACCG
AGGTCTGATTCACGCTTCCTCGTATTCAAACAGAGTCGGTGCAGCTGGAAGCAACCCTGCCA
GGGGTAAGTCAGCCAGTGCAGGTAACATTTCTCCCCCTCGATTTTTCCCAGAACCCGGCTCC
GGGCCCAACACAGGATGCCTGGGGGATGTCAGAGGATTCTAAGTGAGGAAACCCCTGCCGTC
CATTGCTGTTCTGTTCCCCCAGGGACGCTTTGTCCAGGGCCCACCGATCCATCACACATACA

AGGCCCAGGGCCCTGGAATGTTTAAGAGCCCACAAAAAAATAGCCTTTTTTTTTTTTTTGA
GACAGAGTCTCACTCTGTCGCCCAGGCTGGAGGGTAGTGGTGCAATCTTGGC

**KLF2-8 human**
CTCAGCTCCAAGCATTGCAGCGACAAGGTGGTGACCCGAGATAAGAAGCGCACTTGCAGAGG
GTGCCTCTGCCCGTATCTCTCCGCATCCTCAAGAAACACAGAACTATCAAACATTTTTGCAA
CATGAGGAGCCAAGTTTTGCAACATACTGAAAAAGGGTTGTGCAAACTTCAGCAGAAACACC
CCAATTGCCCCAGCTGGGAGTCATCATCCAGTTGCTGCTGTAATGTGATTGTAATTCCTCCA
GTGTTCATAATTTGTCTTGGAAGCTCAGCTGTATTTTACATGACGTGTTTTCTGTAGAATTC
CTGAGTACTTGCTCTTGGGATCTGAGCCAAGTGCACACAATAGCTATTTGGGGCAAAACAGC
TGGTAAAGAGAAAGGCAGCTTAGACCCTGAAGAAAAGTGGAAAGAATTCAAGGTTACTAGAA
ATGTCCAGGGCCCAGTAAAAGAAGATTGTAATCTCTCTATGTGCAGAAACAGGCCATATACT
TCATCAGTTTCCTCTCCATGCTCAGCTCAGCATGGAGCGGAAGCTCAGTATTTATTTGTGGC
TCAGACGAAAAACTTCTTGTTTGCAAGTTTCCTGAGCCCTTGAAGACAGCCTGGCAGGAGAC
AACAAATGCAAAGCTGCACACGTACCACTTATCCAATCCCCTGTTAGTGACAGACATCACCA
GTCAATCAGTCAATCACAGACATGACCAATCCATCACAGCACTGGGCCTGGCTACAACCTCA
GAATCTGCACCAGCACAGTGCTAAAAGCAACTATTATCAACAGATTTGAGGAAAATATTCTG
CCTCCTTGGTCTACATTTATAATGTTGAGTTGTTAGGAGAAGATAAGGGGCAGCAAGAAAGA
GATGACAATAAGATAATCACCATCGAGGCTG

**KLF4-148 human**
GGGCGAGTGCCCCGAGGGAGTCGGGCAAAAGCGAAATCCCGGCTTCCTACCCGGGACAGCGG
GAGAGCCGAGTTTTCCGCAGCGGCCGCGGGCTCGGAGCTTTTAAGGGTTTCTAATCCTGGAA
GGCTGTCTGACATCACCCCGTTTCTTGTCGGCTGATGTTTCGTACAAGCCTCTCATTTCCTC
AGTGTTTTCAGAGCCACCAATTACTGCAACAGTCTCGTGTTTATGTTTGCGCGCGCTCTCCT
GCCTCGCTGGCCCTCCGCCTGGGAGGCTGCGCTTCCCTCCGACGCGCGGAGCCCAGAGGGGG
TGGGTCGGGGGAGCGGGGAGGGGGGAATCTGGATCAGGCATTGAAGAAAAAGGGAGGGGTCC
CCCTTAATTTTTTTTCATTGACTTCAGCACCATGTGATCAGGAAGTCTGGCCTCCCTCCATT
TCCCCTCCCGACTAAAGGGAAACATTGTGTAGCAGCCGCCTGGGCCACCGGTGGGATGGCCT
TCGCTGCCTGACGTAGGGAGATAAAAATAACCGGCATATTTAAGGCAGGATCAGGAATCCCG
GCGCTCACACGCGGCCTGGTCAGTTCCCGAGGCCGCCCGGCAGGCAGCGCAGCCTGCGGGGA
GGCCCCGCGCTCGACTGTGCGCGCCCCACCGCAGGCAGTGCTAGGGCGTACGACCGGACCCC
GACCCTGACCCCGACCTTGACCTCGGGACCCAGGCCGACTCAGGCCTGGCTGAGCCTCCCAG
ATTCCCCAGGGAGGCCGAGCAAGACCGGCCCGAGGCAGTCTGGCTCAGGCTCGCTGGGCTGG
GGCTGGGCCTGGGTTCTCTGTCCGGCCTCCCGGGAGACGAGAAGGAAGGATGCCGTGGCAAA
GGGAGGGTAGTGCGAAGTTTGATGCCGAGGGGGGTTACAGAGCCTCTGTCTCAAGAGAAGGC
GCACAAGGTGCCACCAAAAAAGC

**KLF2-41mutMEF2**
GGTTGCACAACACTGCATGTATAAAATGCCACTACTTTAAAAATGGTTAATTTCATGTGAAT
TGCACTTTAATAAAAATAAACAAGTATTTAAAAATCCACCAATGCAAACACTTCTCTGTTCC
TCTGTGGTGGAGGAAGGGACACGCGCTTTTTTTTCCGACCTTAGGAAGGAACAAGGGAGCCG
GGGTCCCCTCCCAGCCTGGGAGCCCTGGGCACAGTCCCGGCTCATTTGTCAGAGCTATCGGA
GCCGTCCTCGGGCTGGTGGGAGTTCAGGGCTCTGAAAGGTTTTCTGTCAAGGCTTGAAAGGG
GGCCAGGTTTTTTTCCCCCCGGAGCCGCGCAGTCTCGGGGCTGTTGTTCTCAGCAATCGCAG
GGCCTCGTGTTAGCAGGAAGCACAGCCAAGTAGGGTTTCCTGCGTGTTGGAGAGAGGAAGCT
CCGTAATGTTCTGGGAGGCGATG*gtgcaaaac*GACTCCGGTATATAAAGACAGCGGAGGGTC
CCCTTGTTCGCTCACTCGGGCGCCGGCCGGCTGGACGCAGGGCCGAGCAGGTGGTTTGGGGC
CTCGGGAAGGCCAAACCCCCGCCTCTGGGCCCCTGGCTGGGGAAGACACCAGCCAAGTTCAG
AGCCCCAAGTCGGCCTCACTTCCACAACTCAGCGTCAGGGACACCGTGGGCGTTTCTGTTTC
AAAACGCTTTTCTCCAGCAAAGAACGTAACCTCAAGCTGCTGTCAGGGTAGAGGAATCCCTG

CCCCCCGCCGCCACCCAGATAGGTTGGGCCGTGATTTTGGCTGTGCCTGGCAGTGACGATGT
AGCGCACCCAGGCTGGGGGGCGCTGGCAGCCAGGGTTCAAATCCCGGCTCTGCCACCTAACG
CTGGGACTCAGTTTC

**KLF2-66mutMEF2**
TAAAAAATAAATGAAATGGAGACCCCAGTACCCCCCGTTGATAGATAAGAGACTGAGATTTCTAGAGG
GGCTCAGCTTCCCTGGGGGACAGCTGTTTAGACCCAGAGCCAATTTTCAACTCTTGGGCAGCCAGACT
GCAAGTCCAGTGCTCCTTCCAAAACTTGTGTGTATGGAGAACCCACCACCTGTGCCCTGACCCCCTCC
TGGTTCCCCACAGCCCCTGCCCAGGCCAGGACACCCTGTCCCCTGGGTGTCCCTGAGGTCACAGATAA
GGCCGCTGGGCCCGGCTATCGGCAGGAAACATCTTGGCAGGACAGGAAGCGCGGCATGGGTCAGCCGC
GCCTGCACCCGGCTCAGCCTATCAGCGGCCA*cc*T*tgg*TTGGGAGCCACTATATAAACCCCAACTCGGC
TCCAAGCTGTTCCCCGGGGAAGCCTCAGGCTTCATGTTCTGCGGCAGCCCTGAGGCTGCTGGAACACG
GGGCCCGGGAGGAGCTCAGGAGCTGAGCGGGACAGAGTGACAGGGGTGGACTCCGTCCGGGCATCACT
GCAGGCCAGAGGGTGCTGGCTCTGAATCGAGCTAGCCCCTATTCAGTCCTGGTGCAGACACTCTCACT
TGCTGGGCAAGTGGCTTCACCGCCCTGTGCCTCAGTTTCCCCCTCCCCACATCCCTCCTTTGATCCTG
TCCCAGGGAAAAACACACCTCCAGGTCCTGGAGCCTTTTCTGAAGTTTCTGGGCCGACATCTGAGGGC
CAACCCACTTTCTCCCAATGCAAAGCAGGGCCCAGGAATCTGCATCTTAAAACCTGCCTTGGCCGGGC
ACAGTGGCTCACACCTGTAATCCCAGCACTTTGGGAGGCTGAGGCGGGTGGATCATGAGGTCAGGCAT
TCGAGACTAGCCTTG

**KLF2-41mutRBPJ**
CACAACACTGCATGTATAAAATGCCACTACTTTAAAAATGGTTAATTTCAACCAAATTGCAC
TTTAATAAAAATAAACAAGTATTTAAAAATCCACCAATGCAAACACTTCTCTGTTCCTCTGT
GGTGGAGGAAGGGACACGCGCTTTTTTTTCCGACCTTAGGAAGGAACAAGGGAGCCGGGGTC
CCCTCCCAGCCTGGGAGCCCTGGGCACAGTCCCGGCTCATTTGTCAGAGCTATCGGAGCCGT
CCTCGGGCTGGTGGGAGTTCAGGGCTCTGAAAGGTTTTCTGTCAAGGCTTGAAAGGGGGCCA
GGTTTTTTTCCCCCCGGAGCCGCGCAGTCTCGGGGCTGTTG*tccttt*GCAATCGCAGGGCCT
CGTGTTAGCAGGAAGCACAGCCAAGTAGGGTTTCCTGCGTGTTGGAGAGAGGAAGCTCCGTA
ATGTTC*agaggg*GCGATGGTTAAAAATAACTCCGGTATATAAAGACAGCGGAGGGTCCCCTT
GTTCGCTCACTCGGGCGCCGGCCGGCTGGACGCAGGGCCGAGCAGGTGGTTTGGGGCCTCGG
GAAGGCCAAACCCCCGCCTCTGGGCCCCTGGCTGGGGAAGACACCAGCCAAGTTCAGAGCCC
CAAGTCGGCCTCACTTCCACAACTCAGCGTCAGGGACACCGTGGGCGTTTCTGTTTCAAAAC
GCTTTTCTCCAGCAAAGAACGTAACCTCAAGCTGCTGTCAGGGTAGAGGAATCCCTGCCCCC
CGCCGCCACCCAGATAGGTTGGGCCGTGATTTTGGCTGTGCCTGGCAGTGACGATGTAGCGC
ACCCAGGCTGGGGGGCGCTGGCAGCCAGGGTTCAAATCCCGGCTCTGCCACCTAACGCTGGG
ACTCAG

**KLF2-41mutETS**
CACAACACTGCATGTATAAAATGCCACTACTTTAAAAATGGTTAATTTCATGTGAATTGCAC
TTTAATAAAAATAAACAAGTATTTAAAAATCCACCAATGCAAACACTTCTCTGTTCCTCTGT
GGTGGAGGAAGGGACACGCGCTTTTTTTTCCGACCTTAGGAAGGAACAAGGGAGCCGGGGTC
CCCTCCCAGCCTGGGAGCCCTGGGCACAGTCCCGGCTCATTTGTCAGAGCTATCGGAGCCGT
CCTCGGGCTGGTGGGAGTTCAGGGCTCTGAAAGGTTTTCTGTCAAGGCTTGAAAGGGGGCCA
GGTTTTTTTCCCCCCGGAGCCGCGCAGTCTCGGGGCTGTTGTTCTCAGCAATCGCAGGGCCT
CGTGTTAGCA*gaga*GCACAGCCAAGTAGGG*tctc*CTGCGTGTTGGAGA*gaga*GAGCTCCGTA
ATGTTCTGGGAGGCGATGGTTAAAAATAACTCCGGTATATAAAGACAGCGGAGGGTCCCCTT
GTTCGCTCACTCGGGCGCCGGCCGGCTGGACGCAGGGCCGAGCAGGTGGTTTGGGGCCTCGG
GAAGGCCAAACCCCCGCCTCTGGGCCCCTGGCTGGGGAAGACACCAGCCAAGTTCAGAGCCC
CAAGTCGGCCTCACTTCCACAACTCAGCGTCAGGGACACCGTGGGCGTTTCTGTTTCAAAAC
GCTTTTCTCCAGCAAAGAACGTAACCTCAAGCTGCTGTCAGGGTAGAGGAATCCCTGCCCCC
CGCCGCCACCCAGATAGGTTGGGCCGTGATTTTGGCTGTGCCTGGCAGTGACGATGTAGCGC
ACCCAGGCTGGGGGGCGCTGGCAGCCAGGGTTCAAATCCCGGCTCTGCCACCTAACGCTGGG
ACTCAG

**KLF2-41mutTBP**
**GGTTG**CACAACACTGCATGTATAAAATGCCACTACTTTAAAAATGGTTAATTTCATGTGAAT
TGCACTTTAATAAAAATAAACAAGTATTTAAAAATCCACCAATGCAAACACTTCTCTGTTCC
TCTGTGGTGGAGGAAGGGACACGCGCTTTTTTTTCCGACCTTAGGAAGGAACAAGGGAGCCG
GGGTCCCCTCCCAGCCTGGGAGCCCTGGGCACAGTCCCGGCTCATTTGTCAGAGCTATCGGA
GCCGTCCTCGGGCTGGTGGGAGTTCAGGGCTCTGAAAGGTTTTCTGTCAAGGCTTGAAAGGG
GGCCAGGTTTTTTTCCCCCCGGAGCCGCGCAGTCTCGGGGCTGTTGTTCTCAGCAATCGCAG
GGCCTCGTGTTAGCAGGAAGCACAGCCAAGTAGGGTTTCCTGCGTGTTGGAGAGAGGAAGCT
CCGTAATGTTCTGGGAGGCGATGGTTAAAAATAACTCCGG*gcgcgccc*GACAGCGGAGGGTC
CCCTTGTTCGCTCACTCGGGCGCCGGCCGGCTGGACGCAGGGCCGAGCAGGTGGTTTGGGGC
CTCGGGAAGGCCAAACCCCCGCCTCTGGGCCCCTGGCTGGGGAAGACACCAGCCAAGTTCAG
AGCCCCAAGTCGGCCTCACTTCCACAACTCAGCGTCAGGGACACCGTGGGCGTTTCTGTTTC
AAAACGCTTTTCTCCAGCAAAGAACGTAACCTCAAGCTGCTGTCAGGGTAGAGGAATCCCTG
CCCCCCGCCGCCACCCAGATAGGTTGGGCCGTGATTTTGGCTGTGCCTGGCAGTGACGATGT
AGCGCACCCAGGCTGGGGGGCGCTGGCAGCCAGGGTTCAAATCCCGGCTCTGCCACCTAACG
CTGGGACTCAG**TTTC**

**KLF2-66mutTBP**
TAAAAAATAAATGAAATGGAGACCCCAGTACCCCCCGTTGATAGATAAGAGACTGAGATTTC
TAGAGGGGCTCAGCTTCCCTGGGGGACAGCTGTTTAGACCCAGAGCCAATTTTCAACTCTTG
GGCAGCCAGACTGCAAGTCCAGTGCTCCTTCCAAAACTTGTGTGTATGGAGAACCCACCACC
TGTGCCCTGACCCCCTCCTGGTTCCCCACAGCCCCTGCCCAGGCCAGGACACCCTGTCCCCT
GGGTGTCCCTGAGGTCACAGATAAGGCCGCTGGGCCCGGCTATCGGCAGGAAACATCTTGGC
AGGACAGGAAGCGCGGCATGGGTCAGCCGCGCCTGCACCCGGCTCAGCCTATCAGCGGCCAG
ATATTTTGGGAGCCAC*ctatcgcg*CCCCAACTCGGCTCCAAGCTGTTCCCCGGGGAAGCCTC
AGGCTTCATGTTCTGCGGCAGCCCTGAGGCTGCTGGAACACGGGGCCCGGGAGGAGCTCAGG
AGCTGAGCGGGACAGAGTGACAGGGGTGGACTCCGTCCGGGCATCACTGCAGGCCAGAGGGT
GCTGGCTCTGAATCGAGCTAGCCCCTATTCAGTCCTGGTGCAGACACTCTCACTTGCTGGGC
AAGTGGCTTCACCGCCCTGTGCCTCAGTTTCCCCCTCCCCACATCCCTCCTTTGATCCTGTC
CCAGGGAAAAACACACCTCCAGGTCCTGGAGCCTTTTCTGAAGTTTCTGGGCCGACATCTGA
GGGCCAACCCACTTTCTCCCAATGCAAAGCAGGGCCCAGGAATCTGCATCTTAAAACCTGCC
TTGGCCGGGCACAGTGGCTCACACCTGTAATCCCAGCACTTTGGGAGGCTGAGGCGGGTGGA
TCATGAGGTCAGGCATTCGAGACTAGCCTTG

**EMSA oligo sequences**

| Sequence name | Forward sequence | Reverse sequence |
|---|---|---|
| Rbpj_control site | ctagTTATTTTTGGCG**TGGGA**ACGCGGGGAGCAC | ctagGTGCTCCCCGCGT**TCCCA**CGCCAAAAATAA |
| Mef2_control site | ctagGATCCTCATCTT**TTAAAAATAA**CTTTTCAAAAG | ctagCTTTTGAAAAG**TTATTTTTAA**AAGATGAGGATC |
| Ets1_control site | ctagGGAGTTACTCTC**TTCC**TGTTATG | ctagTGTCATAACA**GGAA**GAGAGTAA |
| Gata2_control site | ctagGCTAGGT**TTATCA**CTGCCTC | ctagGAGGCAG**TGATAA**ACCTAGC |
| KLF2-41 Rbpj_A WT | ctagGGCTGTTGT**TCTCA**GCAATCGCAG | ctagCTGCGATTGC**TGAGA**ACAACAGCC |
| KLF2-41 Rbpj_A MT | ctagGGCTGTTGT**TATAG**GCAATCGCAG | ctagCTGCGATTGC**CTATA**ACAACAGCC |
| KLF2-41 Rbpj_B WT | ctagCGTAATGTTC**TGGGA**GGCGATGGT | ctagACCATCGCC**TCCCA**GAACATTACG |
| KLF2-41 Rbpj_B MT | ctagCGTAATGTTC**CAGTA**GGCGATGGT | ctagACCATCGCC**TACTG**GAACATTACG |
| KLF2-41 Mef2 WT | ctagGGCGATGG**TTAAAAATAA**CTCCGGT | ctagACCGGAG**TTATTTTTAA**CCATCGCC |
| KLF2-41 Mef2 MT | ctagGGCGATGG**TGCAAAACGA**CTCCGGT | ctagACCGGAG**TCGTTTTGCA**CCATCGCC |
| KLF2-41 ATmot WT | ctagTAACTCCG**GTATATAAAG**ACAGCGGA | ctagTCCGCTTGT**CTTTATATAC**CGGAGTTA |
| KLF2-41 ATmot MT | ctagTAACTCCG**GCCGCACACA**CAAGCGGA | ctagTCCGCTTGT**GTGTGCGGC**CGGAGTTA |
| KLF2-41 ETS_A WT | ctagAGGTTTTT**TTCC**CCCCGGAG | ctagCTCCGGGG**GGAA**AAAAACCT |

| | | |
|---|---|---|
| KLF2-41 ETS_A MT | ctagAGGTTTTT**GGAC**CCCCGGAG | ctagCTCCGGGG**GTCC**AAAAACCT |
| KLF2-41 ETS_B WT | ctagTGTTAGCA**GGAA**GCACAGCC | ctagGGCTGTGC**TTCC**TGCTAACA |
| KLF2-41 ETS_B MT | ctagTGTTAGCA**TTCA**GCACAGCC | ctagGGCTGTGC**TGAA**TGCTAACA |
| KLF2-41 ETS_C WT | ctagAGTAGGGT**TTCC**TGCGTGTT | ctagAACACGCA**GGAA**ACCCTACT |
| KLF2-41 ETS_C MT | ctagAGTAGGGT**GGAC**TGCGTGTT | ctagAACACGCA**GTCC**ACCCTACT |
| KLF2-41 ETS_D WT | ctagTGGAGAGA**GGAA**GCTCCGTA | ctagTACGGAGC**TTCC**TCTCTCCA |
| KLF2-41 ETS_D MT | ctagTGGAGAGA**TTCA**GCTCCGTA | ctagTACGGAGC**TGAA**TCTCTCCA |
| KLF2-66 RBPJ_AB WT | ctagCCACGTG**TCCCT**GAGG**TCACA**GA | ctagTC**TGTGA**CCTC**AGGGA**CACGTGG |
| KLF2-66 RBPJ_AB MT | ctagCCACGTG**GAAAG**GAGG**TCACA**GA | ctagTC**TGTGA**CCTC**CTTTC**CACGTGG |
| KLF2-66 RBPJ_C WT | ctagATATTT**TGGGA**GCCACTA | ctagTAGTGGC**TCCCA**AAATAT |
| KLF2-66 RBPJ_C MT | ctagATATTT**CAAGA**GCCACTA | ctagTAGTGGC**TAAAC**AAAATAT |
| KLF2-66 MEF2 WT | ctagCGGCCAG**ATATTTTGGG**AGGC | ctagGCTC**CCCAAAATAT**CTGGCCG |
| KLF2-66 MEF2 MT | ctagCGGCCAG**ATAGCTTTT**GGGCGGC | ctagGCCGCAG**AAAAGCTAT**CTGGCCG |
| KLF2-66 TAmot WT | ctagAGCCA**CTATATAAAC**CCCAA | ctagTTGGG**GTTTATATAG**TGGCT |
| KLF2-66 TAmot MT | ctagAGCCA**CGAGAGAAAC**CCCAA | ctagTTGGG**GTTTCTCTCG**TGGCT |
| KLF2-66 ETS_A WT | ctagATCGGCA**GGAA**ACATCTTGGC | ctagGCCAAGATGT**TTCC**TGCCGAT |
| KLF2-66 ETS_A MT | ctagATCGGCA**GGGG**ACATCTTGGC | ctagGCCAAGATGT**CCCC**TGCCGAT |
| KLF2-66 ETS_B WT | ctagAGGACA**GGAA**GCGCGG | ctagCCGCGC**TTCC**TGTCCT |
| KLF2-66 ETS_B MT | ctagAGGACA**GCCA**GCGCGG | ctagCCGCGC**TGGC**TGTCCT |
| KLF2-66 GATA_A WT | ctagGAGGTCAC**AGATAA**GGCCGCTG | ctagCAGCGGCC**TTATCT**GTGACCTC |
| KLF2-66 GATA_A MT | ctagGAGGTCAC**TTTCCC**GGCCGCTG | ctagCAGCGGCC**GGGAAA**GTGACCTC |
| KLF2-66 GATA_B WT | ctagCAGCGGCC**AGATAT**TTTGGGAG | ctagCTCCCAAA**ATATCT**GGCCGCTG |
| KLF2-66 GATA_B MT | ctagCAGCGGCC**TTTCCC**TTTGGGAG | ctagCTCCCAAA**GGGAAA**GGCCGCTG |
| KLF2-66 GATA_C WT | ctagGGCCCG**GCTATC**GGCAGGAA | ctagTTCCTGC**CGATAG**CCGGGCC |
| KLF2-66 GATA_C MT | ctagGGCCCG**GGAGAG**AGCAGGAA | ctagTTCCTGC**TCTCTC**CCGGGCC |
| KLF2-66 GATA_D WT | ctagGCTCAGC**CTATCA**GCGGCCAG | ctagCTGGCCGC**TGATAG**GCTGAGC |
| KLF2-66 GATA_D MT | ctagGCTCAGC**CACACA**GCGGCCAG | ctagCTGGCCGC**TGTGTG**GCTGAGC |
| KLF2-66 GATA_E WT | ctagCCCCCCGT**TGATAGATAA**GAGAC | ctagGTCTCTTAT**CTATCA**ACGGGGGG |
| KLF2-66 GATA_E MT | ctagCCCCCCGT**TGAGAGACAA**AGAGAC | ctagGTCTCTTTGT**CTCTCA**ACGGGGGG |
| KLF2-0.4 MEF2 WT | ctagCGCAGC**CTAAATTTAG**CCGCG | ctagCGCAGC**CTAAATTTAG**CCGCG |
| KLF2-0.4 MEF2 MT | ctagCGCAGC**CAAAAAAAAG**CCGCG | ctagCGCGG**CTTTTTTTTG**GCTGCG |
| KLF2-0.4 TAmot WT | ctagCCGCG**GTATATAAGC**CGGCGG | ctagCCGCCG**GCTTATATAC**CGCGG |
| KLF2-0.4 TAmot MT | ctagCCGCG**GTTTTTTTGC**CGGCGG | ctagCCGCCG**GCAAAAAAAC**CGCGG |

**qPCR primers**

GAPDH_promoter_ChIP_fw TGGTGTCAGGTTATGCTGGGCCAG;
GAPDH_promoter_ChIP_rv GTGGGATGGGAGGGTGCTGAACACGCG[102]; KLF2-0.4 qPCR_fw CCGCCCGCCGGCTTAT;
KLF2-0.4 qPCR_rv AAACGCGTCCCGGATGG;
KLF2-41 qPCR_fw TAGCAGGAAGCACAGCCAAG;
KLF2-41 qPCR_rv TAACCATCGCCTCCCAGAAC;
KLF2-66 qPCR_fw GAGCCACTATATAAACCCCAAC;
KLF2-66 qPCR_rv GCAGAACATGAAGCCTGAG;
KLF4-148 qPCR_fw CTCCCTCCATTTCCCCTC;
KLF4-148 qPCR_rv GCGAAGGCCATCCCAC.

#### Figure S1

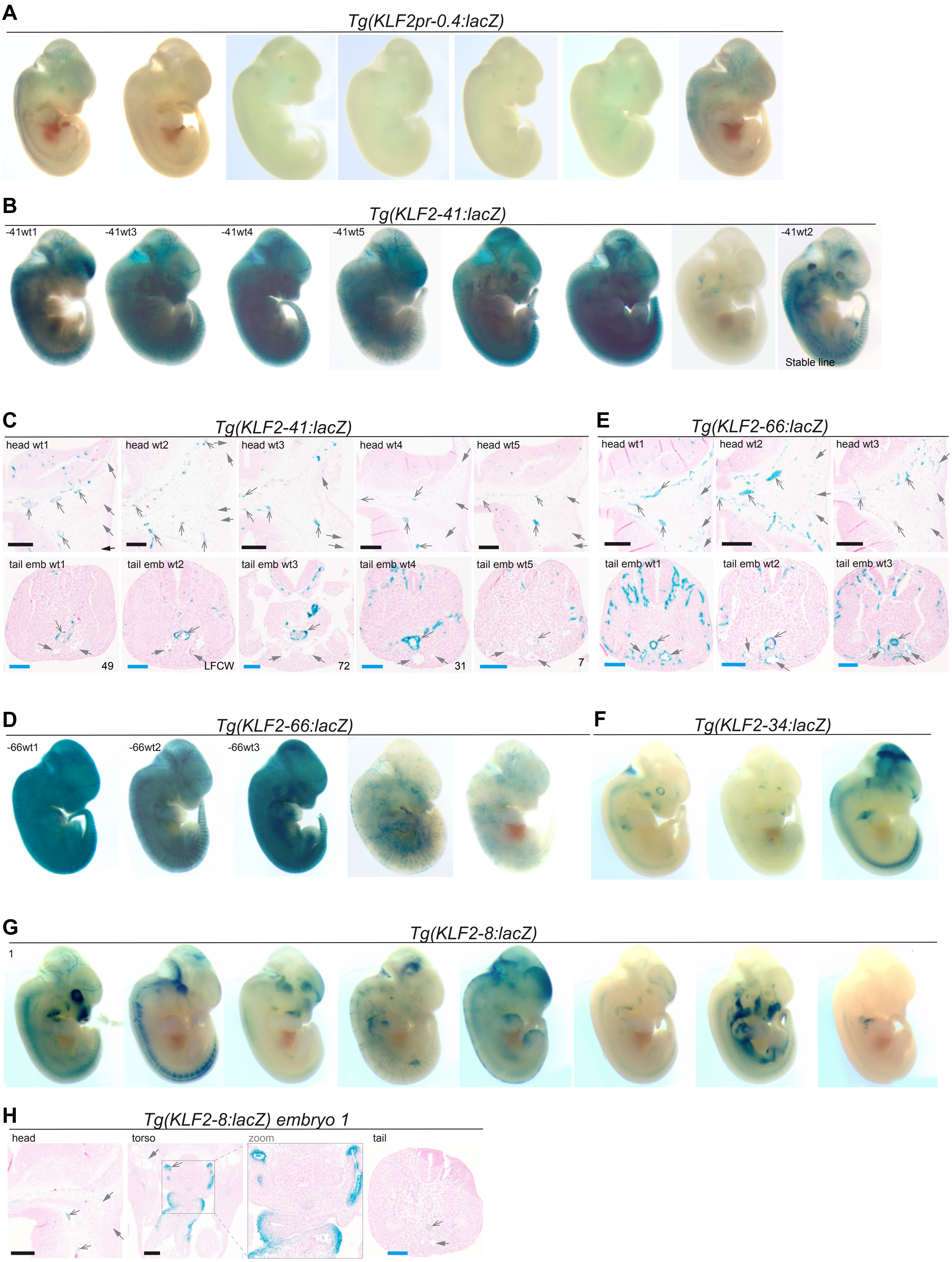
A
Tg(KLF2pr-0.4:lacZ)
B
Tg(KLF2-41:lacZ)
-41wt1
-41wt3
-41wt4
-41wt5
-41wt2
Stable line
C
Tg(KLF2-41:lacZ)
head wt1
head wt2
head wt3
head wt4
head wt5
tail emb wt1
tail emb wt2
tail emb wt3
tail emb wt4
tail emb wt5
49
LFCW
72
31
7
E
Tg(KLF2-66:lacZ)
head wt1
head wt2
head wt3
tail emb wt1
tail emb wt2
tail emb wt3
D
Tg(KLF2-66:lacZ)
-66wt1
-66wt2
-66wt3
F
Tg(KLF2-34:lacZ)
G
Tg(KLF2-8:lacZ)
1
H
Tg(KLF2-8:lacZ) embryo 1
head
torso
zoom
tail

**Supplemental Figure S1**

**F0 transgenic embryos expressing putative enhancers within the *KLF2* locus**
**A** Wholemount images of all E12.5 F0 embryos transgenic for the *KLF2pr-0.4:lacZ* transgene.
**B-C** Wholemount images (**B**) and transverse sections through (**C**) all X-gal positive E12.5 F0 embryos transgenic for the *KLF2-41:hsp68:lacZ* transgene.
**D-E** Wholemount images (**D**) and transverse sections through (**E**) all X-gal positive E12.5 F0 embryos transgenic for the *KLF2-66:hsp68:lacZ* transgene.
**F** Wholemount images of all X-gal positive E12.5 F0 embryos transgenic for the *KLF2-34:hsp68:lacZ* transgene.
**G-H** Wholemount images of all X-gal positive E12.5 F0 embryos transgenic for the *KLF2-8:hsp68:lacZ* transgene (**G**), and transverse sections through (**H**) a *KLF2-8:lacZ* expressing embryo.
Black scale bar is 200µm, blue scale bar is 100µm, grey open arrow is artery, grey closed arrow is vein.

#### Figure S2

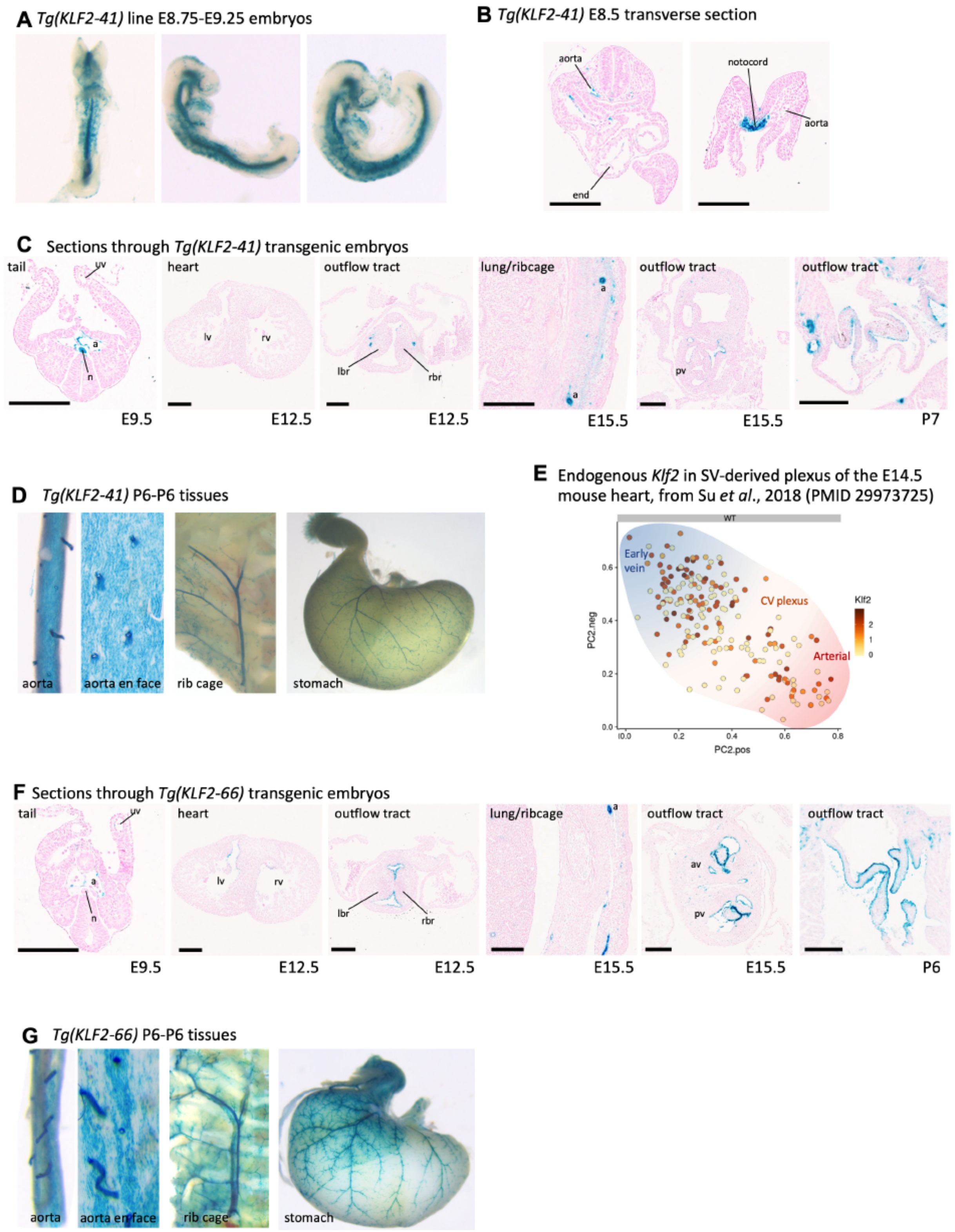
A Tg(KLF2-41) line E8.75-E9.25 embryos
B Tg(KLF2-41) E8.5 transverse section
aorta
end
notocord
aorta
C Sections through Tg(KLF2-41) transgenic embryos
tail
uv
a
n
heart
lv
rv
outflow tract
lbr
rbr
lung/ribcage
a
a
outflow tract
pv
outflow tract
E9.5
E12.5
E12.5
E15.5
E15.5
P7
D Tg(KLF2-41) P6-P6 tissues
aorta
aorta en face
rib cage
stomach
E Endogenous Klf2 in SV-derived plexus of the E14.5 mouse heart, from Su et al., 2018 (PMID 29973725)
WT
Early vein
CV plexus
Arterial
Klf2
2
1
0
PC2.neg
PC2.pos
F Sections through Tg(KLF2-66) transgenic embryos
tail
uv
a
n
heart
lv
rv
outflow tract
lbr
rbr
lung/ribcage
a
outflow tract
av
pv
outflow tract
E9.5
E12.5
E12.5
E15.5
E15.5
P6
G Tg(KLF2-66) P6-P6 tissues
aorta
aorta en face
rib cage
stomach

**Supplemental Figure S2**

**Further images detailing reporter gene expression throughout development in stable *KLF2-41:lacZ* and *KLF2-66:lacZ* transgenic mice lines, in comparison to *Klf2* expression.**

**A-B** Representative wholemount stable *KLF2-41:lacZ* transgenic embryos (A) and transverse sections (B) at E8.75-9.25, showing expression in arterial ECs and notochord.

**C** More representative sections through stable *KLF2-41:lacZ* transgenic embryos.

**D** Expression patterns in stable *KLF2-41:lacZ* pups.

**E** Expression of endogenous *Klf2* is seen throughout the SV-derived plexus at E14.5 (data from [37])

**F** More representative sections through stable *KLF2-66:lacZ* transgenic embryos.

**G** Expression patterns in stable *KLF2-66:lacZ* pups.

Black scale bars represent 200μm. a = aorta, uv = umbilical vein, n = notochord, l/rbr = left/right bulbar ridge, av = aortic valve, pv = pulmonary valve.

#### Figure S3

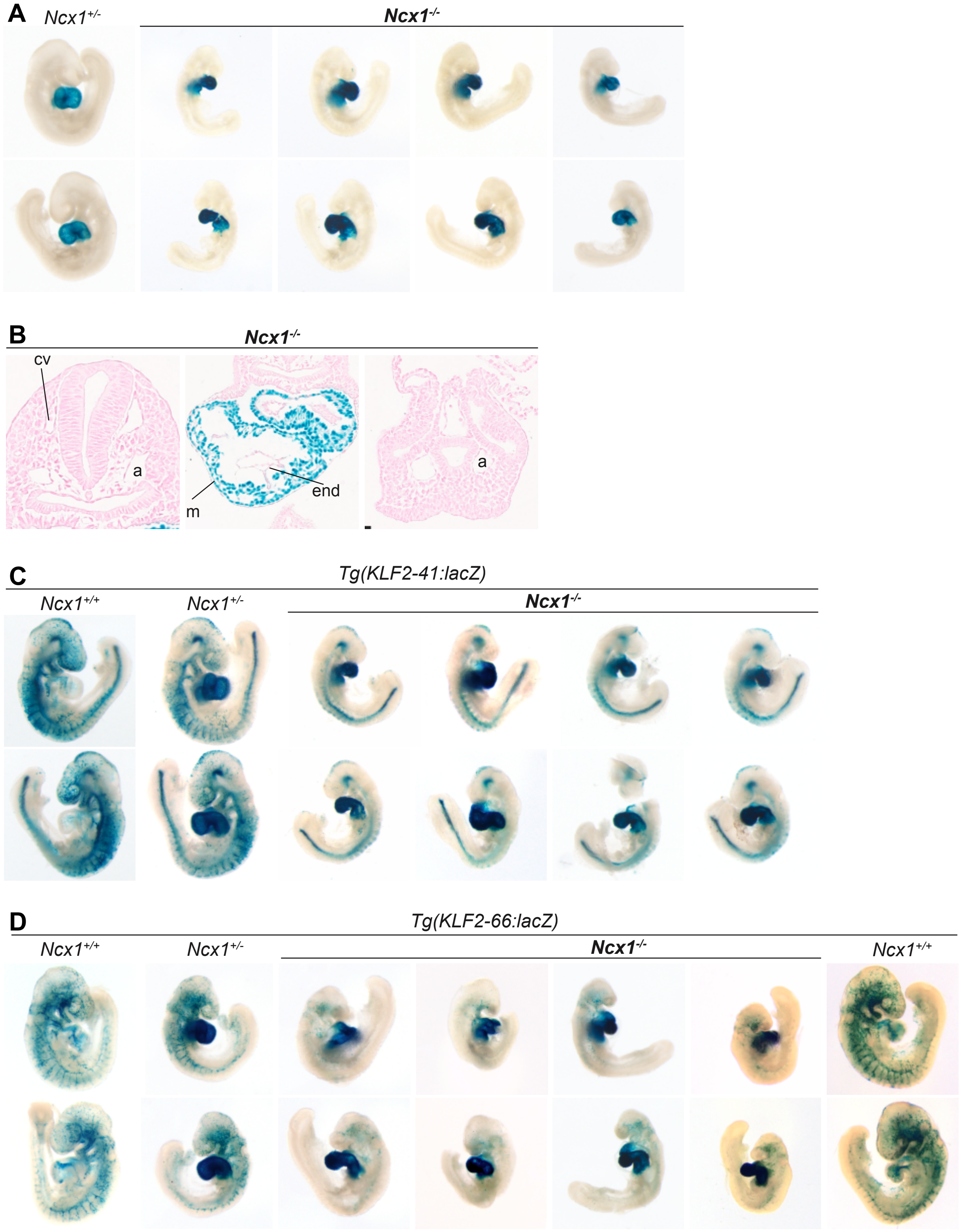
A
Ncx1+/-
Ncx1-/-
B
Ncx1-/-
cv
a
m
end
a
C
Tg(KLF2-41:lacZ)
Ncx1+/+
Ncx1+/-
Ncx1-/-
D
Tg(KLF2-66:lacZ)
Ncx1+/+
Ncx1+/-
Ncx1-/-
Ncx1+/+

**Supplemental Figure S3**

**More images of *enhancer:lacZ* transgenes on the *Ncx1*$^{-/-}$ background**

**A-B** Wholemount E9.25 *Ncx1*$^{+/-}$ and *Ncx1*$^{-/-}$ embryos (**A**) and transverse sections through *Ncx1*$^{-/-}$ embryos (**B**) with no enhancer:lacZ transgene included. **C-D** All generated embryos expressing the *KLF2-41:lacZ* (**C**) and *KLF2-66:lacZ* (**D**) transgenes on a *Ncx1*$^{-/-}$ background compared to representative WT and *Ncx1*$^{+/-}$ controls. Embryos in D on far right were left in stain for 4 days, control included for comparison.

Upper and lower images are images of same embryo from different aspects. a = aorta, cv = cardinal vein, m = myocardium, end = endocardium.

#### Figure S4

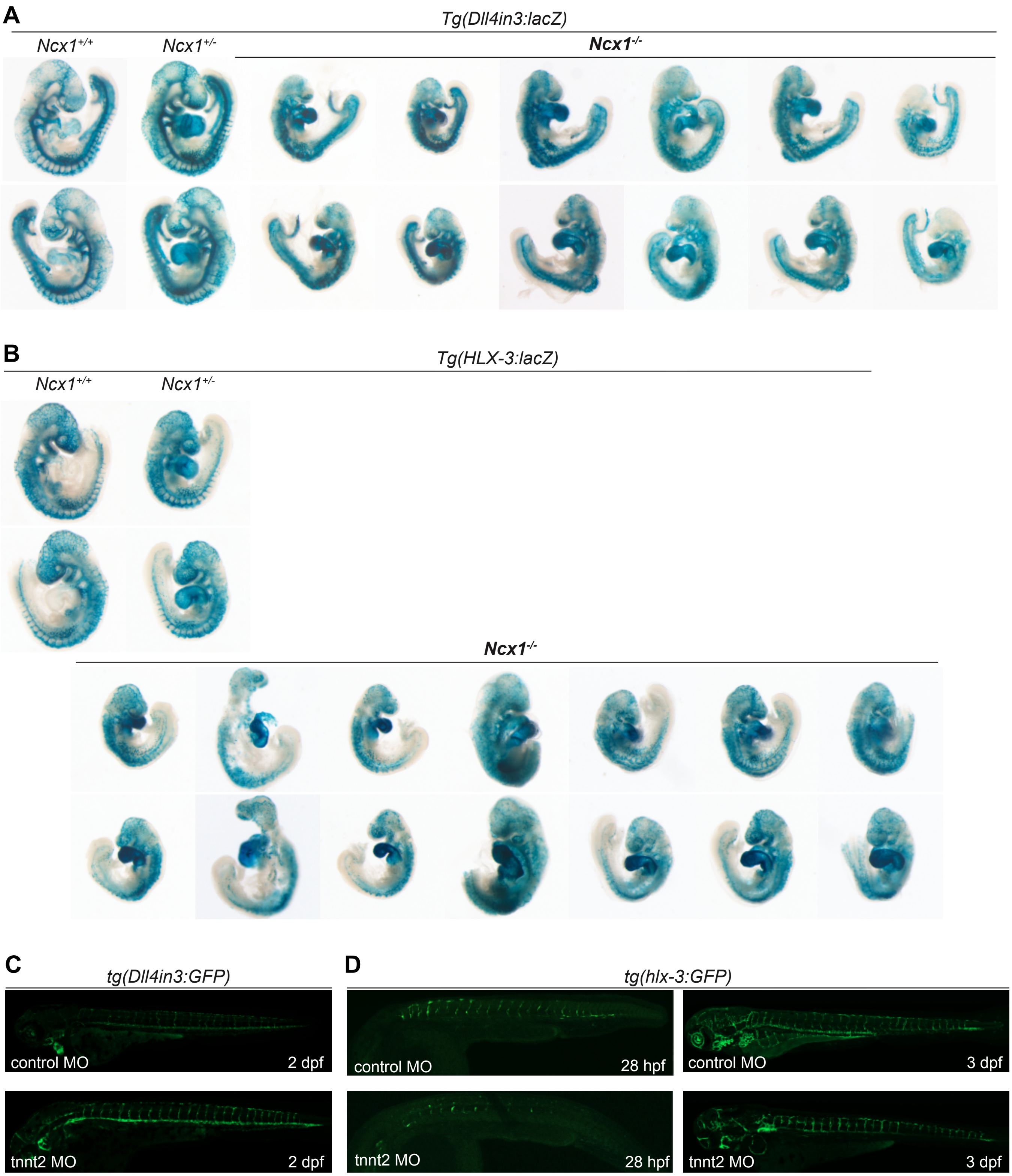
A
Tg(Dll4in3:lacZ)
Ncx1+/+
Ncx1+/-
Ncx1-/-
B
Tg(HLX-3:lacZ)
Ncx1+/+
Ncx1+/-
Ncx1-/-
C
tg(Dll4in3:GFP)
control MO
2 dpf
tnnt2 MO
2 dpf
D
tg(hlx-3:GFP)
control MO
28 hpf
tnnt2 MO
28 hpf
control MO
3 dpf
tnnt2 MO
3 dpf

**Supplemental Figure S4**

**Dll4in3 and HLX-3 enhancer activity is not reduced after loss of flow in both zebrafish and mouse models.**

**A-B** All generated E9.25 embryos expressing the *Dll4in3:lacZ* (**A**) and *HLX-3:lacZ* (**B**) transgenes on a $Ncx1^{-/-}$ background compared to representative WT and $Ncx1^{+/-}$ controls. Activity is not impacted by loss of heartbeat. Upper and lower images are images of same embryo from different aspects.

**C-D** Expression of the *Dll4in3:GFP* (**C**) and *hlx-3:GFP* (**D**) transgenes in zebrafish is unaffected by ablation of flow by the *tnnt2* morpholino (MO) when compared to control morpholino.

### Figure S5

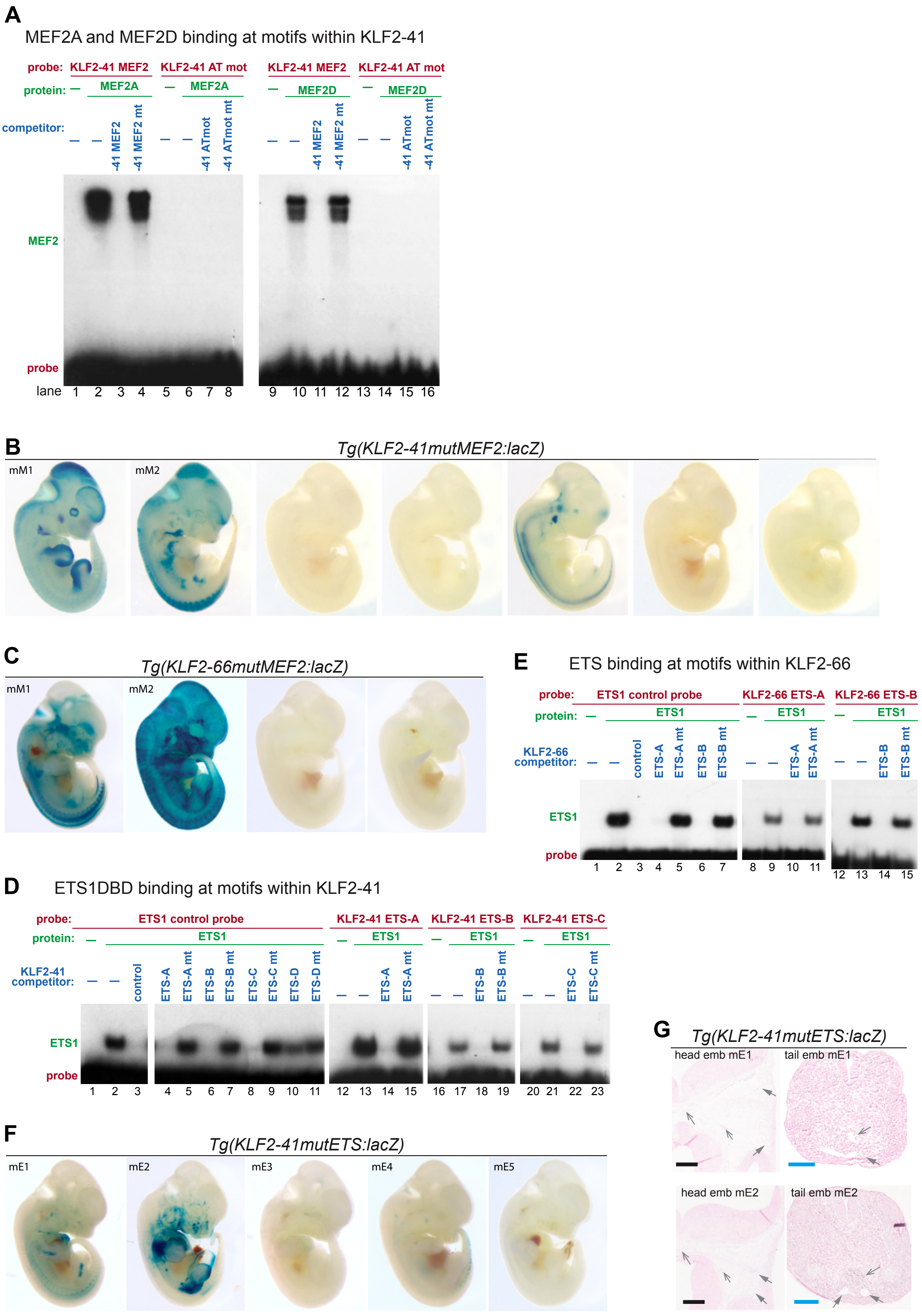
A
MEF2A and MEF2D binding at motifs within KLF2-41
probe: KLF2-41 MEF2 KLF2-41 AT mot KLF2-41 MEF2 KLF2-41 AT mot
protein: — MEF2A — MEF2A — MEF2D — MEF2D
competitor: — — -41 MEF2 -41 MEF2 mt — — -41 ATmot -41 ATmot mt — — -41 MEF2 -41 MEF2 mt — — -41 ATmot -41 ATmot mt
MEF2
probe
lane 1 2 3 4 5 6 7 8 9 10 11 12 13 14 15 16
B
Tg(KLF2-41mutMEF2:lacZ)
mM1
mM2
C
Tg(KLF2-66mutMEF2:lacZ)
mM1
mM2
E
ETS binding at motifs within KLF2-66
probe: ETS1 control probe KLF2-66 ETS-A KLF2-66 ETS-B
protein: — ETS1 — ETS1 — ETS1
KLF2-66 competitor: — — control ETS-A ETS-A mt ETS-B ETS-B mt — — ETS-A ETS-A mt — — ETS-B ETS-B mt
ETS1
probe
1 2 3 4 5 6 7 8 9 10 11 12 13 14 15
D
ETS1DBD binding at motifs within KLF2-41
probe: ETS1 control probe KLF2-41 ETS-A KLF2-41 ETS-B KLF2-41 ETS-C
protein: — ETS1 — ETS1 — ETS1 — ETS1
KLF2-41 competitor: — — control ETS-A ETS-A mt ETS-B ETS-B mt ETS-C ETS-C mt ETS-D ETS-D mt — — ETS-A ETS-A mt — — ETS-B ETS-B mt — — ETS-C ETS-C mt
ETS1
probe
1 2 3 4 5 6 7 8 9 10 11 12 13 14 15 16 17 18 19 20 21 22 23
F
Tg(KLF2-41mutETS:lacZ)
mE1
mE2
mE3
mE4
mE5
G
Tg(KLF2-41mutETS:lacZ)
head emb mE1
tail emb mE1
head emb mE2
tail emb mE2

**Supplemental Figure S5**

**Consequences of mutating the MEF2 and ETS motifs within the *KLF2* enhancers.**

**A** EMSA analysis showing direct binding of the MEF2 motif within the *KLF2-41* enhancer to MEF2A (lanes 1-4) and MEF2D (lanes 9-12) demonstrating that MEF2A, C and D bind with similar affinity. The neighbouring AT-rich motif (ATmot) within the *KLF2-41* enhancer bound neither MEF2A (lanes 5-8) nor MEF2D (lanes 13-16).

**B-C** Images of all E12.5 embryos transgenic for the *KLF2-41mutMEF2:lacZ* (**B**) and *KLF2-66mutMEF:lacZ* **(C)** transgenes.

**D-E** EMSA analysis of ETS motifs within the *KLF2-41* (**D**) and *KLF2-66* (**E**) enhancers showing competition for (left) and direct binding of (right) ETS1 DNA binding domain (ETS1-DBD). In **D**, *KLF2-41* ETS motifs A, B and C can compete for ETS binding of a control probe (controls lanes 1-3, *KLF2-41* sites lanes 4-9), and can also directly bind the ETS1 DBD protein (lanes 12-15, 16-19 and 20-23). In **E**, *KLF2-66* ETS motifs A and B compete for ETS binding of a control probe (controls lanes 1-3, *KLF2-66* sites lanes 4-7), and also directly bind the ETS1 DBD protein (lanes 8-11 and 12-15).

**F-G** Images of all E12.5 embryos (**F**) and transverse sections of transgenic embryos **(G)** for the *KLF2-41mutETS:lacZ* transgenes. Mutation of ETS motifs ablates all EC expression of this transgene.

Black scale bar is 200µm, blue scale bar is 100µm, grey open arrow is artery, grey closed arrow is vein.

### Figure S6

**A** GATA2 binding at motifs within KLF2-66

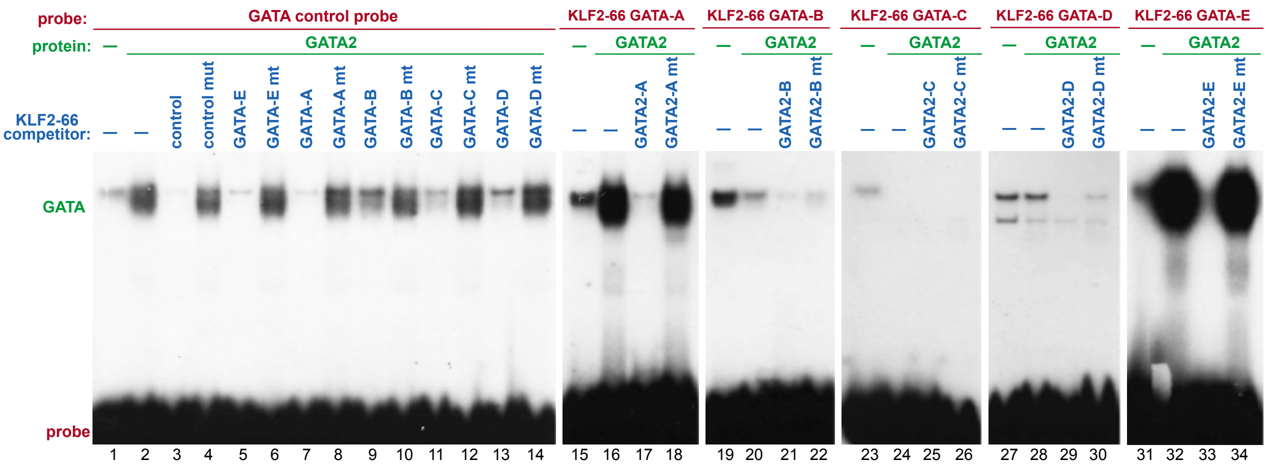


**B**

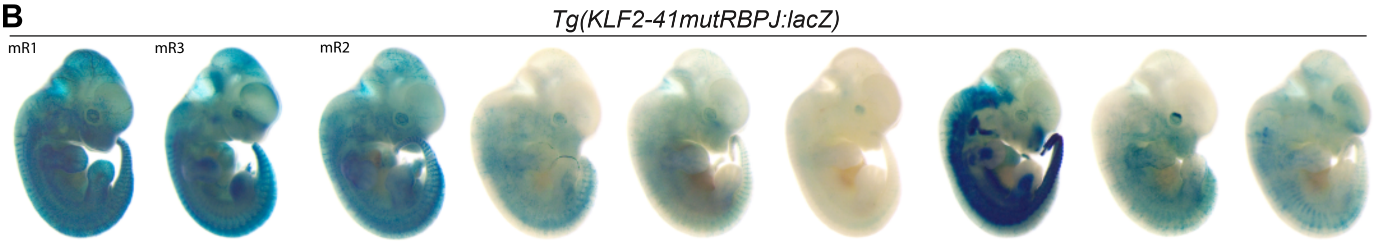


**C**

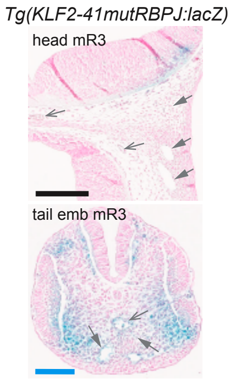


**D** RBPJ binding at motifs within KLF2-41

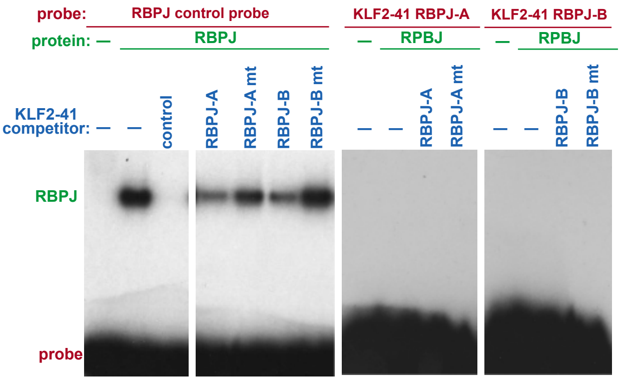


**E**

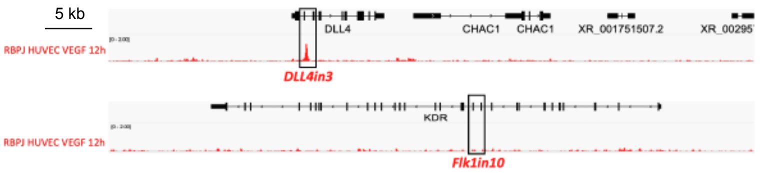

**Supplemental Figure S6**
**A** EMSA analysis showing competition by (left) and direct binding of GATA2 protein to (right) putative GATA motifs within the *KLF2-41* and *KLF2-66* enhancers. Only GATA motifs A and E within the *KLF2-66* enhancer were able to either compete for binding of GATA2 (lanes 5-8) or directly bind GATA2 (lanes 16-18 and 32-34).
**B-C** Images of all E12.5 wholemount embryos (**B**) and sections through one embryo (**C**) transgenic for the *KLF2-41mutRBPJ:lacZ*.
**D** EMSA analysis showing competition by (left) and direct binding of RBPJ protein to (right) putative RBPJ motifs within the *KLF2-41* enhancer. Motifs within the *KLF2-41* enhancer were able to compete for binding of RBPJ (lanes 4-8) but none could directly bind RBPJ.
**E** RBPJ ChIP-seq in HUVECS after VEGFA stimulation shows binding at the angiogenic *Dll4in3* but not at the arterial Flk1in10 enhancer.

### Figure S7

**A** *tg(ETS1+195:GFP)*

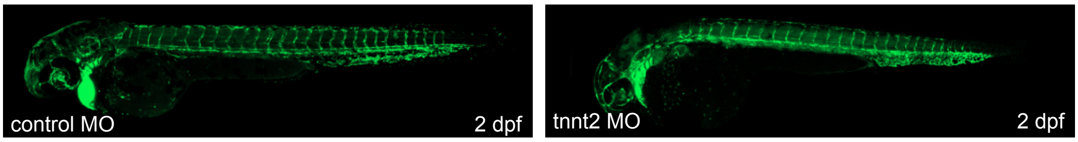


**B** TRANSFAC annotation of MEF2 and TBP motifs

*KLF2-66* *KLF2-41* *KLF2-promoter*

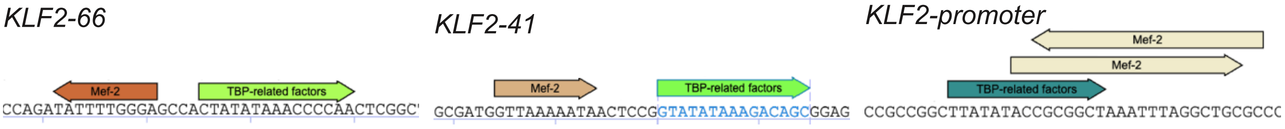


**C**

```
                                              SYTTATATASYGnnnRTAWWWWTAR
                                                   TBP         MEF2          TBP
                                                          RGHTAWWWWTARMnnRSVTATWTAARS
KLF2-41                   ETS
human     TCCTGCGTGTTGGAGAGAGGAAGCTCCGTAATGTTCTGGGAGGCG-ATGGTTAAAAATAACTCCGGTATATAAAGACAGCGGAGGG-TCCCCTTGTTCGCTCA----CTCGGGCGCC-----GGC
mouse     TCCTGCGCACTGGGGAGAGGAAACTCCGTAATGTACTGAGAGCCG-A-AGTTAAAAATAACCCCAGTATATAAAGAGCGCAGAGGG-TCCCCT-GCTCCTTCAG-AGCTCGGCTGTCAGGCTGGA
opossum   TCCTGAGCGTCTGAGAGAGGAAACACTGTAATAAGCATGGAGGCC-AGGGTTAAAAATAACTCGGGTATATAAAGGCTGCCTCAGAATCCCATCTCGTCTCAAA-CTCTCACTCG-CACACTTTC
chicken   TCCTGAGCGTCTGAGAGAGGAAGCGCTGTAATAAGCATGGAGGCT-GTGGTTAAAAATAACCCCGGTATATAAAGGCCGGCAG-GA-TCCCCTCTCTCACTCAC-ACACAGCGCTGCCTCTTGGC
xenopus   TCCTGAATGTCTGAGAGAGGAAATCTTACAAGCAGTGCGCAATCC-GAGGATAAAAATAACTGCCCTATATAAGAGCAGCTGAGGG-TCCCCGTCCACTCTCAC-A-TGCACTGCTCCTGCCTCT
zebrafish TCCTGAGCGTCCGAGAGAGGAAGCCCTGTGATAGGCGAGAGGGAGTGGGGTTAAAAATAGT-GGCGTATATAAAGGCT-CGGGTTT-TCCCTGTTCGGCCTCATTCTCTCTCCTGCTGCTCTCAC
          *****      * ********     *                 * ********       *******        *      ****

KLF2-66                               GATA
human     CA-GCCGCGCCTGC-CCCGG----------CTCAGCCTATCAGCGGCCAGATATTTTGGGAGCCACTATATAAACCCCAACTCGGCTCCAAGCTGCCG-GGCTCAGCTTCCAGCCGCTCCCGCG-
mouse     CA-ACTGCGCCCGCGCCGAG----------CTCAGCCTATCCTTGGCCTGATATTTTGGGAGCCGCTATTTAAGCCCGAACTCGACTCCTGCCATCCCCGACTGGTCTGCGAGTTGTTCCCGGG-
pig       CA-GCCGCGCCCTCACCCGC----------CTCAGCCTATCCGCGGCCAGATATTTTGGGAGGCACTATATAAACCCCGACTGGTCTGCGAGTTGCCC-GGCTCAGCCCGCAGC-ACTCCCGGG-
manatee   CCCGCCGCGCCCGCGCCAGC----------CTTGGCCTATCGGCGGCCAGATATTTTGGGAGCCGCTATATAAACCGGGCCTCAGCTTCCAGCCGCCCCAACTCGGCTCCAAGCTGTTCCCCGG-
tenrec    CTGGCCTCGCCCGCACCAGC----------CTCGGCCTATCCAGGGCCAGATATTTTGGGAGGCACTATATAAGCCCGGCCTCAGCCCGCAGC-ACCC-GACTCGACTCCTGCCATGAGCTGGA-
tas dev   TGGCCCGGGCCAGAAGCAGGACCACGGCTGCTCGGCCTATCAGCGGGCAGATATTTTGGGAGCTGCTATATAAGCTCCTCCCAAGCACACAGTGTCTC-CTCCAAGCACACAGTGTCTCTTGGAC
              *   ***     *          **  *******   ** * *************   **** *** *      *      *        *

KLF2-pr
human     TG---GGCCGCGGCCGGGCCCGCCCAA--GGACGAGCTCCGGG-CGCAGCCTAAATTTAGCCGCGGTATATAAG----CCG-GCGGGCGGCAGCGGCCGTGGCCA--CTGCGGCGGCCTGGGCCC
mouse     TG---TGCTGTGGCCCGGCCCACCCAG--GGACGAGCTCCGGG-CTCAGCCTAAATTTAGCCGCGGTATATAAG----CCT----GGCGGTGGTGGCCG--GCCG--CTGAGGCCGCGCGGGCCG
tenrec    CG---GCCGCCGG--GGTCCCGCCCA---GGACGAGCTCCGGGGCGCAGCCTAAATTTAGCCGCAGTATATAAG----CCGTGCGGGCAGCGGCGGCTGTGGCGG--CTACTGCGG---GAGCCC
opossum   AGT--GGCGGCCCCGGGGCCCGCCCGA--GGACGAGCTCTGGG-CTCAGCCTAAATTTAGCCGCAGTATATAAG----CCG--GCGGCGGCCGCGGCTGCAGTGG--CTGCGG—ACCAGAGCC-
chicken   CGGGACGCGGCGCGCAGAGCCGCCCTCCCAGGCAGCGGCTCGGGCGCAGCCTAAATTTAGCCCCGGTATAAAAGGACGCGGCGGCGGCGGCCGCAGCCCGAGCGGGGCGGAGCGGAGCGGGGCCC
xenopus   TTCCTTGCAGCTGAGAGGGG-GCCTG---GGCTGAGCACTCAG-TGGAGCCTAAATTTAGCAGCACTATATAAA----CCCTGCAGTCAAGTTCAGTCT--GCAG---CGAGAGCAGCAGCTCCG
                *       *    **     *       *  *   **************  *  **** **     *       * *      *
```

**D** Source of MEF2 motifs within known MEF2-bound promoters and enhancers

| Element | Source paper | location on hg38 |
|---|---|---|
| CKM (MCK) enhancer | PMID: 2761536 | chr19:45,323,399-45,323,878 |
| Myogenin Myf4 promoter | PMID: 8006037 | chr1:203,085,952-203,086,329 |
| Desmin enhancer | PMID: 8626009 | chr2:219,417,206-219,417,709 |
| Slc2a4 (Glut4) promoter | PMID: 7545962 | chr17:7,281,026-7,281,499 |
| MHC promoter | PMID: 8366095 | mm10 chr3:154,752,994-154,753,224 |
| Tnni3 promoter | PMID: 9738004 | chr19:55,157,727-55,157,860 |
| MYL2 promoter | PMID: 10207035 | chr12:110,920,505-110,920,764 |
| Bdnf promoter | PMID: 22973001 | chr11:27,701,590-27,701,929 |
| Nur77 promoter | PMID: 10944115 | chr12:52,050,962-52,051,351 |
| Jun promoter | PMID: 10403485 | chr1:58,783,980-58,784,299 |
| ACTC1 enhancer | PMID: 15491989 | chr15:34,796,794-34,797,087 |
| Myoglobin Mb promoter | PMID: 7891680 | chr22:35,617,304-35,617,663 |
| MYH4 promoter | PMID: 9614136 | chr17:10,469,510-10,469,882 |
| MYL1 enhancer | PMID: 2243772 | chr2:210,288,312-210,288,608 |

**E**

| | *KLF2-41* | *KLF2-66* | ***KLF2-41 mutTBP*** | ***KLF2-66 mutTBP*** |
|---|---|---|---|---|
| *Tg mice with EC expression* | 6 | 5 (1*) | 4 (2*) | 0 |
| *Tg mice with non-EC expression* | 1 | 0 | 6 | 3 |
| *Tg mice with no expression* | 3 | 4 | 1 | 1 |
| ***Total number of tg mice*** | 10 | 9 | **11** | **4** |

**F** *Tg(KLF2-41**mutTBP**:lacZ)*

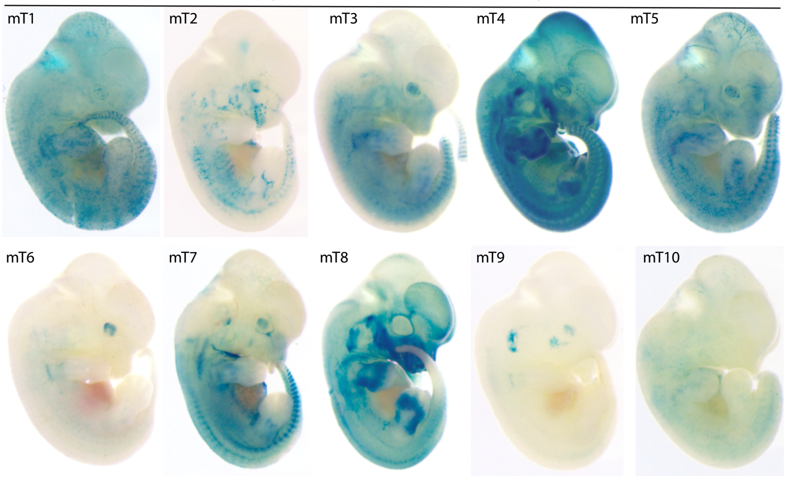


**G** *Tg(KLF2-66**mutTBP**:lacZ)*

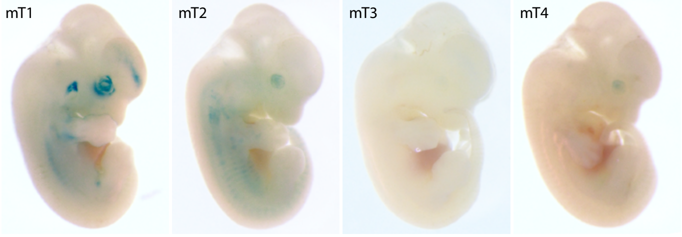


**H** *Tg(KLF2-41**mutTBP**:lacZ)*

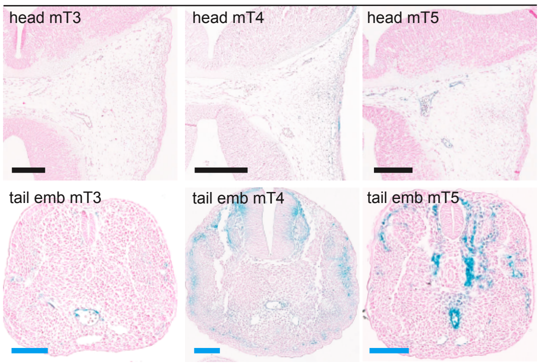

#### Supplemental Figure S7

**Identification and analysis of the MEF2-TBP double motif shared across *KLF2* elements**

**A** The MEF2-bound *ETS1+195:GFP* angiogenic transgene is unaffected by loss of blood flow (via tnnt2 MO injection) in transgenic zebrafish.

**B** TRANSFAC analysis of the *KLF2-66*, *KLF2-41* and *KLF2* promoter also identifies motifs for MEF2 and TBP alongside each other.

**C** The MEF2-TBP double motif within the *KLF2-41*, *KLF2-66* and *KLF2* promoter is deeply conserved across species, with the only notable change being an alteration of spacer from 3 bp to 2 bp in the zebrafish version of *KLF2-41*. For each element, sequences from six species were selected to represent different stages of evolution, with the most related to most distant listed in order top to bottom.

**D** References used to identify MEF2-bound non-EC enhancers and promoters. Although these all contained MEF2 motifs, none had equivalent TBP motifs.

**E** Summary of transgenic numbers for embryos expressing KLF2-41 and KLF2-66 enhancers with TBP mutants.

**F-H** Wholemount images of all embryos (**F, G**) and additional transverse sections of transgenic embryos (**H**) for the *KLF2-41mutTBP:lacZ* (F, H) and the *KLF2-66mutTBP:lacZ* transgene, all embryos E12.5. See also Figure 6.

#### Figure S8

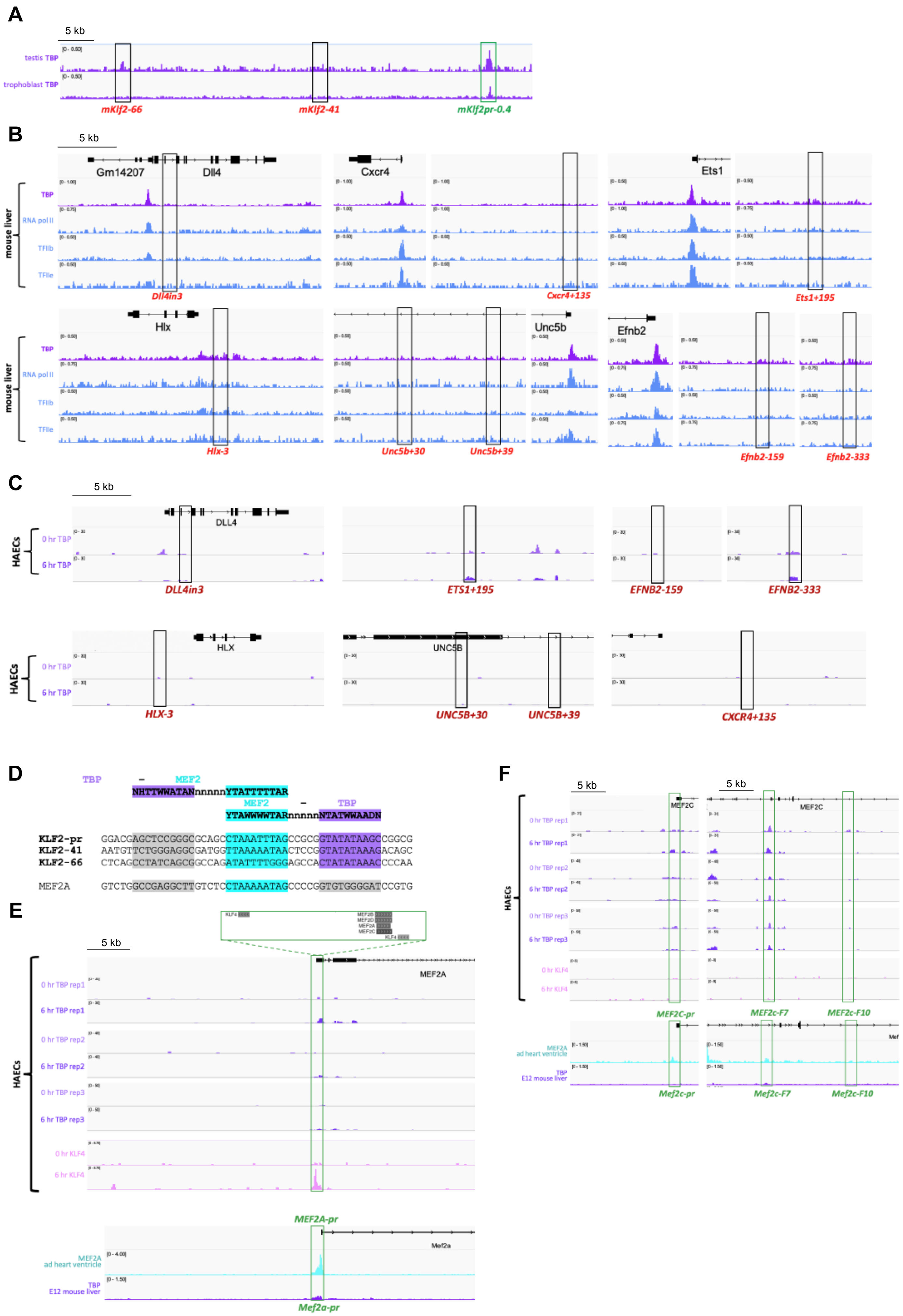
A
5 kb
testis TBP
trophoblast TBP
mKlf2-66
mKlf2-41
mKlf2pr-0.4
B
5 kb
mouse liver
TBP
RNA pol II
TFIIb
TFIIe
Gm14207
Dll4
Cxcr4
Ets1
Dll4in3
Cxcr4+135
Ets1+195
Hlx
Unc5b
Efnb2
Hlx-3
Unc5b+30
Unc5b+39
Efnb2-159
Efnb2-333
C
5 kb
HAECs
0 hr TBP
6 hr TBP
DLL4
DLL4in3
ETS1+195
EFNB2-159
EFNB2-333
HLX
UNC5B
HLX-3
UNC5B+30
UNC5B+39
CXCR4+135
D
TBP – MEF2
NHTTWWATANnnnnnYTATTTTTAR
MEF2 – TBP
YTAWWWWTARnnnnnNTATWWAADN
KLF2-pr GGACGAGCTCCGGGCGCAGCCTAAATTTAGCCGCGGTATATAAGCCGGCG
KLF2-41 AATGTTCTGGGAGGCGATGGTTAAAAATAACTCCGGTATATAAAGACAGC
KLF2-66 CTCAGCCTATCAGCGGCCAGATATTTTGGGAGCCACTATATAAACCCCAA
MEF2A GTCTGGCCGAGGCTTGTCTCCTAAAAATAGCCCCGGTGTGGGGATCCGTG
E
KLF4
MEF2B
MEF2D
MEF2A
MEF2C
KLF4
5 kb
MEF2A
HAECs
0 hr TBP rep1
6 hr TBP rep1
0 hr TBP rep2
6 hr TBP rep2
0 hr TBP rep3
6 hr TBP rep3
0 hr KLF4
6 hr KLF4
MEF2A-pr
Mef2a
MEF2A ad heart ventricle
TBP E12 mouse liver
Mef2a-pr
F
5 kb
5 kb
MEF2C
MEF2C
HAECs
0 hr TBP rep1
6 hr TBP rep1
0 hr TBP rep2
6 hr TBP rep2
0 hr TBP rep3
6 hr TBP rep3
0 hr KLF4
6 hr KLF4
MEF2C-pr
MEF2c-F7
MEF2c-F10
MEF2A ad heart ventricle
TBP E12 mouse liver
Mef2c-pr
Mef2c-F7
Mef2c-F10

**Supplemental Figure S8**

**TBP binding across the genome**

**A** TBP binding at the *KLF2* enhancers was not seen in ChIP-seq from tissues without significant EC portion.

**B** TBP and RNA polII binding in liver was detected at many EC gene promoters, but little TBP binding was seen at non-flow responsive enhancers with MEF2 motifs but without MEF2-TBP motifs

**C** Increased TBP binding in telo-HAECs was not seen at non-flow responsive enhancers with MEF2 motifs but without MEF2-TBP motifs

**D-E** The *MEF2A* promoter contains a MEF2 motif but not TBP motif (**E**), and does not bind TBP (**D**). However, it contains multiple KLF motifs and binds KLF4 after FSS, binding which increases after 12 hours of FSS.

**F** The MEF2C promoter and enhancers (F7 and F10)[101] do not bind TBP or KLF4, in keeping with the lack of increased transcription seen after FSS (see Figure 8).

### Figure S9

**A**

```
                                     SYTTATATASYGnnnRTAWWWWTAR
                                         TBP          MEF2           TBP
                                                  RGHTAWWWWTARMnnRSVTATWTAARS
KLF6-pr
human     AATTCGAAC------TCTGCTCGT-TCTGATTGGAAAGTGGCCTCGCAGGCTATTTTTAGC-AGGGTATATAAGGCGCGGGCGGCCGCGCG--CCGGAGGAGTCCGCGCTGTGAGGT------GC
mouse     AATTCGAAC------TCGGCCTGC-GCTGATTGGAAAGACGTCTCACAGGCTATTTTTAGC-AGGGTATATAAGGCGAGGGCTACGGCGCA--CCGAAGGAGTCGGCGCTGGGAGGT------GT
opossum   AATTCGAAA------TTTCCCCAAATCTGATTGGTGGGCGGCCGTGCAGGCTATTTTTAGC-AGGGTATATAGGGCGCGAGCTCCCGGGCAG-CCCGAGGAGTTGGTGCGGTGTCGTTTCGCTGT
chicken   AATTCGAAA------TTTCCCCAAATCTGATTGGTGGGCGGCCGTGCAGGCTATTTTTAGC-AGGGTATATAGGGCGCGAGCTCCCGGGCAG-CCCGAGGAGTTGGTGCGGTGTCGTTTCGCTGT
xenopus   GATTCGAAAATTCAGCCCCCTCCCCTCGGGTAGCTGGTGGGACGGACCAGCTATTTTTAGA-TGGGTATATAAGGCAAGGACTACACGGCGGTTCGGTACAGTGCGGTCTGTGGCCGCTCGGAGC
zebrafish AATTCAA---------CTGGACGGGCCTGCT-GCGCGCTGATTGGAC--GGTATTTTTAGA-AACATATATAAAGGCTCTGCGCTCAAACCTCTCAGTTCTGTTT-CTTGACAACGGAGTGTGGC
           **** *                **** *          * * *********   ******  *                *    *      **
KLF7-pr
human     TGGCACGCTGCGGATCTCCCTGCTGCCTTACAAGGCGGTGCGAG-GCGAGCTATTTTTAGA-GGGCTATATAAGGG--------GGCTGAGGCGGCCG-CCCGGCGGCCGGGCAGCGCGA-----
mouse     TGGCGCGCTGCGGATCTCCCTGCTGCCTTACAAGGCGGTACGAG-GCAAGCTATTTTTAGA-GGGCTATATAAAGG--------GGTAGAGGAGTCCG-CCAGGCGGCCGGGGAGCTTGA-----
opossum   TGGAGCAAGCCTGATCTCCCTTCTGCCTTACAAGGCGGTGTAAG-GCGAGCTATTTTTAGA-GAGCTATATAAGGG----GCCAGAGTCGGCCGGCCG-TGCGGAGGCCTGGGAGAGGGA-----
chicken   TGGTCCGATTCGAAATTCGCCCCCTCCTGATTGGCGAGCTCGGCTGCGGGCTATTTTTAGC-AGGGTATATAAAGCAGGGAAGCGGCTGCTTTTT-CTGGTAGGGTGGTGTGTGGTTGTGTGCTG
xenopus   TGGC-TGCTGGGTCACTCTCTGCTGCCTTACAAGGCAGTGCAAG-TAGAGCTATTTTTAGA-GGCCTATATAAGCT-------GGAATGAGGCG--CG-TACAGTTGCCTGTGAGAGAGAT----
zebrafish TGGCTCGTGACACTTGTCAATCGGGCCGTAGAAGGCAGTGAAAG-GTGAGCTATTTTTAGA-GAGCTATATAGCAG--CGGCA-GCGCACTGGCGTCGCTCATACGGCCGGAGAGAACGCA----
          ***          **     **  *   *                ***********   *******           *         *        *
KLF4-148                                                                                 ETS
human     -GCCTGGGCCACCG---GTGGGATGGCCTTCGCTG-CCTGACGTAGGGAGATAAAAATAACCGGCATATTTAAGGCAGGAATCCCGGCGCTCACAC-GCG-GCCTGGTCAGTT-CCCGAGGCCGC
mouse     TGCCGCGGCCACTG---GTGGGATGGCCTTCGCTGGCCTGACGTAGGGGGATAAAAATAACCGGCATATTTAAGGCCGGAATCCCGGCGCTCACAC-GCG-GCCTGGTCAGTT-CCCGACGCCGC
opossum   -GCTTCTGCCACTG---GTGGGATGGCCTTTCGCAGCATTACGTAGGAAGATAAAAATAACCGGCATATTTAAGGCCGGCATCCCAGCGCAGATCCAGTGCGTCTGGTCAGTTCACGGAGCACAG
chicken   -GCC-CTGCCACCG---GTGGGACGGCCCCGGGCCGCATTACGTAGGAGGATAAAAATAACCGGGATATTTAAGGCCGGCATCCCAGCGCAGGCAGAGCGAGCCTGGTCAGCT-GCCGAGCGCGC
xenopus   ---------GTCTG---GTAGGCAGGTCTTTTGAGATGCTACGTAGGAGGATAAAAATAACCAGCATATTTAAAGGCTGCATCCCAGCACAGGCA--GAGAGGCAGGTCAGCCATTGAGGCATAA
zebrafish TGTGTGTGTGTGTGTGTGTGGGCGGTACTGGGCTGCTCTTACATGCAGGGTTAAAAATAACCAGCATATTTAAGGGC-GAATCCCTTCAATCCCAGAGGCAGCGCAGACTGCC---TCGCGACAG
                     *    ** **   *  *            ** *      * *********** * ******** *   * *****  *          *   *     * * *
```

**B**

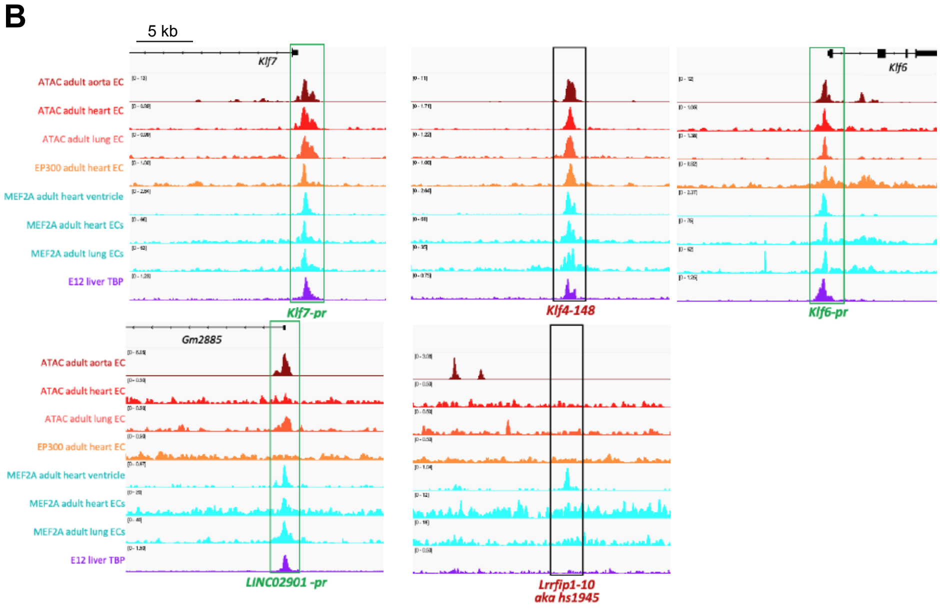


**C** LINC02901 MEF2-TBP motif

```
human     GCGCGACCAAAATAGCGCCATATAAAAGC---TGG
mouse     GCGCGACTAAAATAGCGCAGTATAAAAGC--TGGC
cow       GCAAGACCAAAATAGTGCCGTATAAATGC--GTGC
opossum   ACGCGATTAAAATAGCGCTGTATAAATACCCGGAC
           *   **  ******* **  ******   *
```

**D** LRRFIP1 MEF2-TBP motif

```
human     TTACCATATTTTTGGCAGTGGCTTTTAAAAGCACTC
mouse     TTCTCATATTTTTGGCAGTGACTTTTAAAGCACGTC
pig       TTCCCATATTTTTAGCAGGGGCTTTTAAGAGTGCTC
tenrec    GGAGGTTTTCTTTGGCTGTGGCTTTTTGAAAACAGC
               * * *** ** * * *****         *
```

**E**

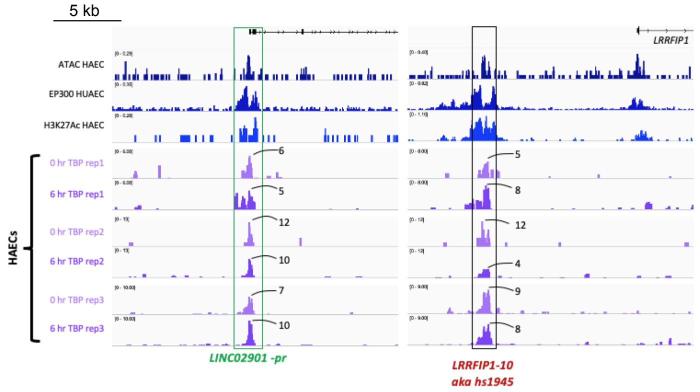


**F** hs1945 enhancer from Vista Enhancer Browser

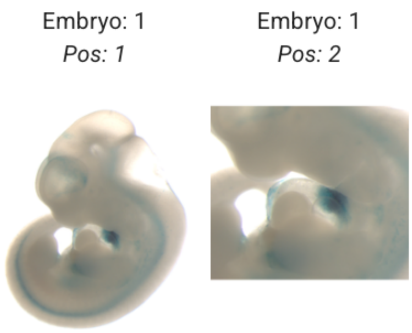

**Supplemental Figure S9**

**The MEF2-TBP double motif is found within regulatory elements for other *KLF* genes, which are bound by TBP in a FSS-sensitive manner**
**A** Sequence conservation around the MEF2-TBP motifs within the *KLF6* promoter, the *KLF7* promoter and the *KLF4* upstream element. In each case, the MEF2-TBP double motif (blue-purple) shows much deeper conservation (bold text and *) than the surrounding sequences (ETS motifs in green are also conserved).
**B** The *KLF6*, *KLF7* and *KLF4* elements containing MEF2-TBP motifs were also associated with enhancer/promoter marks in mouse (ATAC-seq (ATAC) adult aorta EC from [75], adult heart and lung EC ATAC-seq and EP300 binding from[76]) and bound by MEF2A and MEF2C in adult heart and lung ECs. Of the two other elements containing non-consensus MEF2-TBP motifs, *LINC02901* promoter showed similar patterns but *LRRFIP1-10* did not.
**C-D** Two other elements containing non-consensus MEF2-TBP motifs did not show similar levels of deep conservation of the MEF2-TBP double motif (blue-mauve): the LINC02901 MEF2-TBP motif was less conserved than surrounding sequence (**C**, image from UCSD browser), and the LRRFIP1 MEF2-TBP motif showing little conservation beyond human-mouse (**D**).
**E** TBP binding at the two non-consensus MEF2-TBP motifs (*LINC02901* promoter and *LRRFIP1-10*) was low and did not consistently increase after FSS.
**F** Data from the Vista Enhancer Browser indicates that *LRRFIP1-10* lies within an enhancer active in the outflow tract.

#### Figure S10

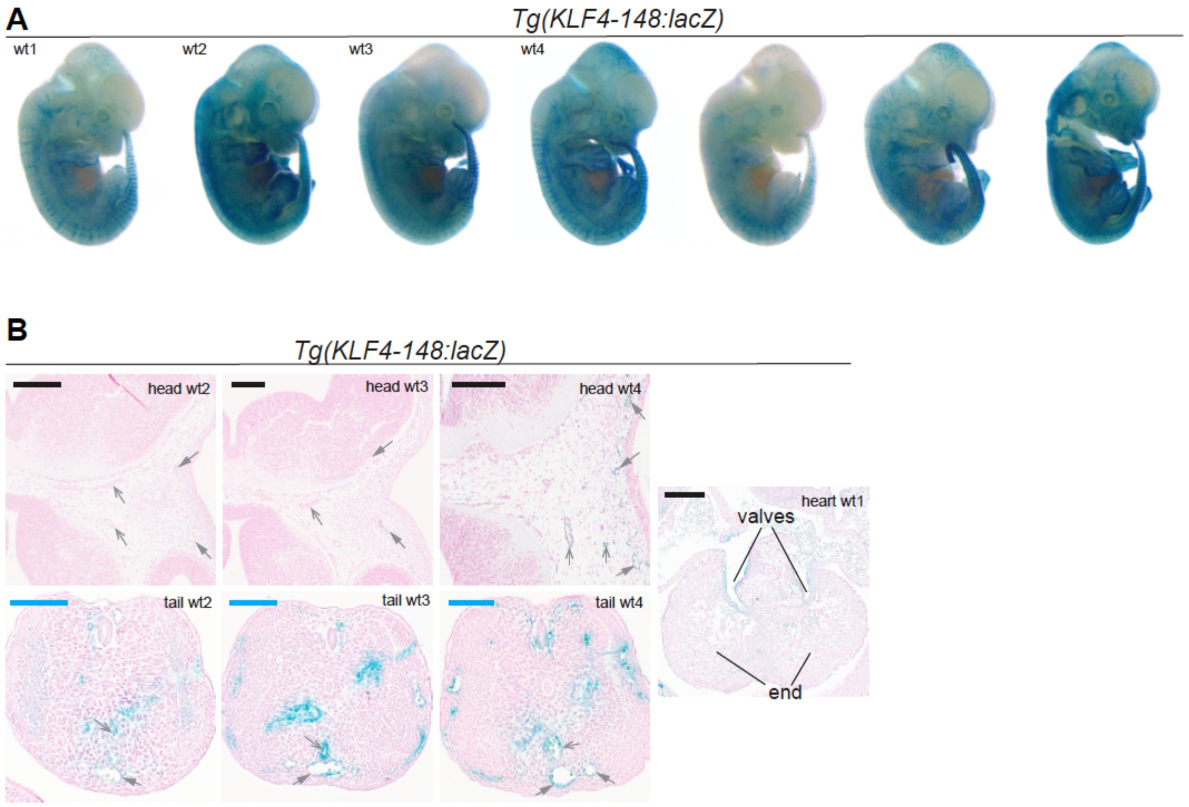


**C**

| | *KLF4-148* |
|---|---|
| *Tg mice with EC expression* | 7 |
| *Tg mice with non-EC expression* | 0 |
| *Tg mice with no expression* | 0 |
| ***Total number of tg mice*** | **7** |

**Supplemental Figure S10**

**F0 transgenic embryos expressing the *KLF4-148:lacZ* transgene**
Wholemount images of all X-gal positive E12.5 F0 embryos transgenic for the *KLF4-148:lacZ* transgene (**A**) and transverse sections through selected embryos (**B**). **C** summarizes numbers. Black scale bar is 200µm, blue scale bar is 100µm, grey open arrow is artery, grey closed arrow is vein.

#### Supplemental Table 1

| Nearest Gene Name | location of TBP peak hg38 | TBP motif start | MEF2 motif start | Distance between motifs | Feature type |
|---|---|---|---|---|---|
| LINC01767 | chr1 56456650 56458250 | 56458102 | 56458093 | 0 | Enhancer |
| S100A10 | chr1 151988545 151989800 | 151989098 | 151989109 | 0 | Promoter/Enhancer |
| FLJ13224 | chr12 28313660 28313861 | 28313809 | 28313798 | 0 | Enhancer |
| CNOT3 | chr19 49641589 49642162 | 49641626 | 49641615 | 0 | Promoter/Enhancer |
| LINC01817 | chr2 135530749 135531426 | 135531073 | 135531084 | 0 | Promoter/Enhancer |
| CDCP1 | chr3 42025726 42025953 | 42025811 | 42025800 | 0 | Enhancer |
| GBE1 | chr3 71055759 71056950 | 71056840 | 71056851 | 0 | Promoter/Enhancer |
| ZBTB20 | chr3 114558388 114558567 | 114558455 | 114558444 | 0 | Enhancer |
| LINC02266 | chr4 138239500 138243150 | 138242833 | 138242822 | 0 | Promoter/Enhancer |
| CLCN3 | chr4 169225400 169226150 | 169225773 | 169225784 | 0 | Enhancer |
| H3C4 | chr6 26155900 26158711 | 26157901 | 26157912 | 0 | Promoter/Enhancer |
| CCN2 | chr6 121482069 121482270 | 121482071 | 121482082 | 0 | Enhancer |
| ICA1 | chr6 158869700 158870000 | 158869779 | 158869768 | 0 | Promoter/Enhancer |
| DIPK2B | chr9 124658250 124659050 | 124659025 | 124659016 | 0 | Promoter/Enhancer |
| LINC01714 | chr1 8194600 8194900 | 8194769 | 8194781 | 1 | Enhancer |
| RNU5D-1 | chr1 44730850 44731450 | 44731421 | 44731411 | 1 | Promoter/Enhancer |
| MIR5191 | chr1 198935643 198936062 | 198935980 | 198935990 | 1 | Promoter/Enhancer |
| PROSER2-AS1 | chr10 10796608 10796764 | 10796738 | 10796748 | 1 | Promoter/Enhancer |
| ME3 | chr11 85947875 85948238 | 85947909 | 85947921 | 1 | Enhancer |
| DPH6-DT | chr15 34334111 34335689 | 34335477 | 34335467 | 1 | Promoter/Enhancer |
| SUMO1P1 | chr20 51661000 51661883 | 51661678 | 51661668 | 1 | Enhancer |
| HES1 | chr3 187265600 187265850 | 187265620 | 187265630 | 1 | Enhancer |
| BTN3A2 | chr6 26327228 26328951 | 26328646 | 26328634 | 1 | Promoter/Enhancer |
| ARHGAP18 | chr6 118708900 118709440 | 118708913 | 118708923 | 1 | Promoter/Enhancer |
| SNORD151 | chr7 39592626 39592850 | 39592767 | 39592755 | 1 | Enhancer |
| MIR1208 | chr8 125219250 125219537 | 125219411 | 125219421 | 1 | Enhancer |
| MACO1 | chr1 25430323 25430850 | 25430652 | 25430665 | 2 | Promoter/Enhancer |
| RNF11 | chr1 51237001 51237978 | 51237698 | 51237707 | 2 | Promoter/Enhancer |
| KLF6 | chr10 3784933 3786509 | 3785521 | 3785508 | 2 | Promoter/Enhancer |
| KLF6 | chr10 3804500 3807040 | 3804970 | 3804983 | 2 | Promoter/Enhancer |
| SAMD4A | chr14 54650200 54650780 | 54650300 | 54650309 | 2 | Promoter/Enhancer |
| MIR1302-4 | chr2 207165507 207166950 | 207166350 | 207166337 | 2 | Promoter/Enhancer |
| LINC00316 | chr21 38486493 38486684 | 38486545 | 38486532 | 2 | Enhancer |
| H3C6 | chr6 26216137 26217900 | 26217607 | 26217620 | 2 | Promoter/Enhancer |
| PLS3 | chrX 45824495 45824800 | 45824514 | 45824523 | 2 | Enhancer |
| LOC100505502 | chr10 31320983 31321067 | 31321005 | 31320991 | 3 | Promoter/Enhancer |
| DUSP6 | chr12 89346416 89347183 | 89347028 | 89347014 | 3 | Promoter/Enhancer |
| MIR5006 | chr13 41050165 41050343 | 41050238 | 41050226 | 3 | Enhancer |
| UQCRC2 | chr16 12801660 12801871 | 12801762 | 12801748 | 3 | Promoter/Enhancer |
| KLF2 | chr19 16258400 16258975 | 16258700 | 16258686 | 3 | Enhancer |
| KLF2 | chr19 16283150 16284100 | 16283782 | 16283768 | 3 | Promoter/Enhancer |
| SNORA118 | chr19 16324450 16325050 | 16324691 | 16324705 | 3 | Promoter/Enhancer |
| H1-1 | chr6 25006720 25006946 | 25006792 | 25006778 | 3 | Enhancer |
| AFDN | chr6 152696436 152698950 | 152697863 | 152697855 | 3 | Promoter/Enhancer |
| TCIM | chr8 26608160 26609883 | 26609150 | 26609164 | 3 | Enhancer |
| PALM2AKAP2 | chr9 107637350 107639043 | 107637748 | 107637762 | 3 | Promoter/Enhancer |
| IL13RA2 | chrX 45802134 45803950 | 45803643 | 45803657 | 3 | Enhancer |
| PPP1R15B | chr1 201713577 201713984 | 201713963 | 201713948 | 4 | Enhancer |
| MIR5188 | chr12 124913750 124918150 | 124916002 | 124915987 | 4 | Promoter/Enhancer |
| LRRFIP1 | chr2 237616850 237617250 | 237617081 | 237617096 | 4 | Enhancer |
| FEM1B | chr15 65295800 65296476 | 65296163 | 65296179 | 5 | Promoter/Enhancer |
| CORO6 | chr17 17446050 17446259 | 17446224 | 17446208 | 5 | Promoter/Enhancer |
| SERPINB2 | chr18 62281751 62281922 | 62281885 | 62281869 | 5 | Enhancer |
| GPD2 | chr2 150487318 150487939 | 150487501 | 150487517 | 5 | Promoter/Enhancer |
| ZSWIM2 | chr2 186590774 186591300 | 186591154 | 186591170 | 5 | Promoter/Enhancer |
| HMGB2 | chr4 173057700 173058200 | 173057874 | 173057890 | 5 | Enhancer |
| LINC02509 | chr4 173318703 173318900 | 173318792 | 173318808 | 5 | Enhancer |
| MIR4638 | chr5 172872985 172873302 | 172873100 | 172873116 | 5 | Enhancer |
| LINC01556 | chr6 28880850 28881661 | 28881605 | 28881621 | 5 | Promoter/Enhancer |
| PNRC1 | chr6 77456618 77457000 | 77456797 | 77456813 | 5 | Promoter/Enhancer |
| GNA12 | chr6 154432797 154433107 | 154432928 | 154432944 | 5 | Enhancer |
| UTP11 | chr1 38012500 38013000 | 38012564 | 38012581 | 6 | Promoter/Enhancer |
| PRSS23 | chr11 86597296 86597497 | 86597427 | 86597410 | 6 | Enhancer |
| PRICKLE1 | chr12 31731650 31732443 | 31732086 | 31732103 | 6 | Enhancer |
| LINC00324 | chr17 8118450 8121800 | 8119287 | 8119270 | 6 | Promoter/Enhancer |
| ANXA5 | chr4 98932569 98932909 | 98932766 | 98932749 | 6 | Enhancer |
| H2BC10 | chr6 26271250 26272181 | 26271935 | 26271952 | 6 | Promoter/Enhancer |
| HSPG2 | chr1 21910062 21910246 | 21910207 | 21910189 | 7 | Promoter/Enhancer |
| FNDC3A | chr13 45464695 45464896 | 45464843 | 45464827 | 7 | Promoter/Enhancer |

| Nearest Gene Name | location of TBP peak hg38 | TBP motif start | MEF2 motif start | Distance between motifs | Feature type |
|---|---|---|---|---|---|
| FOSB | chr19 45428800 45429001 | 45428889 | 45428907 | 7 | Promoter/Enhancer |
| GBE1 | chr3 81425540 81425741 | 81425581 | 81425599 | 7 | Enhancer |
| LRRC17 | chr7 55065148 55065531 | 55065461 | 55065479 | 7 | Enhancer |
| APBA1 | chr9 43140135 43140750 | 43140495 | 43140513 | 7 | Enhancer |
| ZFAND5 | chr9 63347461 63348000 | 63347675 | 63347693 | 7 | Enhancer |
| PLS3 | chrX 115594120 115595356 | 115594891 | 115594909 | 7 | Promoter/Enhancer |
| BCAR3 | chr1 93555060 93555230 | 93555169 | 93555150 | 8 | Enhancer |
| SDE2 | chr1 223831795 223832200 | 223831932 | 223831913 | 8 | Enhancer |
| DPH6-DT | chr15 35692362 35692550 | 35692503 | 35692484 | 8 | Enhancer |
| C2CD4A | chr15 58520828 58521029 | 58520915 | 58520896 | 8 | Promoter/Enhancer |
| TTC19 | chr17 8226027 8227574 | 8227109 | 8227090 | 8 | Promoter/Enhancer |
| LINC02595 | chrX 45479450 45479591 | 45479467 | 45479486 | 8 | Enhancer |
| GEMIN8P4 | chr1 89908872 89909200 | 89908900 | 89908920 | 9 | Enhancer |
| GTF2A1 | chr14 81218850 81220038 | 81219384 | 81219364 | 9 | Promoter/Enhancer |
| CYB561 | chr17 62572754 62573318 | 62573007 | 62572987 | 9 | Enhancer |
| SLC6A6 | chr3 12841587 12841795 | 12841684 | 12841704 | 9 | Promoter/Enhancer |
| RACK1 | chr5 181221808 181223750 | 181222936 | 181222916 | 9 | Promoter/Enhancer |
| STEAP1B | chr7 2757994 2758370 | 2758101 | 2758121 | 9 | Promoter/Enhancer |
| CAV2 | chr7 108416162 108416600 | 108416277 | 108416297 | 9 | Enhancer |
| GADD45A | chr1 67726054 67726600 | 67726076 | 67726097 | 10 | Enhancer |
| PTPRE | chr10 122112050 122112600 | 122112181 | 122112160 | 10 | Promoter/Enhancer |
| DUSP16 | chr12 10723266 10723505 | 10723347 | 10723326 | 10 | Promoter/Enhancer |
| LINC00924 | chr15 92900001 92900700 | 92900508 | 92900487 | 10 | Promoter/Enhancer |
| LINC00324 | chr17 8220485 8223450 | 8220667 | 8220646 | 10 | Promoter/Enhancer |
| DDX5 | chr17 64497223 64497424 | 64497412 | 64497391 | 10 | Promoter/Enhancer |
| LOC105374988 | chr6 26330191 26331850 | 26331232 | 26331253 | 10 | Promoter/Enhancer |
| LOC105374988 | chr6 26330191 26331850 | 26331232 | 26331253 | 10 | Enhancer |
| ABT1 | chr6 26552781 26556955 | 26553871 | 26553850 | 10 | Promoter/Enhancer |
| H2BC11 | chr6 26552781 26556955 | 26553871 | 26553850 | 10 | Enhancer |
| PRKAR2B | chr7 76302371 76302750 | 76302641 | 76302620 | 10 | Promoter/Enhancer |
| RPS20 | chr8 40098173 40098365 | 40098212 | 40098191 | 10 | Enhancer |
| TEK | chr9 18393306 18393783 | 18393724 | 18393703 | 10 | Enhancer |

**Supplemental Table S1**
List of genomic locations bound by TBP in human cells and containing called motifs for both MEF2A and TBP separated by less than 100 bp.

#### Supplemental Table S2

| Nearest Gene Name | location of MEF2A peak mm10 | TBP motif start | MEF2 motif start | Distance between motifs | Feature type |
|---|---|---|---|---|---|
| Sec16b | chr1 157494978 157495143 | 157495025 | 157495036 | 0 | Distal Intergenic |
| Cd34 | chr1 194925860 194925988 | 194925928 | 194925939 | 0 | Distal Intergenic |
| Srgap1 | chr10 121836790 121836959 | 121836863 | 121836852 | 0 | Intron (ENSMUST00000020322.11/117600, intron 10 of 21) |
| 2410006H16Rik | chr11 62602724 62602868 | 62602843 | 62602834 | 0 | Promoter (<=1kb) |
| Serpinb1c | chr13 32922669 32922875 | 32922822 | 32922833 | 0 | Distal Intergenic |
| Dleu2 | chr14 61680790 61681059 | 61680943 | 61680934 | 0 | Promoter (1-2kb) |
| Dach1 | chr14 97588077 97588363 | 97588193 | 97588204 | 0 | Distal Intergenic |
| Dach1 | chr14 97970738 97970968 | 97970776 | 97970787 | 0 | Intron (ENSMUST00000069334.7/13134, intron 2 of 10) |
| Pick1 | chr15 79243455 79243656 | 79243570 | 79243559 | 0 | Promoter (<=1kb) |
| Slc38a2 | chr15 96716338 96716576 | 96716511 | 96716500 | 0 | Distal Intergenic |
| Hmgn1 | chr16 96128089 96128290 | 96128213 | 96128224 | 0 | Promoter (<=1kb) |
| Arhgap12 | chr18 5945389 5945590 | 5945436 | 5945447 | 0 | Intron (ENSMUST00000234953.1/ENSMUST00000234953.1, intron 1 of 4) |
| Dnajc18 | chr18 35677530 35677712 | 35677672 | 35677683 | 0 | Intron (ENSMUST00000025208.6/76594, intron 7 of 7) |
| Plce1 | chr19 38498265 38498472 | 38498333 | 38498324 | 0 | Intron (ENSMUST00000181994.1/ENSMUST00000181994.1, intron 1 of 2) |
| Mov10 | chr3 104819644 104819847 | 104819662 | 104819673 | 0 | Promoter (1-2kb) |
| A930031H19Rik | chr4 131244089 131244300 | 131244217 | 131244206 | 0 | Distal Intergenic |
| Kazn | chr4 142166355 142166592 | 142166543 | 142166554 | 0 | Intron (ENSMUST00000155023.7/71529, intron 1 of 13) |
| Evi5 | chr5 107873816 107874017 | 107873907 | 107873918 | 0 | Promoter (1-2kb) |
| Stard13 | chr5 151047157 151047333 | 151047288 | 151047277 | 0 | Intron (ENSMUST00000110483.8/243362, intron 9 of 13) |
| Kbtbd12 | chr6 88622129 88622322 | 88622197 | 88622208 | 0 | Intron (ENSMUST00000120933.4/74589, intron 1 of 5) |
| Inpp5a | chr7 139434173 139434383 | 139434265 | 139434276 | 0 | Intron (ENSMUST00000106098.7/212111, intron 1 of 15) |
| Col4a1 | chr8 11183698 11183907 | 11183726 | 11183737 | 0 | Distal Intergenic |
| Pth1r | chr9 110750946 110751142 | 110751052 | 110751041 | 0 | Intron (ENSMUST00000200011.4/17897, intron 1 of 5) |
| 1700020M21Rik | chr9 120852542 120852742 | 120852629 | 120852640 | 0 | Exon (ENSMUST00000215206.1/ENSMUST00000215206.1, exon 3 of 3) |
| C730036E19Rik | chr1 151144839 151145070 | 151144935 | 151144945 | 1 | Distal Intergenic |
| Gfod1 | chr13 43317592 43317766 | 43317735 | 43317723 | 1 | Distal Intergenic |
| Cdh18 | chr15 22331482 22331610 | 22331483 | 22331495 | 1 | Distal Intergenic |
| Nsmce2 | chr15 59480658 59480810 | 59480731 | 59480741 | 1 | Intron (ENSMUST00000168722.2/68501, intron 4 of 4) |
| Klhl24 | chr16 20091231 20091449 | 20091282 | 20091294 | 1 | Distal Intergenic |
| 4833419F23Rik | chr18 4354632 4354836 | 4354687 | 4354699 | 1 | Promoter (<=1kb) |
| Hectd2os | chr19 36662865 36663079 | 36662908 | 36662900 | 1 | Intron (ENSMUST00000149601.7/668215, intron 1 of 4) |
| Plcb1 | chr2 134881963 134882164 | 134882062 | 134882050 | 1 | Intron (ENSMUST00000131552.4/18795, intron 2 of 32) |
| Pakap | chr4 57755492 57755627 | 57755574 | 57755586 | 1 | Intron (ENSMUST00000098066.8/677884, intron 6 of 10) |
| Auts2 | chr5 131517750 131517977 | 131517936 | 131517948 | 1 | Intron (ENSMUST00000161226.8/319974, intron 6 of 18) |
| E230013L22Rik | chr8 11474199 11474394 | 11474313 | 11474323 | 1 | Intron (ENSMUST00000033900.6/19332, intron 1 of 1) |
| BC048562 | chr9 108441370 108441611 | 108441587 | 108441575 | 1 | Intron (ENSMUST00000057265.7/434439, intron 2 of 3) |
| Otc | chrX 10281483 10281692 | 10281570 | 10281582 | 1 | Intron (ENSMUST00000049910.12/18416, intron 4 of 9) |
| Klf7 | chr1 64121704 64121931 | 64121824 | 64121811 | 2 | Promoter (<=1kb) |
| Mapkapk2 | chr1 131116518 131116687 | 131116558 | 131116571 | 2 | Distal Intergenic |
| Srgap2 | chr1 131428181 131428341 | 131428264 | 131428277 | 2 | Intron (ENSMUST00000097588.8/14270, intron 3 of 22) |
| Rgs5 | chr1 169685241 169685451 | 169685319 | 169685328 | 2 | Intron (ENSMUST00000027997.8/19737, intron 3 of 4) |
| Hdac9 | chr12 34452657 34452858 | 34452741 | 34452754 | 2 | Intron (ENSMUST00000209902.1/79221, intron 1 of 24) |
| Klf6 | chr13 5861130 5861331 | 5861219 | 5861232 | 2 | Promoter (<=1kb) |
| A330076C08Rik | chr13 44248567 44248804 | 44248770 | 44248757 | 2 | Distal Intergenic |
| Drosha | chr15 12934297 12934498 | 12934437 | 12934450 | 2 | Intron (ENSMUST00000169061.7/14000, intron 34 of 34) |
| Gm5468 | chr15 25459995 25460195 | 25460164 | 25460173 | 2 | Exon (ENSMUST00000228099.1/ENSMUST00000228099.1, exon 1 of 1) |
| Ezr | chr17 6827841 6828061 | 6827948 | 6827935 | 2 | Distal Intergenic |
| Adra1d | chr2 131683835 131684032 | 131683974 | 131683987 | 2 | Intron (ENSMUST00000138080.1/ENSMUST00000138080.1, intron 2 of 2) |
| Nfkb1 | chr3 135603029 135603245 | 135603174 | 135603161 | 2 | Promoter (<=1kb) |
| Camta1 | chr4 151862484 151862685 | 151862583 | 151862596 | 2 | Promoter (<=1kb) |
| Pole4 | chr6 82529077 82529217 | 82529141 | 82529154 | 2 | Intron (ENSMUST00000032122.10/21336, intron 3 of 5) |
| Col4a2 | chr8 11374053 11374277 | 11374136 | 11374127 | 2 | Intron (ENSMUST00000033899.13/12827, intron 4 of 47) |
| Agap1 | chr1 89613558 89613710 | 89613623 | 89613637 | 3 | Intron (ENSMUST00000027521.14/347722, intron 3 of 17) |
| Itga7 | chr10 128931395 128931580 | 128931466 | 128931480 | 3 | Promoter (2-3kb) |
| Pitpnm3 | chr11 72086810 72087019 | 72086887 | 72086901 | 3 | Intron (ENSMUST00000108508.2/327958, intron 3 of 18) |
| Pecam1 | chr11 106730796 106730992 | 106730907 | 106730921 | 3 | Intron (ENSMUST00000068021.8/18613, intron 1 of 15) |
| Prss16 | chr13 22015803 22015943 | 22015882 | 22015896 | 3 | Intron (ENSMUST00000147811.2/ENSMUST00000147811.2, intron 1 of 2) |
| Hrh2 | chr13 54177458 54177660 | 54177500 | 54177514 | 3 | Distal Intergenic |
| 9230112D13Rik | chr14 34544646 34544862 | 34544777 | 34544763 | 3 | Intron (ENSMUST00000022327.12/24131, intron 9 of 13) |
| Epb41l4a | chr18 33876438 33876639 | 33876541 | 33876549 | 3 | Intron (ENSMUST00000025234.6/13824, intron 8 of 23) |
| Jag1 | chr2 137012042 137012243 | 137012130 | 137012144 | 3 | Intron (ENSMUST00000028737.12/74243, intron 3 of 5) |
| Jph2 | chr2 163392510 163392716 | 163392619 | 163392605 | 3 | Intron (ENSMUST00000017961.10/59091, intron 1 of 5) |
| Klf4 | chr4 55651158 55651370 | 55651235 | 55651249 | 3 | Distal Intergenic |
| Igbp1b | chr6 138803858 138804049 | 138803945 | 138803959 | 3 | Distal Intergenic |
| Ap1m1 | chr8 72268461 72268713 | 72268578 | 72268564 | 3 | Distal Intergenic |
| Klf2 | chr8 72295792 72295994 | 72295879 | 72295865 | 3 | Distal Intergenic |
| Klf2 | chr8 72318832 72319054 | 72318929 | 72318943 | 3 | Promoter (<=1kb) |
| Sgpp2 | chr1 78371659 78371858 | 78371771 | 78371786 | 4 | Intron (ENSMUST00000036172.9/433323, intron 2 of 4) |
| Rab17 | chr1 90980737 90980960 | 90980891 | 90980906 | 4 | Distal Intergenic |

| Nearest Gene Name | location of MEF2A peak mm10 | TBP motif start | MEF2 motif start | Distance between motifs | Feature type |
|---|---|---|---|---|---|
| Slit3 | chr11 35231930 35232046 | 35231955 | 35231940 | 4 | Intron (ENSMUST00000069837.3/20564, intron 3 of 35) |
| Gm12159 | chr11 45244717 45244883 | 45244822 | 45244807 | 4 | Distal Intergenic |
| 4930526H09Rik | chr13 110941651 110941888 | 110941718 | 110941733 | 4 | Distal Intergenic |
| Trpm3 | chr19 22905327 22905533 | 22905432 | 22905417 | 4 | Intron (ENSMUST00000237357.1/226025, intron 17 of 24) |
| Hspa12a | chr19 58819052 58819260 | 58819131 | 58819146 | 4 | Promoter (2-3kb) |
| 9030204H09Rik | chr2 35788577 35788705 | 35788665 | 35788650 | 4 | Intron (ENSMUST00000028248.10/74410, intron 8 of 9) |
| Adamtsl4 | chr3 95695328 95695560 | 95695475 | 95695490 | 4 | Distal Intergenic |
| Rps4l | chr6 148360475 148360662 | 148360562 | 148360577 | 4 | Intron (ENSMUST00000204618.1/ENSMUST00000204618.1, intron 1 of 1) |
| Ercc1 | chr7 19332873 19333083 | 19332941 | 19332956 | 4 | Intron (ENSMUST00000207444.1/ENSMUST00000207444.1, intron 1 of 1) |
| Bag3 | chr7 128538297 128538498 | 128538435 | 128538420 | 4 | Intron (ENSMUST00000033136.8/29810, intron 1 of 3) |
| Hand2os1 | chr8 57205129 57205258 | 57205186 | 57205171 | 4 | Distal Intergenic |
| Plcg2 | chr8 117543888 117544108 | 117543999 | 117543984 | 4 | Intron (ENSMUST00000081232.8/234779, intron 2 of 32) |
| Ppp2r3d | chr9 124478218 124478513 | 124478445 | 124478460 | 4 | Promoter (1-2kb) |
| Sox17 | chr1 4617809 4618053 | 4617972 | 4617956 | 5 | Distal Intergenic |
| Pou2f1 | chr1 165993401 165993498 | 165993433 | 165993449 | 5 | Intron (ENSMUST00000160260.8/18986, intron 1 of 16) |
| Kif28 | chr1 179733503 179733704 | 179733539 | 179733555 | 5 | Intron (ENSMUST00000131716.3/383592, intron 4 of 22) |
| 4933440J02Rik | chr10 111583168 111583384 | 111583309 | 111583325 | 5 | Distal Intergenic |
| 5430427M07Rik | chr12 91040481 91040699 | 91040620 | 91040636 | 5 | Intron (ENSMUST00000143415.7/75216, intron 8 of 12) |
| Adtrp | chr13 41729741 41729980 | 41729910 | 41729894 | 5 | Distal Intergenic |
| Cap2 | chr13 46572109 46572319 | 46572204 | 46572220 | 5 | Intron (ENSMUST00000021802.15/67252, intron 4 of 12) |
| St3gal1 | chr15 67147406 67147607 | 67147501 | 67147485 | 5 | Intron (ENSMUST00000229028.1/20442, intron 2 of 8) |
| Crim1 | chr17 77033587 77033833 | 77033722 | 77033738 | 5 | Distal Intergenic |
| Myoz3 | chr18 60595266 60595421 | 60595299 | 60595315 | 5 | 3' UTR |
| Mn1 | chr5 111473569 111473789 | 111473706 | 111473690 | 5 | Distal Intergenic |
| Kcna6 | chr6 126699827 126699947 | 126699881 | 126699897 | 5 | Distal Intergenic |
| Gas2 | chr7 51913818 51914021 | 51913991 | 51913975 | 5 | Intron (ENSMUST00000129604.7/14453, intron 5 of 9) |
| E230029C05Rik | chr7 90023308 90023508 | 90023448 | 90023464 | 5 | Intron (ENSMUST00000207458.1/319711, intron 7 of 8) |
| Hlx | chr1 184748803 184749004 | 184748955 | 184748972 | 6 | Distal Intergenic |
| 1700016G22Rik | chr13 5604299 5604453 | 5604352 | 5604335 | 6 | Intron (ENSMUST00000222871.1/ENSMUST00000222871.1, intron 2 of 2) |
| Xrcc4 | chr13 90012305 90012533 | 90012424 | 90012407 | 6 | Intron (ENSMUST00000159199.7/108138, intron 3 of 8) |
| Nox3 | chr17 3632533 3632715 | 3632619 | 3632602 | 6 | Distal Intergenic |
| Sgms1 | chr19 32169862 32170114 | 32169901 | 32169884 | 6 | Intron (ENSMUST00000099514.9/208449, intron 5 of 9) |
| Camk1d | chr2 5572690 5572861 | 5572809 | 5572826 | 6 | Intron (ENSMUST00000044009.13/227541, intron 1 of 10) |
| Usp13 | chr3 32818681 32818864 | 32818767 | 32818784 | 6 | Promoter (<=1kb) |
| Gm20750 | chr3 52899742 52899905 | 52899858 | 52899841 | 6 | Distal Intergenic |
| Mttp | chr3 138142835 138143036 | 138142959 | 138142942 | 6 | Promoter (<=1kb) |
| Ptafr | chr4 132573793 132573994 | 132573894 | 132573911 | 6 | Intron (ENSMUST00000070690.7/19204, intron 1 of 1) |
| Eif4g2 | chr7 111068251 111068447 | 111068330 | 111068313 | 6 | 3' UTR |
| Otoa | chr7 121111333 121111560 | 121111466 | 121111483 | 6 | Intron (ENSMUST00000047025.14/246190, intron 10 of 28) |
| Ing2 | chr8 47604547 47604748 | 47604652 | 47604669 | 6 | Distal Intergenic |
| Gk | chrX 85731300 85731501 | 85731367 | 85731384 | 6 | Intron (ENSMUST00000142152.1/14933, intron 11 of 19) |
| Sipa1l1 | chr12 82374201 82374346 | 82374289 | 82374271 | 7 | Intron (ENSMUST00000166429.8/217692, intron 7 of 22) |
| Fdft1 | chr14 63150165 63150369 | 63150201 | 63150219 | 7 | Intron (ENSMUST00000238526.1/14137, intron 6 of 6) |
| Gm19510 | chr15 59775929 59776147 | 59776076 | 59776058 | 7 | Distal Intergenic |
| Mrtfb | chr16 13358111 13358329 | 13358237 | 13358255 | 7 | Promoter (<=1kb) |
| Smco1 | chr16 32271340 32271537 | 32271445 | 32271427 | 7 | Promoter (<=1kb) |
| Jam2 | chr16 84773411 84773619 | 84773514 | 84773532 | 7 | Promoter (<=1kb) |
| Bambi | chr18 3665773 3665982 | 3665795 | 3665813 | 7 | Distal Intergenic |
| Orc4 | chr2 49023261 49023472 | 49023395 | 49023377 | 7 | Intron (ENSMUST00000112745.7/109241, intron 2 of 10) |
| Polr1e | chr4 45038194 45038426 | 45038369 | 45038351 | 7 | Intron (ENSMUST00000052236.12/269529, intron 10 of 10) |
| Nfib | chr4 82696162 82696367 | 82696269 | 82696251 | 7 | Intron (ENSMUST00000155821.1/18028, intron 1 of 1) |
| Hspg2 | chr4 137488809 137489024 | 137488900 | 137488918 | 7 | Intron (ENSMUST00000030547.14/15530, intron 1 of 96) |
| Mir1960 | chr5 30172128 30172294 | 30172165 | 30172183 | 7 | Promoter (1-2kb) |
| Tbc1d19 | chr5 53929398 53929606 | 53929497 | 53929479 | 7 | Distal Intergenic |
| Tec | chr5 72802941 72803142 | 72803064 | 72803046 | 7 | Intron (ENSMUST00000071944.12/21682, intron 3 of 18) |
| Ercc1 | chr7 19342130 19342397 | 19342229 | 19342211 | 7 | Promoter (2-3kb) |
| 1700109G14Rik | chr14 61330974 61331228 | 61331072 | 61331091 | 8 | Distal Intergenic |
| Vwa8 | chr14 78990478 78990654 | 78990604 | 78990585 | 8 | Intron (ENSMUST00000040990.6/219189, intron 14 of 44) |
| Casp7 | chr19 56422596 56422799 | 56422735 | 56422716 | 8 | Promoter (2-3kb) |
| 4930547E08Rik | chr2 103881103 103881228 | 103881148 | 103881129 | 8 | Distal Intergenic |
| Fndc3b | chr3 27656890 27657142 | 27657035 | 27657016 | 8 | Intron (ENSMUST00000195008.5/72007, intron 2 of 26) |
| Apbb2 | chr5 66323678 66323882 | 66323728 | 66323747 | 8 | Intron (ENSMUST00000162366.7/11787, intron 10 of 15) |
| Serpine1 | chr5 137072212 137072419 | 137072292 | 137072311 | 8 | Promoter (<=1kb) |
| Oxtr | chr6 112577386 112577584 | 112577449 | 112577468 | 8 | Distal Intergenic |
| Mob2 | chr7 142074150 142074367 | 142074271 | 142074252 | 8 | Distal Intergenic |
| Mir5104 | chr10 7841836 7842071 | 7842001 | 7841981 | 9 | Intron (ENSMUST00000039484.5/237256, intron 2 of 5) |
| Timp3 | chr10 86313660 86313855 | 86313705 | 86313725 | 9 | Intron (ENSMUST00000020234.13/21859, intron 1 of 4) |
| Abca13 | chr11 10002239 10002439 | 10002336 | 10002316 | 9 | Distal Intergenic |
| Bod1 | chr11 31663437 31663642 | 31663527 | 31663547 | 9 | Distal Intergenic |
| Alox12 | chr11 70266404 70266659 | 70266565 | 70266545 | 9 | Distal Intergenic |

| Nearest Gene Name | location of MEF2A peak mm10 | TBP motif start | MEF2 motif start | Distance between motifs | Feature type |
|---|---|---|---|---|---|
| Dip2c | chr13 9516314 9516490 | 9516370 | 9516350 | 9 | Intron (ENSMUST00000174552.7/208440, intron 3 of 36) |
| Mir5624 | chr13 93887144 93887343 | 93887200 | 93887220 | 9 | Intron (ENSMUST00000091403.5/11881, intron 6 of 7) |
| Slc38a9 | chr13 112697956 112698142 | 112698121 | 112698101 | 9 | Promoter (2-3kb) |
| Plpp1 | chr13 112764591 112764758 | 112764655 | 112764675 | 9 | Distal Intergenic |
| Vwa8 | chr14 79076990 79077172 | 79077080 | 79077060 | 9 | Intron (ENSMUST00000040990.6/219189, intron 26 of 44) |
| Arhgap26 | chr18 39215222 39215433 | 39215332 | 39215312 | 9 | Intron (ENSMUST00000097593.8/71302, intron 12 of 22) |
| Bcas1 | chr2 170278103 170278293 | 170278183 | 170278163 | 9 | Distal Intergenic |
| Tcea3 | chr4 136261885 136262086 | 136262010 | 136262030 | 9 | Intron (ENSMUST00000136812.1/21401, intron 5 of 8) |
| Limch1 | chr5 66857201 66857360 | 66857306 | 66857286 | 9 | Intron (ENSMUST00000201852.3/77569, intron 1 of 7) |
| Pitpnm2 | chr5 124171725 124171950 | 124171828 | 124171808 | 9 | Intron (ENSMUST00000086123.10/19679, intron 1 of 23) |
| Smurf1 | chr5 144929010 144929245 | 144929120 | 144929140 | 9 | Intron (ENSMUST00000110677.7/75788, intron 1 of 18) |
| Nup210 | chr6 91076336 91076568 | 91076490 | 91076470 | 9 | Exon (ENSMUST00000032179.13/54563, exon 7 of 40) |
| Asb5 | chr8 54393599 54393807 | 54393698 | 54393678 | 9 | Distal Intergenic |
| Ddx18 | chr1 121631541 121631742 | 121631596 | 121631575 | 10 | Distal Intergenic |
| Tmem163 | chr1 127596242 127596415 | 127596285 | 127596306 | 10 | Intron (ENSMUST00000027585.13/72160, intron 2 of 8) |
| Mbnl2 | chr14 120299749 120299981 | 120299786 | 120299807 | 10 | Promoter (2-3kb) |
| 4930553E22Rik | chr16 83272969 83273155 | 83273104 | 83273083 | 10 | Distal Intergenic |
| Myom1 | chr17 71019368 71019599 | 71019456 | 71019435 | 10 | Promoter (<=1kb) |
| Slc8a1 | chr17 81538475 81538689 | 81538593 | 81538614 | 10 | Intron (ENSMUST00000234131.1/20541, intron 1 of 7) |
| Fnbp1 | chr2 31124745 31124964 | 31124910 | 31124889 | 10 | Intron (ENSMUST00000073879.11/14269, intron 1 of 13) |
| Bmpr1b | chr3 141950769 141950962 | 141950804 | 141950783 | 10 | Intron (ENSMUST00000029948.14/12167, intron 1 of 10) |
| Prxl2b | chr4 154908598 154908802 | 154908731 | 154908752 | 10 | Distal Intergenic |
| Ywhah | chr5 33018622 33018835 | 33018726 | 33018705 | 10 | Promoter (<=1kb) |
| Ppargc1a | chr5 51477323 51477496 | 51477346 | 51477367 | 10 | Intron (ENSMUST00000132734.7/19017, intron 7 of 12) |
| Mgll | chr6 88732833 88732995 | 88732916 | 88732895 | 10 | Intron (ENSMUST00000113585.8/23945, intron 2 of 7) |
| 4933427D06Rik | chr6 88941782 88941985 | 88941916 | 88941895 | 10 | Distal Intergenic |
| Sbf2 | chr7 110325827 110326061 | 110325949 | 110325970 | 10 | Intron (ENSMUST00000164759.7/319934, intron 32 of 39) |

**Supplemental Table S2**

List of genomic locations bound by MEF2A in mouse cells and containing called motifs for both MEF2A and TBP separated by less than 100 bp.

#### Supplemental Table S3

| Nearest gene hg38 | genes hg38 | bp between motifs hg38 | bp between motifs mm10 | Annotated location hg38 | TBP peak location hg38 | TBP motif location hg38 | MEF2A peak location hg38 | MEF2A motif location hg38 | Nearest gene mm10 | Neighbouring genes mm10 | MEF2A peak location mm10 | TBP motif location mm10 | MEF2A motif location mm10 | TBP peak E12 mouse liver |
|---|---|---|---|---|---|---|---|---|---|---|---|---|---|---|
| NSMCE2 | WASHC5; TRIB1 | **1** | **1** | Intron | chr8 125219251 125219537 | chr8 125219411 | chr8 125219339 125219503 | chr8 125219421 | Nsmce2 | Washc5; Trib1 | chr15 59480658 59480810 | chr15 59480731 | chr15 59480741 | no |
| LINC02901 | OSTCP1; RSPH3 | **2** | **2** | Promoter (<=1kb) | chr6 158869701 158870000 | chr6 158869779 | chr6 158869674 158869923 | chr6 158869768 | Linc02901 | Ezr; Rsph3b | chr17 6827841 6828061 | chr17 6827948 | chr17 6827935 | yes |
| **KLF6** | LINC02660; LINC0 | **2** | **2** | Promoter (<=1kb) | chr10 3784934 3786509 | chr10 3785521 | chr10 3785420 3785629 | chr10 3785508 | **Klf6** | 1700016G22Rik; Pitrm1 | chr13 5861130 5861331 | chr13 5861219 | chr13 5861232 | yes |
| **KLF7** | CPO; MYOSLID | **2** | **2** | Promoter (<=1kb) | chr2 207165508 207166950 | chr2 207166350 | chr2 207166231 207166455 | chr2 207166337 | **Klf7** | Creb1; Fastkd2 | chr1 64121704 64121931 | chr1 64121824 | chr1 64121811 | yes |
| AP1M1 | **KLF2**; FAM32A | **3** | **3** | Distal Intergenic | chr19 16258401 16258975 | chr19 16258700 | chr19 16258581 16258798 | chr19 16258686 | Ap1m1 | **Klf2**; Fam32a | chr8 72268461 72268713 | chr8 72268578 | chr8 72268564 | yes |
| **KLF2-dt** | **KLF2**; AP1M1 | **3** | **3** | Exon | chr19 16283151 16284100 | chr19 16283782 | chr19 16283695 16283895 | chr19 16283768 | Klf2-DT | Klf2; Ap1m1 | chr8 72295792 72295994 | chr8 72295879 | chr8 72295865 | yes |
| **KLF2** | AP1M1; EPS15L1 | **3** | **3** | Promoter (<=1kb) | chr19 16324451 16325050 | chr19 16324691 | chr19 16324567 16324816 | chr19 16324705 | Klf2 | Ap1m1; Eps15l1 | chr8 72318832 72319054 | chr8 72318929 | chr8 72318943 | yes |
| LOC105376207 | **KLF4** | **3** | **3** | Exon | chr9 107637351 107639043 | chr9 107637748 | chr9 107637665 107637881 | chr9 107637762 | Loc105376207 | **Klf4** | chr4 55651158 55651370 | chr4 55651235 | chr4 55651249 | yes |
| **LRRFIP1** | RAB17; RBM44 | **4** | **4** | Intron | chr2 237616851 237617250 | chr2 237617081 | chr2 237616932 237617151 | chr2 237617096 | Lrrfib1 | Rab17; Rbm44 | chr1 90980737 90980960 | chr1 90980891 | chr1 90980906 | no |
| HSPG2 | RAD2; CELA3B | **7** | **7** | Intron | chr1 21910063 21910246 | chr1 21910207 | chr1 21910087 21910309 | chr1 21910189 | Hspg2 | Cela3b; Ldlrad2 | chr4 137488809 137489024 | chr4 137488900 | chr4 137488918 | no |
| **ERCC1** | POLR1; FOSB | **7** | **7** | Promoter (<=1kb) | chr19 45428801 45429001 | chr19 45428889 | chr19 45428732 45428998 | chr19 45428907 | Ercc1 | Cd3eap; Fosb | chr7 19342130 19342397 | chr7 19342229 | chr7 19342211 | no |

#### Supplemental Table S3

List of genomic locations bound by both MEF2A and TBP and containing MEF2A and TBP motifs separated by less than 10 bp in both mouse and human. Light pink shading indicates known MEF2-TBP motifs within the *KLF2* regulatory elements. Bold numbers indicate evolutionary conserved distances between the TBP and MEF2 motifs in human and mouse.

### Supplemental Table S4

| Genome location of enhancer marks hg38 | Sequence of potential MEF2-TBP motif | Strand | Score | Notes |
|---|---|---|---|---|
| chr19:16324114-16325344 | cctaaatttagccgcggtatataagc | - | 34 | KLF2 promoter |
| chr10:3784928-3785918 | gctatttttag.cagggtatataagg | + | 33.7 | KLF6 promoter |
| chr2:207165693-207166965 | gctatttttag.agggctatataagg | + | 33.4 | KLF7 promoter |
| chr9:107637670-107638171 | gataaaaataaccggcatatttaagg | - | 29.4 | KLF4 enhancer |
| chr19:16283535-16284036 | gttaaaaataactccggtatataaag | + | 28.7 | KLF2 enhancer |
| chr19:16258401-16258902 | gatattttgggagccactatataaac | + | 24.8 | KLF2 enhancer |
| chr2:74893303-74893804 | gctaatttcagccaagctatttaaac | - | 23.6 | |
| chr12:76226253-76226754 | gatatttttagattccatatataagt | + | 21.3 | |
| chr3:152047255-152047756 | gataataataacagcaatagataaga | - | 20 | |
| chr12:9167266-9167767 | gctataaatgaatcaggtatatgagg | - | 18.5 | |
| chr6:25835849-25836350 | gctataaattgccccagtctttaagc | - | 17.1 | |
| chr2:46351593-46352094 | gctaaaaatagccccggtaaacaggc | - | 17 | |
| chr4:139014289-139016106 | gttataaatgaaaaaagtaaataagg | - | 16.4 | |
| chr19:47226196-47226697 | gctataaatagcccgggatatatgagc | + | 16.1 | |
| chr1:21974668-21975169 | gctatttttat.cacagtata.aagg | + | 15.6 | |
| chr9:21802180-21803012 | gataataatggagaccatttatagag | - | 15.5 | |
| chr5:32193314-32194202 | gatattttgaaacagtaaatttaagg | - | 15.4 | |
| chr2:46118813-46119587 | gataaaattagcagagctaagtaggc | + | 15.1 | |
| chr4:10096175-10096676 | gctattttggg.agctgtattgaagg | - | 14.9 | |
| chr19:17135187-17136020 | gatatttttagatgggctttgaaaag | + | 14.8 | |
| chr4:109548377-109549216 | cctaaaatttg.gggactatattaag | - | 14.8 | |
| chr5:172873038-172873761 | gataaaaatacagacggcatttaaac | - | 14.4 | |
| chr1:158037579-158038475 | gtcatttattgaacacgtatataagc | - | 14.2 | |
| chr11:109033757-10903425 | gttattcatagccaaagtatttcaac | - | 13.8 | |
| chr16:74554183-74554684 | cttacatttagccagaatatattgag | - | 13.7 | |
| chr8:26257823-26258618 | gatatatttag.tatactgtaaaagc | + | 13.7 | |
| chr6:118388374-118388875 | gctgtttttagcatgactttataaaa | + | 13.6 | |
| chr6:48826696-48827663 | gctaaatttac.cctgttatttaaga | + | 13.6 | |
| chr3:23740942-23741443 | ggtaagaaggg.caccatatttaagg | - | 13.6 | |
| chr3:170744985-170745486 | catttatttaa.caaagtatttaaaa | - | 13.5 | |
| chr16:12801491-12801992 | gatgattttaaaaagggtttaaaaag | + | 13.5 | |
| chr16:30052712-30053488 | aataaatataaataaaatataaaaag | + | 13.4 | |
| chr14:89137248-89137749 | g.taaaattaaaagcagcatataaac | + | 13.3 | |
| chr6:106546732-106547638 | catataa.tagcagcactttataaag | - | 13.3 | |
| chr1:25429860-25431242 | cctatttataa.gcctctatttaaat | + | 13.2 | |
| chr5:150021800-150022735 | gataaaacaaaagatagtatataagg | + | 13.2 | |
| chrY:3003635-3004136 | gaaataattgg.ctgtgtatttaagg | + | 13.2 | |
| chrY:2608882-2609878 | actattgagagcttcggtatttaaag | + | 13.1 | |
| chr1:25429860-25431242 | gttatatttatagggcctatttataa | + | 13.1 | |
| chr17:29140327-29140828 | gctatttttag.gagcc.atatcagc | - | 12.8 | |
| chr9:109098276-109098777 | gctaaaaatag.gcacataaacaagg | + | 12.8 | |
| chr5:144189559-144190550 | attaaaatgagatgaccaatataagg | - | 12.5 | |
| chr6:2943060-2943874 | gttatttctaaaaagggtattttaaa | + | 12.5 | |
| chr1:101170400-101170901 | tttatttttaaactgaaaatataaac | + | 12.5 | |
| chr2:224417353-224417854 | gatattatgat.taggctatttacag | - | 12.5 | |
| chr6:3796584-3797085 | gattaaatgaaataatgtatataaag | - | 12.4 | |
| chr4:41427212-41427713 | gataaatttta.tggggtacttaaaa | - | 12.4 | |
| chr2:218229053-218229554 | gttataaagaa..gagatatttaaag | - | 12.2 | |
| chr18:23279742-23280459 | gttatatagag.agacatatatatgt | + | 12.2 | |
| chr5:103119681-103120677 | cctaaattcaaacgacagatttaagc | + | 12.2 | |
| chr2:131104881-131105849 | cctatttttagagagcggtatttagaa | + | 12.1 | |
| chr6:110527898-110528399 | aataaaattaattcaagtatataaaa | - | 12.1 | |
| chr5:122727000-122727501 | gctatcttggaagcaagtataaaaac | + | 12 | |
| chr1:91404492-91404993 | gatatattcaaaaccagt.tataaac | - | 12 | |
| chr3:149575923-149576832 | aatatatttaacctgcctatatatgt | + | 12 | |
| chr14:68682584-68683411 | gaaaaaaataacaggcatattttggg | - | 11.9 | |
| chr21:25952104-25952605 | ggtatatttag.aatgatatatatgt | + | 11.9 | |
| chr9:71646978-71647818 | gtta.ttttagctaacatatatgagg | - | 11.9 | |
| chr6:11655440-11656321 | caaaattttaaccagaccataaaagc | - | 11.9 | |
| chr15:74460754-74461731 | gattaagtgagacatggtatataaaa | - | 11.9 | |
| chr10:32961742-32962474 | gaaaaaatggg.gcccctatatatgg | - | 11.9 | |
| chr4:101419510-101420011 | gctgagaagga.cagagtatataaag | + | 11.8 | |
| chr7:134432539-134433040 | gattaaatgagacagtgtatgtaaag | + | 11.8 | |
| chr6:129709805-129710683 | gttaaaattagccatagta.aaaaac | + | 11.8 | |
| chr1:203673868-203674369 | gttatttttag.aaacgaatttcagc | - | 11.8 | |
| chr4:89753392-89753893 | gataaatgttgctccgaaatttaagg | + | 11.7 | |
| chr3:7382176-7383226 | caactaattaaacaagatatataaac | - | 11.7 | |
| chr7:22219895-22220686 | cttatttttag.caaaatatttcacc | - | 11.7 | |
| chr17:47305272-47305773 | gctcattagcaccctcatatataaac | + | 11.6 | |
| chr17:2179335-2180127 | gctacaattag.gaagatatatagca | + | 11.6 | |
| chr15:87609586-87610879 | gataaatataa.atgcttatttcaag | + | 11.6 | |
| chr2:20568855-20569716 | gtcaaatttggccaccctatgtagga | - | 11.6 | |
| chr10:44981202-44981703 | ccta.ttttagattccttatataagg | - | 11.6 | |
| chr12:10716444-10717846 | catattaataa.cagaataaataaca | + | 11.5 | |
| chr2:75502770-75503903 | ccaaaatagaa.aacgatattaaagg | + | 11.4 | |
| chr15:29800466-29800967 | gttaatattagagacactctgaaagg | + | 11.4 | |
| chr6:158869387-158869888 | gaccaaaatag.cgccatataaaagc | + | 11.4 | |
| chr12:79420241-79420742 | gttaaaattgg.ctgaacatttaaaa | - | 11.3 | |
| chr17:29140327-29140828 | cctaaaaatagcagaggaatttgaga | + | 11.3 | |
| chr4:109576661-109577162 | gataattagag.ggaagtttatacac | + | 11.3 | |
| chr19:51571114-51571615 | gctaattttag.ttggataattagcc | + | 11.2 | |
| chr3:14651135-14652057 | gttcttaagagctgaaatatgtaaag | + | 11.1 | |
| chr8:69492459-69492960 | gataataatacgcgggcttatataacc | + | 11.1 | |
| chr3:29324038-29324983 | gttaaaaataaaaacaaaataaaaaa | - | 11.1 | |
| chr2:64644091-64645355 | cttatttacagacagggtaagtaagg | + | 11 | |
| chr15:47665874-47666622 | gaaatttttag.ccagctgttaaagg | + | 11 | |
| chr6:2634157-2635042 | gctttttataaacaaaatattgaaac | + | 11 | |
| chr18:3247212-3247713 | cctagaatatag.agcaatatataatg | - | 11 | |
| chr12:70507213-70508033 | gctaaaactgg.atgaatatagaaag | + | 11 | |
| chr11:33161107-33162011 | cctaaatataa.gaggc.atttaacg | - | 11 | |
| chr5:68248682-68249183 | cataaatattaccaaggaatttaaga | + | 10.9 | |
| chr3:29695456-29695957 | cttaattttag.ccaaaaatgtaaac | + | 10.9 | |
| chr2:36663886-36664387 | gctaataa.agccagggaatatagga | - | 10.9 | |
| chr18:6624573-6625074 | tatagttttggccaagagatataaag | + | 10.8 | |
| chr13:20814308-20814809 | tttaataatag.aaaggcatataaga | - | 10.8 | |
| chr11:72483511-72484371 | gatatttttaa.ccccatattatatg | + | 10.8 | |
| chr6:30769519-30770020 | gctagttttaa.caaaata.ataaag | - | 10.8 | |
| chr13:76552150-76552886 | catatatctaaaacatatatataaaa | + | 10.7 | |
| chr10:17023498-17023999 | gatatatttaa.tggccaatttgaaa | - | 10.7 | |
| chr22:49632322-49633120 | cttaaaattaa.gtaaatacataaac | - | 10.7 | |

| Genome location of enhancer marks hg38 | Sequence of potential MEF2-TBP motif | Strand | Score | Notes |
|---|---|---|---|---|
| chr9:14221402-14222184 | gctaataataa.tgcac.atataaat | - | 10.6 | |
| chr9:89477982-89478483 | gatatatttcagcagggctttatcagc | - | 10.6 | |
| chr1:116509024-116509897 | gatagacattaaccagggtaaataagg | - | 10.6 | |
| chr1:113658997-113659498 | cataaaattggccgagtcatagaagg | + | 10.6 | |
| chr6:21846758-21847752 | cataaatttgaagggagaatatatga | + | 10.5 | |
| chr19:58347241-58348102 | tttaactttaa.aagtgtatttaagg | + | 10.5 | |
| chr16:71894910-71895976 | gctaactagag.accgctaaatcagg | - | 10.5 | |
| chr10:95252782-95253283 | cctaaatttgg.caaaagacataaag | + | 10.4 | |
| chr1:165674701-165675202 | gataaattgtg.atacatatttaaaa | + | 10.4 | |
| chr7:93231868-93232732 | cttatttttaa.acgaatatttctgg | - | 10.4 | |
| chr11:59668724-59669690 | cttaaaaatac.caccctagttaaac | - | 10.4 | |
| chr4:103197910-103198887 | gctaaacagga.gaggggatttaaag | + | 10.3 | |
| chr17:17416284-17416785 | ggtaatttgaccccccctttttaaaa | + | 10.3 | |
| chr18:57780586-57781087 | ggtaatattgtacacgatatgtaaaa | + | 10.3 | |
| chr1:180502373-180502874 | gctaatttcagcgaaacgattaaaag | - | 10.3 | |
| chr20:56468087-56468961 | ctcaaaattaacaacagtattaagag | + | 10.3 | |
| chr9:5499641-5500592 | cctataaattgccccagcctataagc | - | 10.3 | |
| chr11:60913762-60914263 | gggaaaaagaaacgcaatagataaag | + | 10.3 | |
| chr6:22827717-22828792 | gaattttttaa.aggcctaaataagc | + | 10.2 | |
| chr11:110795239-11079574 | gattaaaatga.aaaagtatttaggc | - | 10.2 | |
| chr5:43514742-43515597 | cttatttttaaataaggtaaaaaacg | + | 10.2 | |
| chr19:4246590-4247433 | gctattattggctgcagcatacacag | - | 10.2 | |
| chr8:127206307-127206808 | caaaaaatgagcaggcatttaaaagg | + | 10.2 | |
| chr10:68600206-68600707 | ggtattattat.catggtgtttaaag | - | 10.2 | |
| chr14:100221524-10022202 | gatctttgcagcggagatatttaagc | + | 10.2 | |
| chr13:31161562-31162820 | cttatttttaaaaggtctgatttaagc | + | 10.1 | |
| chr7:106216240-106217432 | gctaacataaactccaaaatataagc | + | 10.1 | |
| chr22:42079164-42080412 | gctaattttgacaggtgtgtagaaaa | + | 10.1 | |
| chr14:58752741-58753242 | aataaatttaa.ggagatat.taaac | + | 10.1 | |
| chr6:18343591-18344271 | tctattatggaaatggatatgtaaag | + | 9.97 | |
| chr1:39132224-39133324 | gctacaggtgaccccaatatttaaag | - | 9.95 | |
| chr4:145097733-145098662 | gatagaaataaagcccatctttaggg | + | 9.94 | |
| chr19:12956938-12957439 | gttatacttgg.aaacgtataaaacc | - | 9.93 | |
| chr14:35414154-35414899 | gatataatgaaaatgcatattagaag | - | 9.91 | |
| chr7:140656503-140657004 | gaaataattgg.ctgtgtatttaggg | + | 9.9 | |
| chr4:77921807-77922308 | gatgtatataaccccattatctagag | - | 9.9 | |
| chr21:25962996-25963780 | ccaaatttgagaaatcgtataaaaga | - | 9.88 | |
| chr17:68312370-68313226 | ggaaatattagtcccattatttaagc | + | 9.86 | |
| chr1:192104955-192105456 | gataaattgac.caacacatataaaa | + | 9.85 | |
| chr1:192559614-192560115 | gttaaaattag.tccagttaataacc | - | 9.84 | |
| chrX:77442489-77443420 | gctattattat.cccgattttaaca | - | 9.82 | |
| chr1:243287415-243288263 | gctaaatataatgcaggtattgaaat | + | 9.8 | |
| chr3:38833879-38834710 | tataaaattagactccat.tataaac | - | 9.8 | |
| chr7:108488987-108489488 | catatttatatgccaactatagaaag | - | 9.8 | |
| chr11:122196745-12219762 | gcttaattgggaaaacagatataaac | - | 9.8 | |
| chr14:24224319-24225073 | catatat.tagctggaatatttgaac | - | 9.78 | |
| chr6:168007292-168007793 | gctattttcggccagcttatacagac | - | 9.77 | |
| chr10:63414596-63415097 | gttatttttagcaaagaaagatgaag | - | 9.76 | |
| chr6:129749546-129750350 | gctagataaag.tggactttttaagg | - | 9.73 | |
| chr1:205121812-205122726 | cttaaa.ttgt.cgcgctatttaaag | - | 9.72 | |
| chr19:45177894-45179108 | gctaaattgag.cgcgatgattcagg | - | 9.71 | |
| chr1:12434786-12435965 | caaaatattag.taccatttttaaag | - | 9.7 | |
| chr5:80247387-80247888 | cctgaatttagctacactacacaaag | + | 9.63 | |
| chr1:44731076-44731577 | ga.aaaagtaa.cgaactatttaagg | - | 9.62 | |
| chr21:34553380-34553881 | cataaaaagtggaacaatatataaaa | + | 9.59 | |
| chr12:115353816-11535459 | catatatataa.ttatgtatataaat | + | 9.57 | |
| chr1:50967824-50969176 | attaaatttaa.agcaaaatttaaaa | - | 9.57 | |
| chr4:159002700-159003562 | tataaacttggcagggagatataaaa | - | 9.56 | |
| chr9:21591430-21592262 | gacatattttgcgggcagatataggc | - | 9.54 | |
| chr2:27081266-27081767 | cataattatagtaaggctatctcggg | - | 9.52 | |
| chr1:58898733-58899552 | gaaatattt.g.tcgcatatttaaag | - | 9.51 | |
| chr14:71130405-71131290 | gctaaagttgtcagaggtatttgaac | + | 9.48 | |
| chr12:26730583-26731084 | tctatttagggctatcatatgtaaag | + | 9.47 | |
| chr20:3795635-3796713 | gctgttatttt.tcgaatatataagg | + | 9.43 | |
| chr14:38290385-38291226 | aataatttaaatgtattatttaaaa | - | 9.42 | |
| chr15:62135344-62136285 | gataatcttagaaaagccatttaaat | - | 9.41 | |
| chr1:145996248-145997110 | cttaaaaataa.aataagatttaaac | + | 9.4 | |
| chr5:73370799-73371300 | cctgaagtgagcaaccttatttaagg | + | 9.39 | |
| chr12:14251046-14251866 | ggtaattagcactaggattttaaag | + | 9.38 | |
| chr6:143673538-143674039 | gcttaatataactagaatatgcaaag | + | 9.37 | |
| chr2:57180007-57180508 | gttcaaaatag.tgccctaaaaaaag | - | 9.33 | |
| chr4:184355039-184355844 | cataaactaggatggggaatataaag | + | 9.31 | |
| chr20:40990631-40992027 | gctattaaaaa.caggaaatatgagc | + | 9.29 | |
| chr14:69079588-69080089 | gaaaatattac.agacatatataatg | + | 9.28 | |
| chr13:100100302-10010080 | gctgtgattggctagcctttataaac | + | 9.27 | |
| chr1:25429860-25431242 | tataaatataa.catcctatattacg | - | 9.24 | |
| chr17:59093475-59093976 | gataaaaataccacaaatattcagag | - | 9.24 | |
| chr15:52491254-52491755 | cttatttttta.aagactctataaaa | - | 9.21 | |
| chr14:52025049-52025976 | gatacttttggtggtggtattgaaag | - | 9.17 | |
| chr1:198776396-198777165 | aatattagaag.ggacgtatataaag | - | 9.17 | |
| chr10:114748754-11474925 | cttaattctag.cacgttatttgagc | - | 9.16 | |
| chr10:17445380-17445881 | gatattattaacaaaagcaaatatgg | + | 9.13 | |
| chr17:42920935-42921436 | cctattgttag.tttggtaaataaac | + | 9.1 | |
| chr12:49188580-49189628 | tttatgtttagattaaatatttaaag | + | 9.09 | |
| chr4:24991204-24991705 | tctaaatgtgactggcctatatagtg | - | 9.07 | |
| chr1:174990404-174990905 | gctgaatagag.tggaatatagaacc | + | 9.07 | |
| chr1:60989762-60990263 | cataaaatgaaaaggccta.ataaat | - | 9.06 | |
| chr11:77589202-77589988 | gcgaattttatctggcatatattacc | + | 9.04 | |
| chr2:235026301-235026802 | cctaaattgaactcagttgtttaggg | + | 9.04 | |
| chr2:204452531-204453032 | gctattttagactatgggatttaaga | + | 9.03 | |
| chr21:41247425-41248298 | gttaaagttagaagccattttcaagc | - | 9.02 | |
| chr17:69466973-69467474 | gtaataaaggg.gtctctatataagg | + | 9.02 | |
| chr11:13135922-13136423 | gctcaatatatatggggtatatatcc | - | 9.01 | |
| chr3:80189837-80190716 | cctatatttatctatggttttaaagg | - | 9 | |
| chr4:11905131-11905632 | aagataagtaaccgcaatatataggc | - | 8.99 | |
| chr11:113875000-11387609 | cagattttggaaaaccaaatataaag | - | 8.94 | |
| chr5:149464902-149465403 | gttattattacccacagtttacaggc | - | 8.93 | |
| chr1:170074270-170075199 | gcagttttgagcgtccgtgtataagg | + | 8.93 | |
| chr2:15359754-15360657 | catatatatagcctaggtatgtagta | - | 8.92 | |
| chr3:143443493-143444578 | cagatttttat.ttccgtatttaaac | + | 8.91 | |

**Supplemental Table S4**

List of top 200 regions with enhancer signatures in human ECs that contain sequences similar to MEF2-TBP motifs. Higher score numbers indicate best alignment as assigned by Glam2Scan [98] . Highlighted in orange are the motifs in the KLF regulatory regions.

### Supplemental Table S5

| Genome location of enhancer marks mm10 | Sequence of potential MEF2-TBP motif | Strand | Score | Notes |
|---|---|---|---|---|
| chr8:72318689-72319365 | cctaaatttagccgcggtatataagc | - | 34 | KLF2 promoter |
| chr13:5860993-5861426 | gctatttttag.cagggtatataagg | - | 33.7 | KLF6 promoter |
| chr1:64121539-64122396 | gctatttttag.agggctatataaag | + | 32.6 | KLF7 promoter |
| chr4:55651050-55651702 | gataaaaataaccggcatatttaagg | - | 29.4 | KLF4 enhancer |
| chr8:72295614-72296079 | gttaaaaataaccccagtatataaag | + | 29.3 | KLF2 enhancer |
| chr8:72268191-72268797 | gatattttgggagccgctatttaagc | + | 25 | KLF2 enhancer |
| chr7:16313883-16314116 | gctataaatag.cccgggatatagga | - | 18.7 | |
| chrX:150477992-150478389 | gataaaaagagagggactatagaatg | + | 15.5 | |
| chr3:53863739-53864182 | cacattttagtcaggcaatataagg | + | 14.6 | |
| chr6:145340073-145341009 | ggtatttatag.ctcg.tatttaaag | - | 14.2 | |
| chr11:101551977-10155288 | gttattttgag.gggcagatagaaag | + | 13.6 | |
| chr2:27539857-27540475 | gataatttcagcgcggttatttagaa | + | 13.4 | |
| chr4:134853131-134854213 | gttatatttatagggcctatttataa | - | 13.1 | |
| chr5:105824209-105824314 | gctatttttgacctccttataaaagt | + | 13 | |
| chr18:34859811-34861220 | gccatattgggccactccatataagg | - | 12.4 | |
| chr8:31149747-31150343 | gctaaatagagcggcagcatattagt | - | 12 | |
| chr10:77434175-77434987 | ggtatttttag..ggcatattaaagc | - | 11.9 | |
| chr4:41275085-41275695 | cctatagagaa.cccactattaaagc | + | 11.8 | |
| chr8:71395788-71396840 | gctaaat.gagacgctgtataaaaac | - | 11.5 | |
| chr8:111854053-111854494 | cctatatctagcttcgctattttggg | - | 11 | |
| chr12:52390353-52390618 | gctatgaatagcaatggtgtacaagg | + | 10.9 | |
| chr6:84058752-84059047 | gttatttttag.tggtctaaatatac | + | 10.8 | |
| chr13:5803066-5803804 | gatattttgaa.gaggagatttcaac | - | 10.5 | |
| chr7:84689639-84690126 | gctattattggtcaaggcatttgagg | + | 10.1 | |
| chr4:134853131-134854213 | cctatttataa.gcctctatttaact | - | 9.99 | |
| chr2:161511873-161512127 | gcgaaatataacgctcctaaaaaagg | - | 9.88 | |
| chr2:131186795-131187039 | gctgttatttt.tcgaatatataagg | + | 9.43 | |
| chr6:34610562-34610768 | gctattaatagcttccctagaatagc | + | 9.38 | |
| chr6:37463960-37464429 | gcgaaatataactctcctaaaaaagg | + | 9.34 | |
| chr10:81429768-81430793 | tctatatttag.aagatgatataaaa | + | 8.99 | |
| chr18:68943595-68943966 | gttattaataaaacagctgtaaaaat | + | 8.77 | |
| chr8:94703248-94704207 | aataaattaaa.aaccctaaataaac | - | 8.71 | |
| chr11:62281042-62281772 | cctctatttagttaagcgatttaagg | + | 8.69 | |
| chr15:77820314-77821020 | gctattcatagggagggtacacaaag | + | 8.51 | |
| chr2:158734657-158734932 | gatgaaaataactgcccgatttcagg | + | 8.47 | |
| chr4:126677419-126677684 | tctaaaattcgacttggtagataagg | + | 8.46 | |
| chr15:78405886-78406722 | catacataaag.ggggctatttgaag | + | 8.26 | |
| chr10:68090279-68090933 | attaaaaatag.tctgctatttcaac | + | 8.1 | |
| chr10:111102284-11110264 | gccaaagatgccagagatatttaagc | - | 8.1 | |
| chr19:6118330-6118938 | gttattattag.cattctgtctaagc | + | 8.09 | |
| chr4:43587680-43588129 | gttaaaaataaagacaattatctaaag | - | 8 | |
| chr11:116086908-11608769 | gattatatttacaagactacttaaac | + | 7.74 | |
| chrX:164419507-164419822 | cctgaaaagagacgcagtaactaagc | + | 7.63 | |
| chr10:81429768-81430793 | tatatttagaa.gatgatataaaaaa | + | 7.62 | |
| chr12:40222337-40223617 | gctaatatgag.gcccctgtccaagc | + | 7.47 | |
| chr3:52198261-52198616 | gattttttcag.gtaggtgtataagg | + | 7.45 | |
| chr4:134853131-134854213 | tataaatataa.catcctatattatg | + | 7.44 | |
| chr18:23954573-23954911 | cctatttttagcttccgggttaaagc | - | 7.39 | |
| chr11:69579255-69580320 | gatactatgaa.aagcctttctaaag | + | 7.28 | |
| chr8:4677401-4678693 | tatatattgagaaaagaaaaataaag | + | 7.28 | |
| chr4:134853131-134854213 | tatatttatagggcctatttataagc | - | 7.22 | |
| chr7:110164176-110164998 | cttcaattgag.cggggt.taaaagg | + | 7.17 | |
| chr3:80014516-80015003 | gggactaatatccaacatatataaag | - | 7.13 | |
| chr1:91062404-91062562 | cttatattcgg.cagactctaaaaac | - | 6.96 | |
| chr5:17923936-17924483 | gataacattga.aaatgtaaataaag | + | 6.87 | |
| chr6:120665677-120666773 | gctaataatagtaataata.ataaaa | + | 6.84 | |
| chr3:8510437-8511095 | actatgtttaaatatcctatttaatg | - | 6.82 | |
| chr9:66511708-66512561 | cctgataatggaccccgtagatgaag | - | 6.81 | |
| chr8:104534376-104535058 | cttgtatatag.tattctatatagag | + | 6.79 | |
| chr10:77434175-77434987 | gctaatttgagattccggttatttaagg | + | 6.63 | |
| chr14:67188586-67189097 | gataaagttaaatgggaaaaaaaaag | + | 6.58 | |
| chr4:134853131-134854213 | gttaaatagag.gcttataaataggc | + | 6.52 | |
| chr3:88409883-88410361 | gttgaaattgaaccgccaaaataaac | + | 6.47 | |
| chr9:46012219-46013025 | ggcaggaatagctacagtatttaagc | - | 6.41 | |
| chr11:54860543-54860885 | gctaagcagag.ctaagtataaaaac | + | 6.33 | |
| chr5:32328232-32328451 | cttatttatag.accaggatcttagc | - | 6.32 | |
| chr12:85473523-85474427 | cctaatatgga.catcctgtgtaaag | - | 6.3 | |
| chrX:99199249-99199900 | cataattgt.gccgggctatttcaga | - | 6.29 | |
| chr2:6463869-6464105 | gctattttgga.aacatgagataaag | - | 6.13 | |
| chr6:71271298-71271955 | cttatattaagactaaatat.taagc | - | 6.1 | |
| chr7:45128726-45129207 | aataaaaatggcccgcctcgatagac | - | 6.01 | |
| chr8:27023565-27023957 | gcaaaaaataaaaaaaataaataaat | - | 5.89 | |
| chr11:78550632-78550957 | tcaaactatagccccgctatatgggc | - | 5.88 | |
| chr7:16312676-16313457 | gctagatttga.ccctgtattaaaat | + | 5.85 | |
| chr5:109556825-109557819 | caaatatagggaatgaatatgtaaac | + | 5.82 | |
| chr6:51469358-51471117 | gagataaacagcggggttctttaagc | - | 5.8 | |
| chr12:21227375-21227828 | gttattattgg.gacactttgtacag | + | 5.71 | |
| chr16:92466092-92466456 | cgtattattgg.tctgctctatagga | - | 5.7 | |
| chr15:76659746-76660285 | ggtaatgtaaa.gagactatctaaac | - | 5.62 | |
| chr6:84058752-84059047 | ggtatatttag.accactaaaaataa | - | 5.62 | |
| chr16:25050305-25050433 | gagaaaaacaaaggcaggatataggc | - | 5.61 | |
| chr2:91236999-91237514 | cctattattaa.tggggaaatgaga | - | 5.61 | |
| chr4:55651050-55651702 | gttatttttat.ccccctacgtcagg | + | 5.6 | |
| chr6:117552423-117552802 | gcta.atttag.ttaagtgtttaaac | + | 5.57 | |
| chr3:88492466-88493249 | cttatttttag.cccagtgtagagtc | - | 5.57 | |
| chr4:117120374-117120891 | gttaaagagga.tgagctacatagag | + | 5.54 | |
| chr16:3908667-3909042 | gttataatgtg.ctccatatatcagt | + | 5.41 | |
| chr17:26596288-26596650 | ggtaaaaatac.agcagcatttaaat | - | 5.4 | |
| chr13:22015830-22016039 | tctatttttaattaacctagaaaagc | - | 5.4 | |
| chr6:108709878-108710131 | gccatttttgg.gtcaatatggaagg | + | 5.35 | |
| chr10:26828703-26829026 | gttaataatga.atagatgtgtaaac | - | 5.34 | |
| chr17:80718582-80718764 | actattttcag.aacaatacaaaaag | + | 5.34 | |
| chr3:96576211-96577188 | cccactttgaa.aggcatatttagga | - | 5.21 | |
| chr7:63911634-63912518 | gctacatctgaccccagactataaac | + | 5.15 | |
| chr8:110997905-110998064 | gctaaaaagag.gaggaaaaaaaaaa | + | 5.15 | |
| chr14:103815761-10381607 | gctattttcaggaataacataaaaag | + | 5.12 | |
| chr15:73511956-73512648 | gctaaatttaaccgaggttcttgcag | + | 5.08 | |
| chr19:43752692-43753484 | gataaaaaaggccaggataaagtaaac | - | 5.06 | |
| chr15:102670839-10267200 | cctaaaatgtagcaggggaaaataaaa | + | 5.04 | |

| Genome location of enhancer marks mm10 | Sequence of potential MEF2-TBP motif | Strand | Score | Notes |
|---|---|---|---|---|
| chr19:53314010-53314446 | cctaaaataaa.aaaaataaaaaaag | - | 5.03 | |
| chr4:9068060-9068258 | tataacatttag.cacactatttatag | - | 5.03 | |
| chr5:114443984-114444349 | gttaatgtttgacccaatctgtaaag | + | 4.96 | |
| chr9:31279997-31281158 | gctaataagaccccgcct.tttatgc | + | 4.91 | |
| chr13:111954392-11195481 | gctatttttga.agacagat.tagag | + | 4.88 | |
| chr2:59438619-59438833 | gacaaaaagagaacccgta.ataagt | + | 4.86 | |
| chr11:103115453-10311631 | gatattttggacaactatttattaaa | - | 4.85 | |
| chr12:80643899-80644208 | cattttttagaaaggct.taaaaaa | - | 4.84 | |
| chr2:38926154-38927036 | gacaaattggaccagag.attaaagg | + | 4.81 | |
| chr19:3575464-3576328 | gttaattatagctggggaat.ta.gc | + | 4.77 | |
| chr13:46929807-46930552 | aagaaaatgag.cgccgtatttacaa | - | 4.7 | |
| chr13:83523815-83524699 | gctcttttaca.cgacctatctaagg | + | 4.67 | |
| chr4:11485631-11486219 | gctatttttag.at.act.tttaaac | + | 4.63 | |
| chr12:80643899-80644208 | taaaaaaatgg.cggca.atataagg | + | 4.56 | |
| chr19:21913887-21914362 | cttattgatagcagccatttgtaatg | + | 4.54 | |
| chr6:86403942-86404527 | tttcttttt.g.cacactatttaaag | - | 4.54 | |
| chr17:26715461-26716634 | gctgcttagag.aggac.atataaac | - | 4.54 | |
| chr7:63887181-63887448 | aataaaagtag.tacggt.tttaaag | + | 4.53 | |
| chr13:112927372-11292788 | cttatttttgccgcccatgtttcagc | - | 4.51 | |
| chr19:21942490-21942905 | cctaatgacat.taaaatatataaag | - | 4.5 | |
| chr5:151065244-151065618 | aataaaatgaacaacaaaaattaaac | - | 4.46 | |
| chr7:68692707-68693001 | cataaaaaagg.gggaacatatcagg | - | 4.46 | |
| chr1:180387418-180387798 | ggaaagaagggcggagctaaataagg | - | 4.44 | |
| chr11:12336132-12336621 | gctaaaaatag.gtaagaatctaatc | + | 4.42 | |
| chr13:23761137-23762440 | ggtcttattgg.cggggtttatgacg | - | 4.41 | |
| chr17:86150989-86151179 | gttaaagttaaagggcgggaataaag | + | 4.35 | |
| chr7:6728416-6728819 | gccaaattgggcggggctctgtgagc | + | 4.34 | |
| chr9:25252133-25253148 | gaaaaaaagaaaaaccatacaaaaga | + | 4.34 | |
| chr13:110538131-11053838 | gataaatttgg.cactgcctttaagt | - | 4.27 | |
| chr12:54695572-54697217 | gctattttcag.cggaaaagaaaacg | - | 4.26 | |
| chr6:134034978-134035585 | gctttaaatggccgggggatttcgaa | + | 4.26 | |
| chr5:119670355-119671009 | ggggaatagagccgagttataaaaag | - | 4.24 | |
| chr15:74967558-74968427 | cctaagtagga.cccagaatctaagc | + | 4.23 | |
| chr5:93266992-93268445 | gctactttggg.cggacttttcaaaa | + | 4.09 | |
| chr10:67285311-67285693 | cttttttttaaaaagcaaatttgagc | + | 4.05 | |
| chr16:3847045-3847396 | gataaatcaaa.cagccaataaaaga | + | 4.04 | |
| chr2:155074153-155074553 | tattcaaataa.gaagatatttaaag | - | 4.03 | |
| chr16:87784547-87784774 | gttaaaaagggaaagagtttatttag | + | 4.03 | |
| chr12:21135836-21136246 | gatatttctagaaatgcaatggaagg | - | 4 | |
| chr8:71395788-71396840 | gatatctttag.agcgggaca.aagg | - | 3.98 | |
| chr4:148140141-148140765 | ccaatctacgaccgggctatagaaac | - | 3.97 | |
| chr16:34952280-34952516 | gctaattttagttgctgtatctgaca | + | 3.94 | |
| chr18:34758761-34759577 | gttataaatctcaggtttataaaagc | + | 3.92 | |
| chr5:77357888-77358691 | cctagtttgaa.cgccctgatatagaa | + | 3.92 | |
| chr12:85374738-85375130 | gctaaacagaaaaggaatgta.aagc | + | 3.89 | |
| chr5:115326944-115327201 | gg.atttttaa.agcgatcttaaagg | - | 3.87 | |
| chr11:87426696-87427010 | aatata.tgagacaaagaatataaac | - | 3.8 | |
| chr14:11811261-11811650 | cataaattcagagggcttgttttagg | + | 3.8 | |
| chr7:100371853-100372332 | gcgcaaattga.tacactatttgagg | - | 3.78 | |
| chr7:100371853-100372332 | gcgcaaattga.tacactatttgagg | - | 3.78 | |
| chr4:151861813-151862676 | cagtattttaa.gccgctagaaaagg | - | 3.76 | |
| chr10:28113394-28114159 | actatatttaa.aaggagaaacaaag | - | 3.76 | |
| chr11:95413780-95414915 | tttaaattatg.cgagggatttaagg | + | 3.76 | |
| chr11:21370563-21371414 | gctatttatgg.gctgg.agaaaagg | - | 3.75 | |
| chr11:6546362-6547289 | cctaaagatggcggcagaattgaaaa | + | 3.73 | |
| chr3:95950850-95951320 | gc.aattagaa.aacggaaaataaag | - | 3.7 | |
| chr17:79895885-79896280 | gttaattttaa.atcggtacaagatg | - | 3.69 | |
| chr17:25067489-25067743 | tctaaaattagcagtgtgatagaaga | + | 3.68 | |
| chr5:113226671-113227009 | cccagtttggaccccagtatttaata | + | 3.67 | |
| chr11:106659493-10666017 | gccatttttttg.ccttgtttataaac | - | 3.66 | |
| chr14:79426181-79426726 | aaaaaaaataaagaggttatgtaaac | + | 3.63 | |
| chr15:102393593-10239403 | gtgggatttaaataccatataaaaag | + | 3.61 | |
| chr1:87900973-87901300 | gctatatttag.cagagcatttccca | + | 3.59 | |
| chr19:10204349-10204690 | gatcataagaa.agaaaaataaaaac | - | 3.58 | |
| chr11:120727242-12072741 | cttgaaaagggcgaggatattaaaat | + | 3.58 | |
| chr18:55095978-55096318 | attaatttgagacagagtttatcatc | + | 3.58 | |
| chrX:8074507-8074831 | gagaaagatgg.cggaaaatttaaaa | - | 3.57 | |
| chr19:55244832-55245287 | gctaattacagacaagggagttgagg | - | 3.56 | |
| chr17:73970444-73970818 | gaaattatgtgcacagggataaaaac | + | 3.54 | |
| chr6:93150460-93150817 | gcaactttgag.ggccaaagataagg | + | 3.54 | |
| chr7:118129398-118129863 | gggaaaggcagctagcctatataagc | - | 3.52 | |
| chr13:99238363-99239043 | gataaagttag.ttgcctggaaaagg | + | 3.51 | |
| chr17:48409553-48410294 | tatataattcagggaggtgtttaagc | - | 3.44 | |
| chr5:86065346-86065929 | gggaacaagagccggagtctttaaaa | - | 3.44 | |
| chr1:64305453-64305898 | gctcttttttt.ccccccatttaaac | + | 3.41 | |
| chr4:105157255-105157481 | cttatatttcacgagcgcctttaaaa | - | 3.39 | |
| chr11:109722812-10972312 | gatatcaggagaaagcctaagtaagc | - | 3.39 | |
| chr6:31518567-31518745 | cagagttagggccagggtattttgag | - | 3.32 | |
| chr6:113483133-113483502 | gctttaaagggcagcggggttaaagg | - | 3.3 | |
| chr5:48372308-48372565 | gctaatgctag.tccaat.tataggg | - | 3.3 | |
| chr16:45349606-45349975 | cttagttttgactgtgatagatacac | + | 3.28 | |
| chr1:130462583-130462771 | gctgttttgagccgcagttaattgag | + | 3.27 | |
| chr7:6728416-6728819 | gctataa.gga.ggggctttataatc | - | 3.24 | |
| chr12:4817231-4817746 | cctaatttgagacgcggactttccgg | + | 3.22 | |
| chr2:85129540-85129823 | taaattaatga.ggaactatatcagc | - | 3.2 | |
| chr7:30094784-30095150 | gttaaaaagag.agcgaaagagagag | + | 3.18 | |
| chr11:101442089-10144246 | gcgtttttgaaccaagctttattagc | - | 3.16 | |
| chr12:72664656-72665161 | gctgtttttaggacgacttttcaggg | + | 3.16 | |
| chr11:20915340-20915518 | gagattttcaaagtgggtgtatcagg | - | 3.13 | |
| chr18:5931967-5932718 | tttattctgggttctgatatttaaag | - | 3.12 | |
| chr19:32182138-32182403 | gacata.atag.agccatatctagag | - | 3.07 | |
| chr8:104534376-104535058 | tctctatatag.aatactatatacaa | - | 3.07 | |
| chr3:130709231-130709765 | ctgataaataatatacatatttaaaa | + | 3.06 | |
| chr4:116074913-116075904 | caaattctttgcaggcgtatatagca | - | 3.04 | |
| chr4:116685487-116685980 | gagaactttt.g.taggctataaaagc | + | 3.01 | |
| chr4:134923250-134923848 | cctcaaaatac.ccggatgtagaagg | + | 3.01 | |
| chr2:144555914-144556419 | gatatattcaa.gaaagtatgtatcc | + | 3 | |
| chr18:34859811-34861220 | ccttatttgggcagcgccttatatgg | + | 2.98 | |
| chr17:24220803-24221080 | cctactttgaa.agcag.agataagc | + | 2.97 | |
| chr18:9393295-9393566 | gttattattaaagggaccagactagg | - | 2.94 | |
| chr2:30441242-30442049 | tccaaatttagaagccagattgaaac | - | 2.9 | |
| chr11:78183565-78183701 | catccaatcag.tgctatatataagc | - | 2.89 | |

**Supplemental Table S5**

List of top 200 regions with enhancer signatures in mouse ECs that contain sequences similar to MEF2-TBP motifs. Higher score numbers indicate best alignment as assigned by Glam2Scan [98] . Highlighted in orange are the motifs in the KLF regulatory regions.
